# Unravelling the Enantioselective Mechanism of Benzylsuccinate Synthase: Insights into Anaerobic Hydrocarbon Degradation Through Multiscale Modelling and Kinetics

**DOI:** 10.1101/2024.10.11.617960

**Authors:** Maciej Szaleniec, Toamsz Borowski, Gabriela Oleksy, Ivana Aleksic, Kai Krämer, Johann Heider

## Abstract

Fumarate-adding enzymes (FAE) are a subset of the glycyl radical enzyme superfamily involved in anaerobic hydrocarbon degradation. Benzylsuccinate synthase (BSS) catalyzes the enantiospecific formation of *R*-benzylsuccinate from toluene and fumarate, initiating anaerobic toluene degradation. In this paper, we present a detailed theoretical study of the reaction mechanism using classical molecular dynamics and multiscale modelling (QM:MM). We describe the potential energy surface of the reaction, confirming the previously postulated mechanism. However, the multiscale character of our model allowed to elucidate the origins of several experimentally observed catalytic phenomena, such as the inversion of the configuration of the benzylic atom upon C-C bond formation, *syn* addition of the abstracted H atom back to the benzylsuccinyl radical, or kinetic isotope effects in the range of 1.7-2.1. The obtained model is supported by microkinetic analysis and was able to explain and quantitatively predict the strict *R*-enantioselectivity of BSS, which is not enforced by the binding orientation of the fumarate, but by dynamic kinetic behaviour of toluene in the active site leading to faster production of the *R*-enantiomer. We were also able to explain the experimentally observed slow H/D exchange in the product during incubation with BSS in D_2_O, confirming the partial reversibility of the reaction. Our study contributes to the elucidation of the catalytic processes catalyzed by BSS and its role in the bioremediation of hydrocarbon pollutants.

## Introduction

Fumarate-adding enzymes (FAE) comprise a subset of the glycyl radical enzyme (GRE) superfamily^1, 2^, which are involved in anaerobic degradation pathways of hydrocarbons or similar substrates and catalyse the addition of chemically inert alkyl carbon atoms to the double bond of a fumarate cosubstrate^3–6^. The first recognized FAE was benzylsuccinate synthase (BSS)^7, 8^, which enantiospecifically forms (*R*)-benzylsuccinate from toluene and fumarate^8–10^ and initiates anaerobic toluene degradation in all known organisms capable of this trait. Very similar initial reactions were also reported for the anaerobic degradation of xylene and cresol isomers^11–13^, as well as for 2-methylnaphthalene^14, 15^, ethylbenzene^16^, and even aliphatic alkanes^17–19^ or cycloalkanes^20^. The discovery of both the characteristic succinate adducts as well as the genes coding for the respective FAE in field studies of contaminated sediments or aquifers indicated that these enzymes are widespread in nature and contribute an important part in bioremediation situations^21–26^.

Like other GRE, the FAE members contain a large subunit of approximately 100 kDa, which forms the active site and contains a conserved glycine close to the C-terminus (Gly829, to be activated to a glycyl radical) and a conserved cysteine at about the middle of the primary sequence (Cys493)^7^. Although the similarity with other GREs outside of the functional canters is limited, the available X-ray structure information on BSS shows very good structural conservation with other members of the GRE^27, 28^. In contrast to the other known GRE, BSS and all related aromatic-activating FAE contain two additional small subunits, which are encoded by separate genes (*bssBC*) in a common operon with the *bssA* gene for the large subunit. The alkane-activating FAE may even be expected to contain three small subunits in addition to the large glycyl-radical containing one, judging from the presence of an additional gene in the respective operons ^29^. These small subunits contain unusual Fe_4_ S_4_ clusters with low redox potential^28, 30^ and appear to be necessary for enzyme activity^3, 31^, but their actual contribution is unknown. Like all GRE, BSS and the other FAE need to be activated to the active glycyl-radical status by an activating enzyme, which is usually encoded in the same operons containing the genes for the subunits (*bssD* in the case of BSS), or in close vicinity to the other *bss* genes (e.g. separated by a gene coding for a transposase in the operons of some alkane degraders^29^). The activating enzymes belong to the large family of S-adenosylmethionine (SAM)-dependent radical enzymes and abstract a hydrogen atom from the specifically recognized conserved Gly residue of the large subunit to convert the enzyme to the catalytically active glycyl radical form. Like all SAM-radical enzymes, these activating enzymes carry an Fe_4_S_4_ cluster in their N-terminal domains, which contains a bound SAM molecule that can be converted to methionine and an adenosyl radical by a one-electron reduction via the Fe_4_S_4_ cluster. The generated adenosyl radical then reacts with the active site glycine of the GRE and abstracts the pro(*S*) hydrogen atom, while being reduced to 5’-deoxyadenosine^32–40^. The glycyl radical in activated GRE is involved in mesomeric interchange with the electrons of the peptide bond and therefore relatively stable. However, exposure to oxygen results in irreversible destruction of the GRE by oxygenolytically cleaving the peptide bond at the site of the glycyl radical^7, 41^. The introduced glycyl radicals in GRE are continuously recycled during the reaction cycles and allow many turnovers without re-activation, instead of requiring the energetically expensive degradation and regeneration of SAM for every reaction round^39, 41, 42^.

The mechanism of BSS has initially been proposed on the basis of generally known GRE enzymes^3, 6, 43^. The glycyl radical represents a stable state of the activated enzyme but is not very reactive. Therefore, it is assumed that the substrates toluene and fumarate bind to the enzyme in this state, followed by closing the active site^31, 44^. The reaction is then initiated by the transfer of a hydrogen atom (HAT) from the conserved Cys493 of the active site to the glycyl radical, generating a reactive thiyl radical, which then abstracts a hydrogen atom from the methyl group of toluene. The generated benzyl radical intermediate attacks the double bond of the bound fumarate, forming a new C-C bond of a benzylsuccinyl product radical. Finally, the product radical re-abstracts the hydrogen atom from the side chain of Cys493, and the cascade continues backwards from the resulting thiyl radical to Gly829. This re-establishes the stable glycyl radical form, which allows the enzyme to open the active site for product release and bind new substrates. The mechanism has been found reasonable by DFT calculations in the gas phase^45^ and more recently by preliminary QM studies based on the X-ray structure of BSS, which yielded somewhat modified values of the transition states, but in

In particular, this version of the reaction mechanism is fully in accordance with the experimentally determined (*R*)-enantiospecificity of the reaction [9], the return of the abstracted hydrogen in a *syn*-addition^46^, and the inversion of the methyl group of toluene during the reaction^47^, and a specific preference of toluene adding to the distal C-atom of the double bond of fumarate to enable the product radical to retrieve the hydrogen from Cys493^44^.

In this study, we present a more detailed QM:MM model of the BSS mechanism. By this advanced computational modelling, we obtained new insights into the geometry of the transition states limited by the added constraints of the whole protein, as well as the potential energy surface of the whole reaction. We were able to identify factors by which the enzyme is imposing the regioselectivity and enantioselectivity of the C-C bond formation as well as propose a mechanism for H/D exchange events observed during the reaction. We also verify the proposed mechanism with the comparison of kinetic isotope effects obtained from theoretical predictions and experiments.

## Methods

### BSS Model Preparation

The initial structure of the α subunit of BSS_T1_ in complex with monoprotonated fumarate and toluene was obtained from the crystal structure (PDB codes: 5BWD, 5BWE)^27^ as described previously by Salii *et al*^31^ and Szaleniec *et al*^48^. The monoprotonation state of the fumarate was selected based on the docking studies presented in^44^ and is consistent with a recently published investigation of the initial phase of the reaction^48^. Furthermore, modelling attempts of the whole pathway with completely deprotonated fumarate turned out to be extremely challenging due to problems with the convergence of such QM:MM models.

### MD simulations

All classical MD simulations were performed for holo and apo BSS models using the AMBER ff03 force field^49^. The calculations were conducted with AMBER 18^50^ according to the previously described protocol^31^. The following MD simulations were conducted for the study: five 62 ns simulations for BSS with radical Gly829 in complex with toluene and monoprotonated fumarate in pro*R* orientation (Fig. S2), three 62 ns simulations for BSS with radical Gly in complex with toluene and monoprotonated fumarate in pro*S* orientation (Fig. S3), four 62 ns simulations of BSS with radical Cys493 in complex with toluene and monoprotonated fumarate in pro*R* orientation (Fig. S4), two 62 ns simulations of radical Cys493 and monoprotonated benzylsuccinate (protonation at carboxyl group close to Arg508) (Fig. S5), as well as two 100 ns simulations for BSS with radical Gly829 in complex with mono-protonated fumarate in pro*R* orientation (Fig. S6A,B), pro*R* orientation with the protonated group rotated for 180° (Fig. S6C,D)and pro*S* bound orientation (Fig. S6E,F). The reagent binding ΔG_b_ values were calculated using MM/PBSA protocol^51^ using the Poission-Boltzman and Generalized Born approach.

### QM:MM

All QM:MM calculations were conducted using the Gaussian16 C.01 program^52^. The QM:MM model obtained from MD simulations was stripped from sodium ions and most of the water molecules, leaving only H_2_O molecules penetrating a 20 Å radius from Cys493. The positions of all residues and water molecules outside a 15 Å radius from Cys493 were frozen in geometry optimization. The fumarate was modelled in the monoprotonated state according to previous docking studies^44^, and the model’s overall charge was -7. Two sizes of high layer (HL) were used for the QM portion of the study: a small one (QM1, Fig. 1a) used for geometry optimization and vibrational analysis or a big one (QM2 Fig. 1b), used for single point correction of the energy. The QM1 comprised of Cys493 and Gly829 residues with adjacent fragments of the main chain, Gln707, fumarate and toluene. The QM2 comprised of QM1 extended by all residues and solvent molecules penetrating a 3 Å radius of Cys493, toluene and fumarate (330 atoms). The charge of both models was -1. All calculations were conducted for a doublet state due to the presence of a single radical.

**Figure 1.**
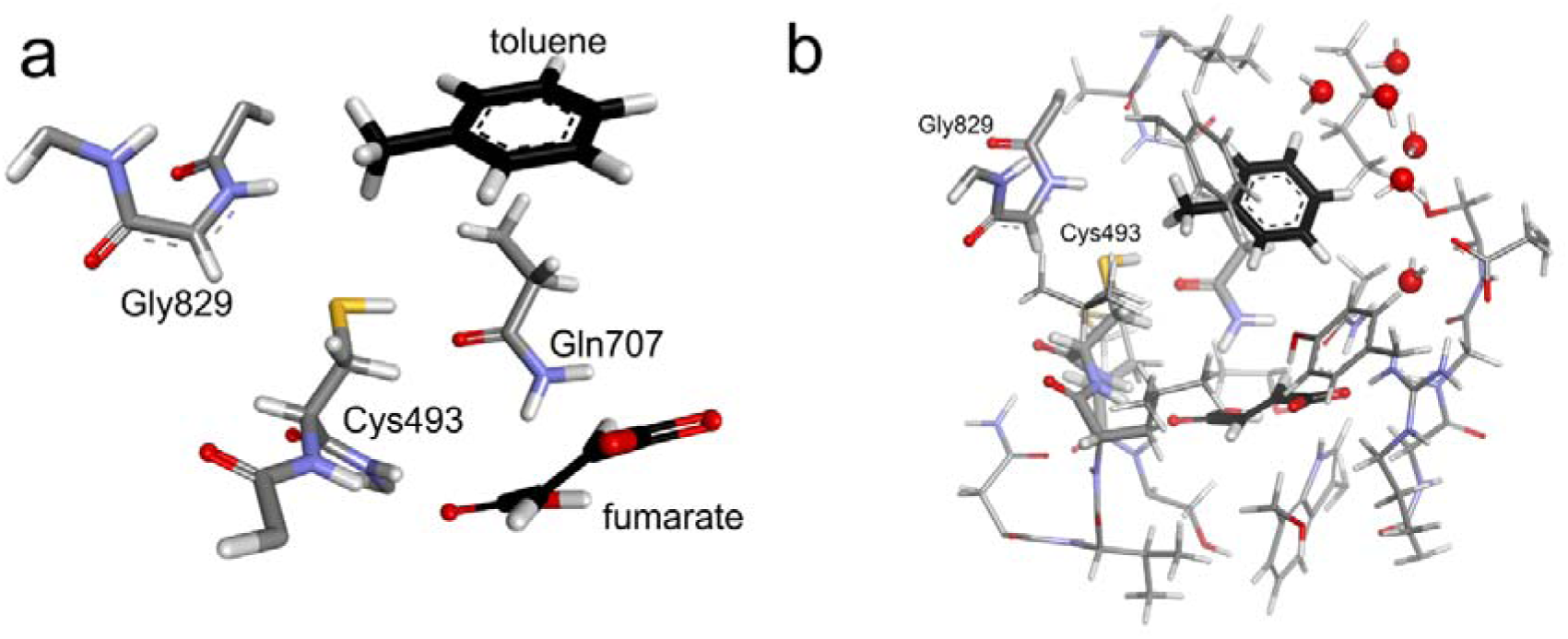
The QM (HL) part of the QM:MM BSS models: a) a small one (QM1) used for geometry optimization and vibrational corrections and b) a big one (QM2) used for single point energy corrections.

The geometry of the QM:MM models was optimized at the B3LYP/6-31g(d,p):AMBER level of theory, using an electronic embedding approach^53^ which was followed by vibrational analysis introducing vibrational corrections for stationary points. The transition states were localized using relaxed scans along the reaction coordinate (e.g. d(Cys-S-H^…^C^rad^-Gly)) followed by TS optimization using the Berny algorithm. Each stationary state preceding or following a particular TS was optimized individually, with initial geometries derived from an intrinsic reaction coordinate (IRC) scan. The energy of the final stationary points was corrected with single point calculations using QM2 at B3LYP/6-311g+(2d,2p):AMBER level of theory with electronic embedding and Grimme D3 corrections for dispersed interactions^54^. The electronic energies were corrected with zero-point energies calculated at default conditions (1 atm., 273 K, no scaling factor). The inevitable discontinuities in the potential energy surface along the multistep reaction which originated from changes in residue conformation(s) due to reagent movement during the reaction scans were corrected by applying reference calculations (i.e. calculation of the particular HL geometry in the new conformation of the MM part, see Tables S1-S3).

For the estimation of the intrinsic kinetic isotope effects, we used d_8_-toluene. The electronic energies were corrected with thermal energy corrections calculated at the B3LYP/6-31g(d,p):AMBER level of theory at 303 K, 1 atm. and using 0.9806 scaling factor according to B3LYP/6-31g(d) correction calculated by Scott and Radom^55^.

### Kinetic Rate Estimations

All kinetic constants were calculated according to a standard equation from transition state theory:

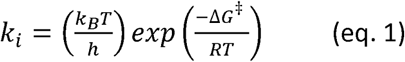

where k_B_ is the Boltzmann constant, h is the Planck constant, R is the gas constant, T is 303 K, and the transmission coefficient is assumed to be a unity.

However, to avoid potential influence of biased entropy prediction (due to partially geometrical constrains of the QM:MM model) on values of kinetic isotope effect, for its prediction we used thermal energy corrections instead of Gibbs free energy^56^.

To account for the tunnelling effect that may be involved in the H atom transfer the obtained rates were corrected by Wigner’s tunnelling corrections Γ(T)^57^ according to eq. 2-3:

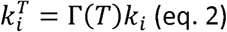

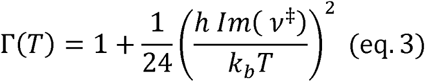

where ν^‡^ is the imaginary vibration frequency [Hz] associated with the transition state.

The calculated kinetic constants were used to estimate the reaction rate based on the equation derived according to the King-Altman algorithm^58^ (see SI methods section).

### BSS production in *Aromatoleum sp*

Cell-free extract from toluene-grown *Aromatoleum sp.* was prepared as described in Szaleniec et. al^48^.

### Kinetic isotope effect

The kinetic isotope effect was measured using two methods i.e., a direct method where kinetic assays are conducted independently with labelled and unlabelled substrate or with a competitive method, where the reaction is carried out with the mixture of both isotopologues of the substrate.

The experimental values were compared with the kinetic isotope effects predicted based on microkinetic constants calculated for the reaction pathway with toluene and d_8_-toluene.

### Direct kinetic isotopic effect

Direct KIE was measured by conducting separate assays for unlabelled and isotopically labelled d_8_-toluene. The reaction mixture comprised 0.2 mL of cell-free extract (C 39 mg/ml) buffered with 0.73 mL of 20 mM TEA/HCl pH 7.8, 50 μL of 100 mM aqueous solution of sodium fumarate (final concentration of 5 mM) and 3 mM of toluene (8.2 μL of a 364.5 mM stock solution in isopropanol) or d_8_-toluene (8.4 μL of a 358.2 mM stock solution in isopropanol). The reactions were conducted anaerobically at 30°C. Samples of 150 μL were collected at 0, 5, 10, 15 and 20 min, and the reaction was stopped by the addition of acetonitrile in a 1:1 (v/v) ratio. Then samples were centrifuged (8000 x g, 20 min) to remove the precipitated protein and the supernatants were analysed by LC-DAD. All experiments were carried out in triplicates.

The catalytic activities of BSS with toluene and d_8_-toluene were determined by linear regression of product concentration over time for the first 20 min of the reaction using OriginPro 2021. The KIE was calculated as a ratio of averaged enzyme activities recorded for toluene to the activity recorded with d_8_-toluene.

### Competitive kinetic isotopic effect

The reaction mixture, comprised of 0.2 mL of cell-free extract (39.8 mg/ml), 0.73 mL 20 mM TEA/HCl buffer pH 7.8, and 50 μL of 100 mM sodium fumarate (final concentration of 5 mM), was assembled under anaerobic conditions and incubated at 30 ⁰C at thermoblock for 2 min before reaction initiation. The reactions were started by the addition of a 3 mM toluene/d_8_-toluene mix dissolved in isopropanol, resulting in concentrations of 1.5 mM toluene and d_8_-toluene, respectively. The isopropanol content in each reaction mix was adjusted to 2%. Samples of 150 μL were collected at 0, 5, 10, 15 and 20 min, and mixed in a 1:1 ratio with acetonitrile to stop the reaction progress. Then they were further processes as described above. All experiments were carried out in triplicates.

The signal ratio of lighter and heavier products (207→163 m/z vs 215→171 m/z) from MS/MS was combined with the quantitative analysis from DAD. The competitive KIE on 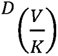 was estimated by following the changes of benzylsuccinate and d_8_-benzylsuccinate molar fractions over the course of the reaction up to 30% of conversion of the lighter isotopologue, according to the formula:

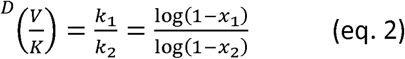

Molar fractions of heavier (x_2_) and lighter (x_1_) products were plotted in OriginPro 2021 (Fig. XB). 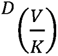 was obtained by non-linear regression to the x (x), following the equation^59^:

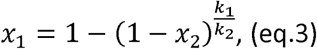

### UHPLC-DAD-MS/MS

Samples were analysed with UHPLC (Agilent 1260) equipped with DAD and 6460 Triple Quad MS detectors according to procedures described in ^48^. In short, the separations of direct KIE samples were conducted on ZORBAX 300 SB-C18 column (RRHD, 2.1×50 mm, 1.8 μm, Agilent) at 30 °C, at the flow rate 0.4 ml/min with a gradient method in H_2_O/ACN/0.01% HCOOH mobile phase (see SI) using 2 μl injection volume and 210 nm as a detection wavelength.

The samples from competitive KIE were separated using the same chromatographic program and the MS signal was detected in Multiple Reaction Monitoring mode (MRM), mode using a jet-stream ESI ion source in negative polarity mode following the transition of the parent [M-H]^−^ ions of 207 m/z and 215 m/z respectively for benzylsuccinate and d_8_-benzylsuccinate and fragmentation [M-H_2_O-H]^−^ quantifier ions of 189 or 197 m/z as well as [M-CO_2_-H]^−^ qualifier ions (respectively 163 or 171 m/z). Additionally, the signal of d_7_-benzylsuccinate (214 m/z), i.e. the product of H/D exchange occurring in H_2_O during the reaction, was also analysed (fragmentation ions of 188 and 196 m/z) and added to the abundance of d_8_-benzylsuccinate (see Table S1-S3 for method parameters). Additionally, the 135→91 m/z transition of phenylacetic acid as an internal standard was monitored and used for the correction of sample preparation errors. The data from the MS experiment were used to establish a ratio of labelled and unlabelled products while the total concentration of the product was established by analysis of the undiluted samples with the DAD detector (Fig. S1).

## Results

### Step 1 – activation of Cys493

The first step of the reaction (Fig. 3) involves the hydrogen atom transfer (HAT) from Cys493 onto radical Gly829. As indicated by MD simulation, the conformation of the Cys493 residue is predominantly (96% of simulation time) oriented toward Gln707 with an H-bond formed between the SH group of Cys493 and the carbonyl group of Gln707 (Fig. 2a and Fig. S7). Therefore, to transfer the H atom, Cys493 has to first break the H-bond stabilizing its position, turn the SH group to face towards the glycyl radical and slightly rotate around the Cα-C*β* bond (Fig. S8). This conformational change elevates the energy of the system by 3.63 kJ/mol and this geometry (ES^rot^) is used as a reference point for the whole reaction.

**Figure 2.**
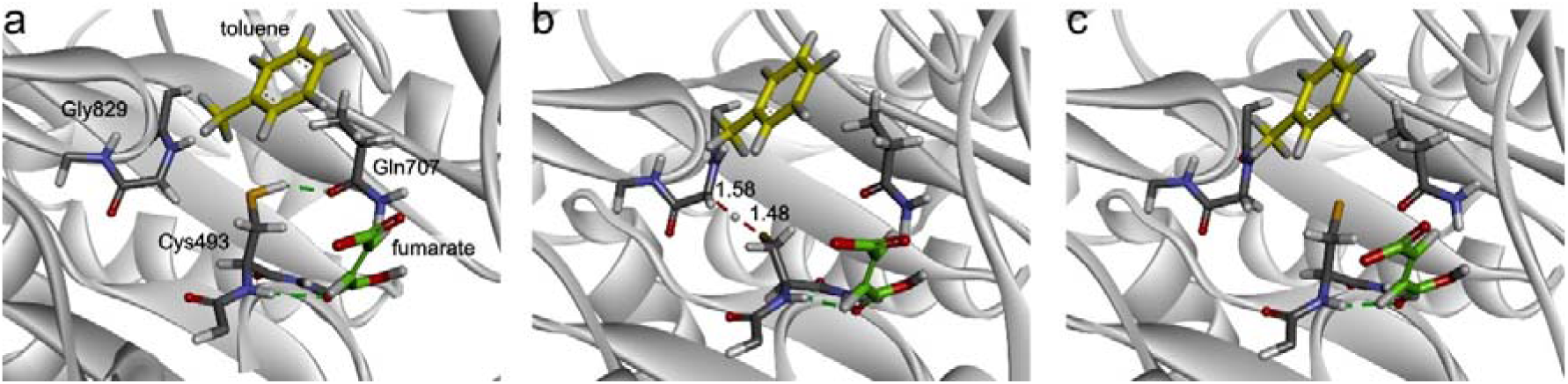
Geometry of BSS during Cys activation: a) E:S complex, b) TS1, c) intermediate I1a. Fumarate in green, toluene in yellow.

**Figure 3.**
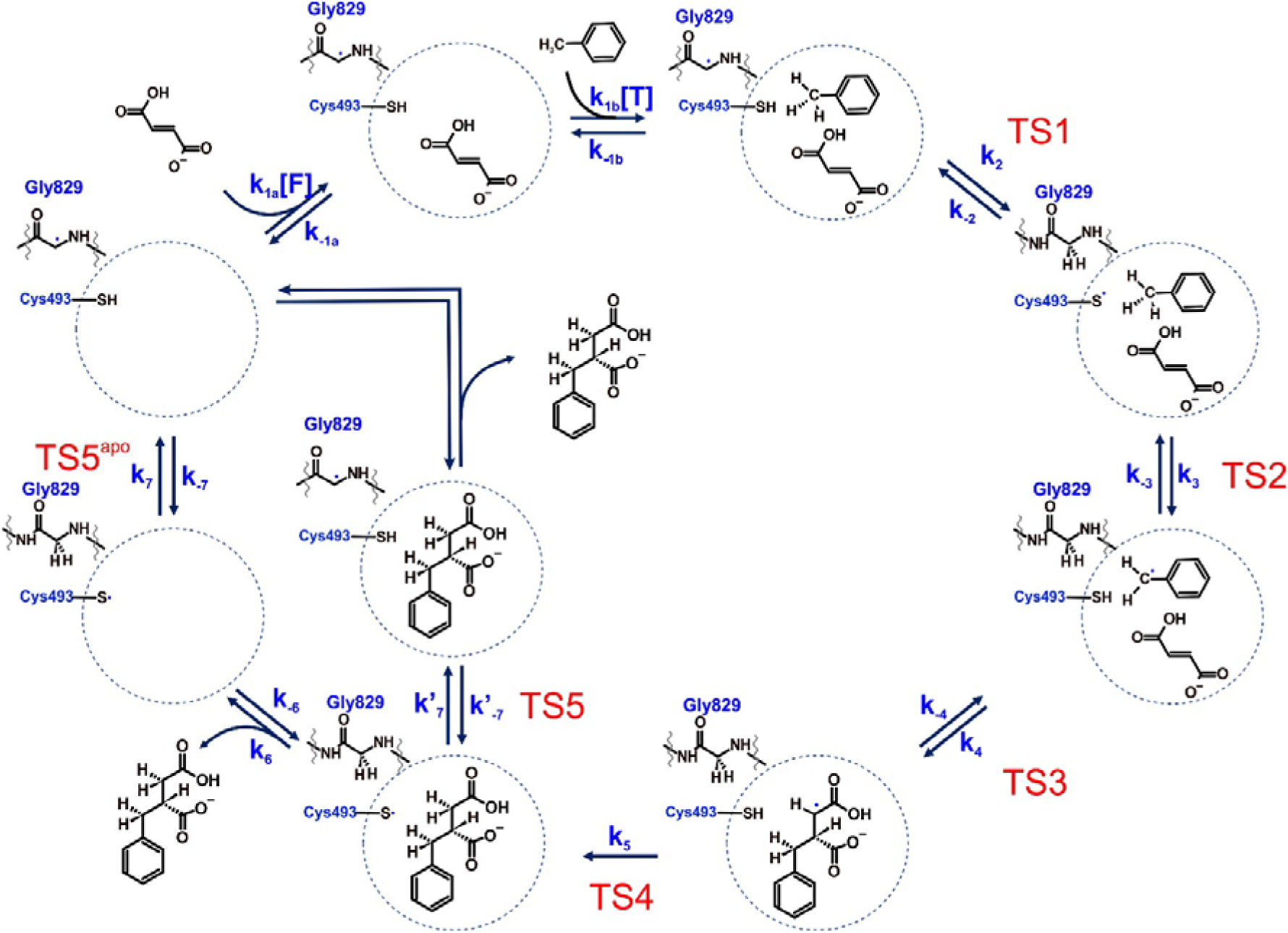
The mechanism proposed for *R*-benzylsuccinate formation taking into account current work and a recent report on HAT between glycyl radical and Cys493^48^.

We have recently devoted much attention to this process^48^ showing that Cys493 can transfer the H atom from either the *re* or *si* face of the glycyl radical, and while the former is associated with a lower energy barrier (55.9 kJ/mol), the latter is significantly slower but still expected to occur occasionally in the same time frame (barrier 71.0 kJ/mol). In the transition state (Fig. 2b), the bond lengths are 1.48 Å for S-H and 1.58 Å for C-H, while the C-H-S angle is 167°. The radical spin density of the transition state is divided between the Cα of Gly829 and the S atom of Cys493 (0.66 vs 0.31, respectively).

The HAT is completed with the formation of Gly829 in the non-radical state and the formation of the (almost) fully radical thiyl group at Cys493 (0.875 spin located at the S atom), which is calculated to be more stable than ES^rot^ (by 31.84 kJ/mol, Fig. 8) and positions its radical residue in between the glycine loop and Gln707 with a dihedral angle C-Cα-Cβ-Sγ of 52.5°. Because the removal of an H atom from the sulfhydryl group results in higher mobility of the resulting radical cysteine, allowing different conformations of the thiyl radical, the changed values of this angle were convenient markers for assessing the next reactions continuing the reaction cycle.

Although the radical placed between Gln707 and Gly829 (at an angle of 52.5°) remained the most probable conformation (69% of simulation time in 59°±29°), the chance of attaining alternative conformations with dihedral angles C-Cα-Cβ-Sγ of 287°±21° and 169°±29°, which both point towards the active site, became higher for the thiyl radical form (the former at 28% of simulation time, the latter at 2 %; see Fig. S7B). To gain more insight into the energy barriers associated with the conformation changes of the radical Cys, we have performed a QM:MM-scan (with a 15° step) of the effects of changing the C-Cα-Cβ-Sγ dihedral angles of Cys493 on the respective energy barriers. First, we have discovered that clockwise and anticlockwise rotation of the Cys residue does not yield the same barriers, most probably due to the conformational flexibility of the active site and shifts of residues triggered by the movement of Cys. Nevertheless, we were able to identify the same energy minima as observed in MD simulations at similar C-Cα-Cβ-Sγ dihedral angles, namely 52.5°, 190° and 280°. The radical is oriented towards the active site in the two conformations with 190° or 280° dihedral angles, whereas the radical is directed towards the opposite side in the conformation at 52.5°. Both scans show that the highest rotation barrier is associated with the transition between the minima at 52.5° and 190°, with transition energies in the range of 76 to 86 kJ/mol (depending on the direction of rotation). The rotation barrier between the minima at 52.5° and 280° turned out to be much smaller, approximately 33 kJ/mol. Finally, the rotation barrier between the two minima presenting the thiyl radical towards the active site (at 280° and 190°) is calculated to be around 24 kJ/mol for clockwise rotation and only 7 kJ/mol for counterclockwise rotation (see Fig. S9 for the rotational analysis). This analysis shows that the radical-carrying cysteine residue will very rarely rotate in the counterclockwise rotation, mostly due to steric hindrance introduced by the loop carrying Gly829. Instead, it will rotate clockwise, breaking away from the interaction with Gln707 at 52.5° to reach the first local minimum at 287° and finally the shallow second minimum around 150-190° (Fig. S9).

### Step 2 – activation of toluene

The activation of toluene proceeds by the HAT from its methyl group to the thiyl radical (Fig. 3). As was shown above, the radical Cys can exist in three different conformations corresponding to C-Cα-Cβ-S dihedral angles of 59°±29°, 169°°±29°° and 287°° ±21° (Fig. S7), which results in different geometries and energies of the transition states reached from the respective conformations (TS2a-c). To elucidate this issue, we have studied the activation of toluene associated with all three different starting conformations of radical Cys (denoted a, b, and c in the following section). As a result, the C-Cα-Cβ-S dihedral angles of the resulting TS changed somewhat, but stayed close to those of the starting states, yielding values of 83° for TS2a, 295° for TS2b and 179° for TS3c (Fig. 4).

**Figure 4.**
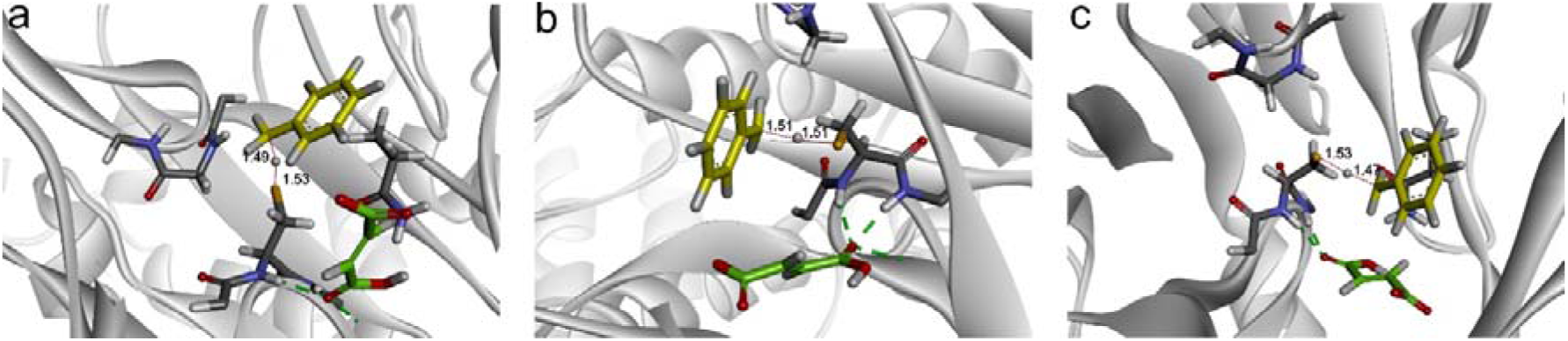
Three variants of toluene activation by radical Cys: a) TS2a with C-Cα-Cβ-S dihedral of 83°, b) TS2b with C-Cα-Cβ-S dihedral of 295°, c) TS3c with C-Cα-Cβ-S dihedral of 179°. Fumarate in green, toluene in yellow, and distances are in Å. Red dashed line mark formed/broken bonds in TS.

As was already obvious in the previous MD simulation, the respective conformations of the radical Cys rotation isomers differ in their relative abundance, with I1a being the most abundant and I1c the rarest. It became clear during the calculations that the active site structure of the model used had to be redefined because the initial position of Tyr197 was in the way of positioning toluene for H-abstraction in pathway c. Although we started with an initial frame of the E:S complex containing Tyr197 bound to the fumarate, the steric clash with toluene and subsequent re-positioning of Tyr197 observed in the TS2c stage prompted us to re-evaluate all values with the alternative conformation of Tyr197. Inspection of the respective geometries showed that TS2c significantly changes the binding mode of fumarate, while toluene pushes Tyr197 out of the active site into a different conformation. Prompted by this observation, we analysed MD simulations, which indicated that Tyr197 probably acts as a gating residue that can also attain three different conformations: the two major populations display C-Cα-Cβ-Cγ dihedral angles with maxima at 68° ±23° and 309° ±19 and a smaller population is represented by a small and wide peak of dihedral angles with a maximum of 128° ±42° (Fig. S10A). Only for the population of the first peak (dihedral angle 68%), Tyr197 is directed toward fumarate. Furthermore, the OH group of Tyr197 comes into the H-bonding distance (below 4 Å) of the fumarate O1 oxygen atom only for approx. 4% of the simulation time (Fig. S10B). The re-positioned Tyr197 in TS2c corresponds to the minor conformer with a 143° dihedral angle (i.e. conformation within a wide population described by a 128° ±42° peak). Under these conditions, the energy of the whole system is elevated by approx. 18.2 kJ/mol, but we consider this step necessary because the reaction cannot proceed further with Tyr197 bound to fumarate. Therefore, the energy-minimized geometries of the respective I1a,b,c states with the alternative conformation of Tyr197 were used for the QM:MM calculations, creating comparable data sets for all three Cys conformations. The energy discontinuity of the reaction profile associated with the conformational rearrangement associated with the shift of Tyr197 was corrected by matching in a factor of -18.2 kJ/mol (i.e. the energy difference between I1a obtained before and after the Tyr197 shift). Therefore, all energies reported for these and later steps in the text are corrected for this factor while the originally calculated energy values are reported in SI (see Table S3)

As expected, I1a turned out to have the lowest energy of -28.2 kJ/mol (corrected to the same value as obtained with the initial conformation of Tyr197), surprisingly followed by I1c with -13.7 kJ/mol and finally I1b with 4.9 kJ/mol. This suggests a reversal of the energetic preferences of I1b and I1c, compared to their calculated energies after step 1 of the reaction. This change in relative energy may be associated with a significantly bigger part of the enzyme treated with the QM method (QM2 instead of QM1) and using Grimme dispersion D3 corrections to describe van der Waals interactions instead of an AMBER forcefield. This explanation is additionally corroborated since we had observed an energy order of these intermediates consistent with their abundances in MD simulations at lower levels of theory (i.e. QM1/AMBER).

Each of the three conformations of radical Cys theoretically allows to activate toluene by the thiyl radical, but via totally different approaches (Fig. S11). The TS2a transition state is characterized by S-H and C-H distances of 1.53 Å and 1.49 Å, respectively, and the three atoms are aligned along a slightly bent line (angle S-H-C of 166°). As the radical Cys is oriented away from the active site and towards Gly829, the toluene has to approach very closely to the surface of the active site cavity, and the only possible orientation enforces positioning the toluene above the fumarate cofactor and the thiyl radical. Because the activated hydrogen is localized between toluene and fumarate, the subsequent C-C bond formation would proceed without inversion of the methyl group configuration. In the TS2a the spin is already mostly localized on the activated substrate (0.54 on the methyl carbon and 0.23 on the aromatic ring) with 0.35 still localized on the Sγ atom of Cys. The energy of TS2a is 49.7 kJ/mol which results in a barrier for toluene activation of 77.9 kJ/mol.

TS2b allows for seemingly more convenient positioning of toluene in the active site as the radical Cys is facing towards the active site, close to Gln707. In this conformation, both distances in TS2 for d(S-H) and d(C-H) equal 1.51 Å, while the S-H-C angle is slightly more stretched, reaching 178°. The spin density on the substrate reaches 0.56 at the methyl carbon atom, 0.16 at the aromatic ring and 0.29 at the Sγ atom of Cys. The barrier of activation turned out to be of the same magnitude as for TS2a (74.4 kJ/mol) which together with an elevated energy of I1b results in the highest absolute energy of TS2b (79.3 kJ/mol).

Finally, the distances in TS2c for d(S-H) and d(C-H) are 1.47 Å and 1.53 Å, respectively, while the S-H-C angle is 167°. The spin density is similar to the previously reported situations in TS2a and TS2b, with 0.5 at the methyl carbon, 0.14 spread over the aromatic ring and 0.26 at the Sγ atom of Cys. In this conformation, the radical Cys is localized directly over the get activated, but its approach is checked by Leu492. As a result, it is positioned at a 109.65° angle with respect to the fumarate plane. Remarkably, only this conformation leads to H abstraction from the face of toluene opposite to that oriented toward fumarate, resulting in inversion of configuration after C-C bond formation. While the energy of I1c is slightly higher than that of I1a (by 15 kJ/mol), the barrier associated with toluene activation in TS2c turns out to be the lowest of all three conformants (47.6 kJ/mol), and the resulting energy difference of TS2c and I1c is only 34.4 kJ/mol. Therefore, the TS2c-type activation is the most kinetically favourable of all three variants, as well as consistent with the experimental observations that the reaction involves inversion of configuration at the methyl group^47^.

In each case, the I2 intermediates exhibit higher energy than the I1 intermediates by 20-26 kJ/mol (see Fig. S11, Table S6 and Fig. 8) and the relative energies exhibit the same order as in case of I1, with I2a at the lowest (-7.7 kJ/mol), I2c at an intermediate (13.2 kJ/mol) and I2b at the highest level (27.7 kJ/mol). As the TS2c barrier was clearly the lowest and the reaction path is favoured by the known involvement of a methyl group inversion, the following steps of the reaction were continued from I2c.

### Step 3 - C-C bond formation

In the I2 intermediate, the benzyl radical derived from toluene is localized directly over the fumarate co-substrate. The C-C bond can principally be formed by the attack of the benzyl radical at any one of the carbon atoms of the double bond of fumarate, which are labelled as “distal” (C2) or “proximal” (C3) to Cys493 in the following, based on their distances (further away or closer, respectively). To determine the regioselectivity of the process, we have studied both pathways, designating attack on the distal C2 atom as variant a and on the proximal C3 atom as variant b. First, we used an MD simulation conducted for the E:S complex to model the probability of the attack of the benzyl radical on either of the fumarate carbon atoms. Although this complex contains the Gly radical and toluene instead of the benzyl radical, these simulations provided a good approximation of the behaviour of the benzyl radical intermediate, especially since Cys493 is in the protonated non-radical state in both cases (which models the behaviour of the Cys residue correctly), whereas the state of Gly829 does not have much influence on the active site.

The distances from the methyl C atom of toluene to each of the C atoms of the fumarate double bond were analysed in the MD trajectory with the histograms of distance distributions presented in Fig. S12. We observed that toluene approaches closer to the distal C atom. Therefore, we estimate that there is a higher probability of the benzyl radical attacking fumarate at the distal C atom (pathway a) compared to an attack at the proximal atom (pathway b).

Similarly to the analysis of the previous step, we observed yet another conformational shift while assessing the QM:MM models of this step, which became apparent during the optimization of the TS3a. This time the change applies to a conformational adjustment of Trp613 which also triggered changes in the conformations of Asn611, Ser495 and Asn615. As Trp613 and Asn615 directly interact with the unreactive face of fumarate, these modifications resulted again in a change of the MM energy of the whole system, amounting to a value of 212 kJ/mol. The discontinuity of the energy profile was corrected by aligning the energies of the I2 state obtained from a scan of internal reaction coordinate (IRC) from TS3a and TS3b with the previously shown energy of the I2c state (13.2 kJ/mol), obtained from TS2c.

In TS3a, the benzyl radical approaches the distal C atom of the double bond of fumarate, forming a new C-C bond. The C-C distance in TS3a is 2.23 Å (Fig. 5A). Due to the steric interaction of the phenyl ring of the radical with some active site residues (Gln707 and Ile617) and some distortion of the fumarate structure, the bond being formed is not directly perpendicular to the fumarate frame, but at a little wider angle of 106°, which is in good agreement with the resulting sp^3^ hybridization of the intermediate. The fumarate is also significantly distorted with the distal atom moving out of the plane and closer to the benzyl radical, which corresponds to the change from sp^2^ to sp^3^ hybridization of the C atom. The spin density is distributed on the benzylic carbon atom (0.58) and the aromatic ring (0.18) as well as the proximal carbon atom of the double bond of fumarate (0.32), while the distal carbon atom exhibits a negative spin value of -0.12.

**Figure 5.**
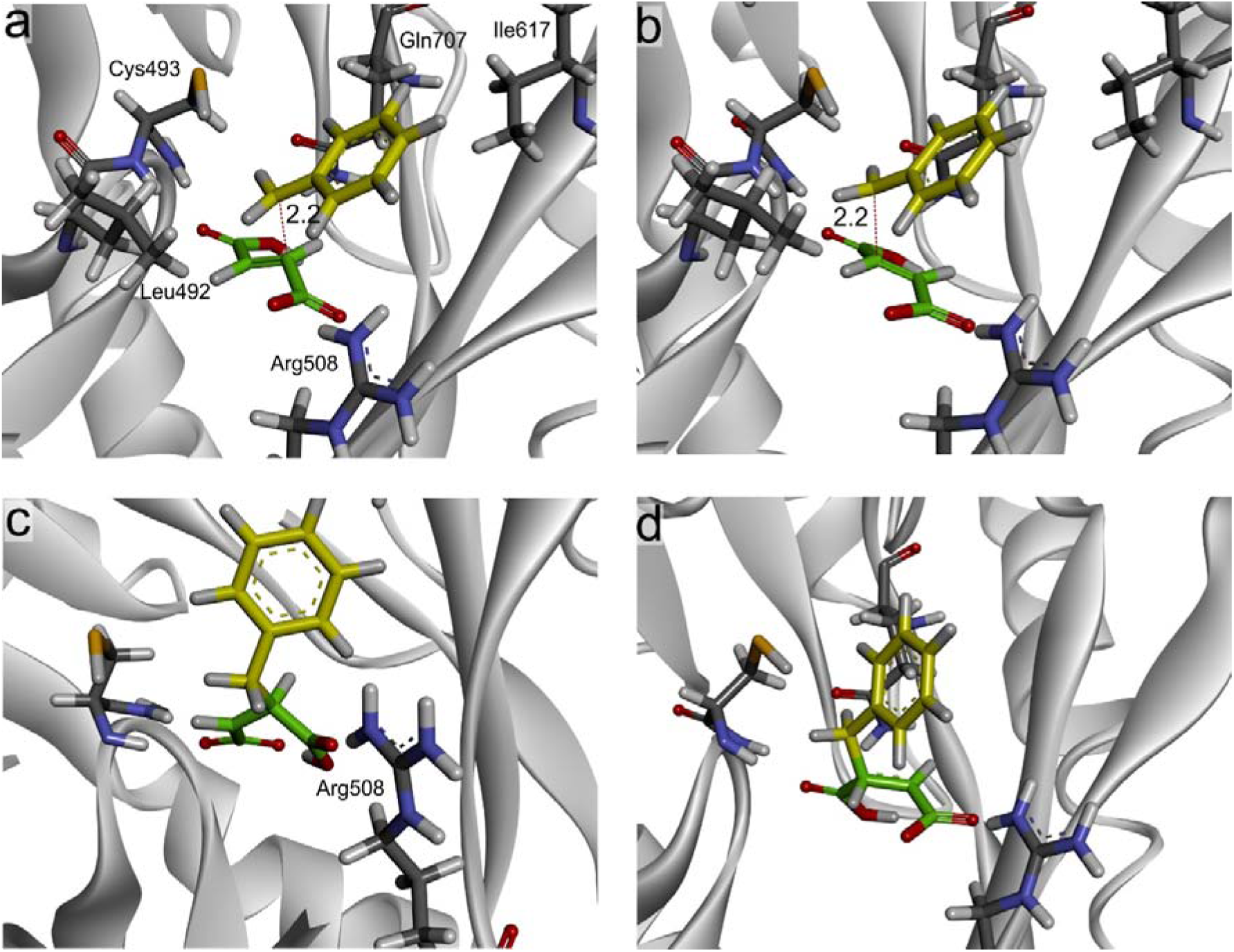
Formation of the C-C bond: a) TS3a attack of radical toluene on the distal C atom of the fumarate double bond b) TS3b attack of radical toluene on the proximal C atom of the fumarate double bond; radical benzylsuccinate intermediates c) I3a and d) I3b.

The completion of the reaction leads to the formation of an (*R*)-benzylsuccinyl radical intermediate, I3a, with a newly formed C-C bond of 1.55 Å and a radical at the proximal C carbon (Fig. 5C). The change of hybridization of the distal C atom from sp^2^ to sp^3^ bends the acidic part of the benzylsuccinyl molecule, bringing the two carboxylic groups together. As a result, an internal H-bond is formed between the carboxylic groups, and the proton shifts spontaneously between the formerly protonated and deprotonated carboxyl groups (Fig. 4C). This effect also significantly weakens the electrostatic interaction with Arg508 and may later help in the release of the product.

In the alternative TS3b, the new C-C bond is formed with the benzyl radical approaching the proximal C atom of the double bond of fumarate. In this case, the C-C bond distance is 2.2 Å and the C^dis^-C^prox^-C^meth^ angle is 108° with the bend towards Cys493. Similarly, as in the case of TS3a, the plane of the benzyl radical is not parallel to that of the fumarate, but slightly bent to one side (Fig. 5B). Although the proximal C atom is also drawn out of the plane of the fumarate towards the benzyl radical, the distortion is significantly smaller than in TS3a, indicating higher sp^2^ character in TS3b. The spin density is evenly distributed between the benzylic carbon (0.56) and the distal carbon atom of the double bond of fumarate (0.54), while the proximal carbon atom exhibits a negative spin value of -0.16. The discrepancies in charge distribution indicate a significant difference between TS3a and TS3b. In the former (TS3a), the benzyl radical part is almost neutral (-0.22), but the benzylic atom and the fumarate cosubstrate are highly negatively charged (-0.74 and -1.17, respectively), whereas in the latter (TS3b) the benzyl radical part is positively charged (0.47) while the negative charge of the fumarate cosubstrate is significantly lower (-1.7). This difference seems to be associated with different protonation patterns of the intermediate product and its polarization by Arg508.

The formation of the C-C bond leads to the production of an (*R*)-benzylsuccinyl radical intermediate shown in I3b (Fig. 5D). Unlike in the case of I3a, the carboxyl groups are still far apart from each other and no internal H-bond is formed between them. As a result, the distal carboxyl group still forms a strong salt bridge with the side chain of Arg508 which will have a profound influence on the next step of the reaction.

The analysis of the energy profile (Fig. 8) indicates that the formation of the C-C bond proceeds preferentially at the distal C atom, as the energy barrier for TS3a is 64.2 kJ/mol while that for TS3b is 110.4 kJ/mol. This high difference in energy barriers is associated with the steric constraints imposed on the formation of the C-C bond at the proximal C atom, which is in close vicinity to Cys493 and Leu492. Also, the absolute energy of I3a is significantly lower than that of I3b (-15.4 vs 39.5 kJ/mol).

### Step 4 – quenching of the benzylsuccinyl radical intermediate

The final step of the reaction is associated with quenching of the radical intermediate. This step involves HAT from Cys493 to the radical carbon atom. During the optimization of this step, yet another conformational shift occurred, which resulted in a decrease of total energy by 73 kJ mol^−1^. This conformational shift was associated with a flip of Arg826 combined with the relaxation of the loop containing Gly829, as well as a small accommodation of the SH group of Cys493. Besides these details, the models are identical (RMSD 0.076) and the residues that changed their conformations are not in any contact with the reactants.

As a consequence of the two potential pathways considered in step 3, the conversion of the benzylsuccinyl radicals to the product benzylsuccinate can also proceed along two variants. Pathway a involves the HAT to the proximal C2 atom of the benzylsuccinyl radical intermediate (TS4a) while pathway b requires HAT to the distal C3 atom (TS4b) – see Fig. S13.

Furthermore, quenching of the radical intermediate has been shown experimentally to proceed usually in a *syn* geometry (i.e. at the same face of the double bond as the C-C addition)^60^. However, we recently reported results of an H/D exchange experiment^48^ indicating that *R*-benzylsuccinate is slowly deuterated upon incubation in D_2_O in the presence of active BSS, producing d_1_-benzylsuccinate as a major product, but also traces of d_2_-benzylsuccinate at a significantly lower rate ^48^. These observations indicated on one side that the last reaction steps are reversible, while on the other side, the HAT reactions involved in quenching of the benzylsuccinyl radical are not absolutely enantioselective and H/D atoms may occasionally be transferred or removed from the other (*anti*) face of the intermediates. Therefore, we decided to investigate a potential *anti*-addition variant of the pathway a (using a β exponent to denote *anti* addition, in contrast to α for *syn* addition) (Fig. S13).

After the C-C bond is formed at the distal C atom in pathway a, the radical is localized at the proximal carbon atom, close to Cys493. This allows a relatively easy HAT from the SH group, which at TS4^α^ exhibits almost linear geometry (S-H-C angle of 171°) and 1.52 Å distance between the H and both S and C atoms (Fig. 6a). Because this transition state is reached with minimal rearrangement of the intermediate, its barrier turns out to be the lowest (45.1 kJ/mol – see Fig. 8). The majority of the spin density at the TS is divided between the S and C atoms (0.37 and 0.65).

**Figure 6.**
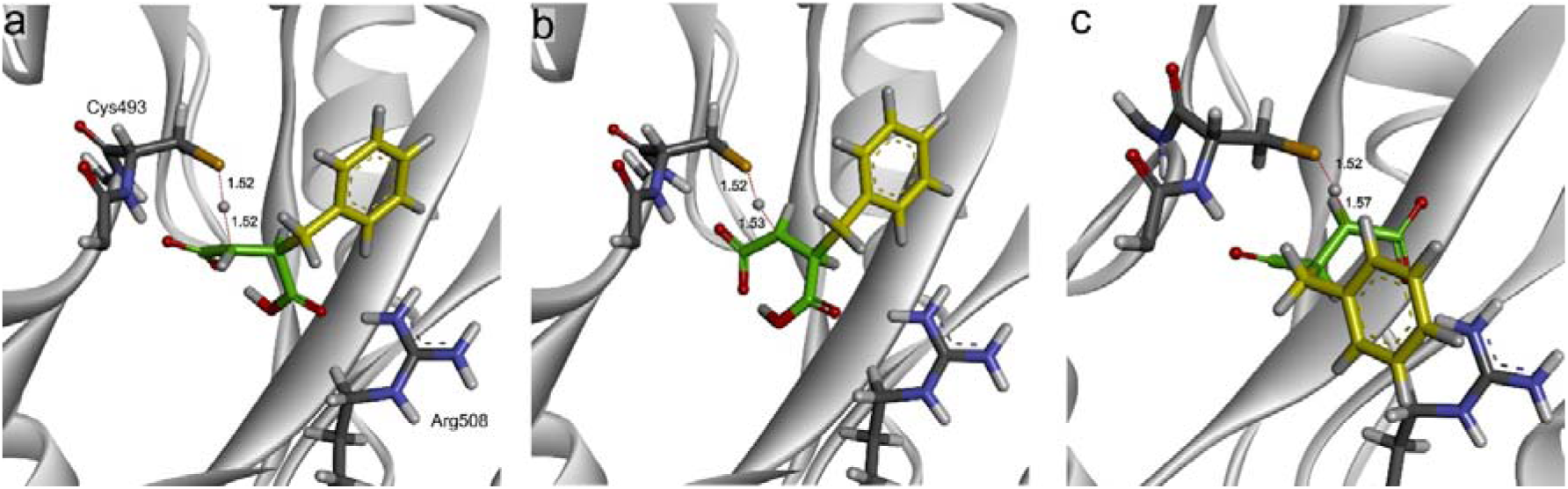
TS4 for quenching the radical benzylsuccinate intermediate by HAT from Cys493: a) TS4^α^ a *syn* addition to the proximal C2 atom, b) TS4^β^ an *anti*addition to the proximal C2 atom, c) TS4b *syn* addition to the distal C3 atom.

In the TS4^β^a (Fig. 6b) the H atom from the SH group of Cys489 would be transferred in the *anti*-geometry to the benzylsuccinyl radical, which requires partial rotation of the now deprotonated C1 carboxyl group, removing it from the H-bonding pocket. As a result, despite the similar geometries of TS4^β^a and TS4^α^a (C-H and S-H distances of 1.53 and 1.52Å, 170° S-H-C angle), its energy barrier is quite high (90.9 kJ/mol -see Fig. 8).

Quenching of the benzylsuccinyl radical formed by addition at the proximal C atom in pathway b turns out to be more difficult. This is because the radical atom is located far from Cys493 on the distal C-atom, and to reach TS4b the intermediate has to rotate in the active site (Fig. 6c). This also implies that the deprotonated carboxyl group has to move away from the positively charged guanidyl group of Arg508. The geometry of TS4b resembles that of TS4a with an almost linear angle between S-H-C (170°) and slightly longer C-H distance with respect to S-H distance (1.57 vs 1.52 Å) while the transfer of the spin density is less advanced than in case of TS4a (0.29 and 0.72 at S and C atoms, respectively). The energy of the TS4b is very high (93.4 kJ/mol) but the energy difference between I3b and TS4b is in fact smaller (53.9 kJ/mol) than that between I3a and TS4a (60.5 kJ/mol). This is mainly due to the highly elevated energy of I3b.

The completion of HAT in both cases, results in the formation of *R*-benzylsuccinate and the obtained I4a state exhibits significantly lower energy (-29 kJ/mol) compared to that of I4b (-2.9 kJ/mol). Even though in both states, I4a and I4b, the enzyme is in the complex with *R*-benzylsuccinate, I4b has higher energy due to steric clashes of the benzyl ring positioned closer to the cysteine.

### Step 5 – HAT between Gly829 and Cys493 for the E:P complex

For over two decades, it was postulated that the last step of the BSS mechanism is the final transfer of the radical from Cys493 to Gly829 (Fig. 3). This is due to the fact that the thiyl radical is not observed in EPR. Recently, we have analyzed this step in detail^48^ and demonstrated that the process preferentially proceeds via a *re*-side attack. In order to take over the H atom from Gly829, Cys493 has to change its conformation again, i.e. rotating counterclockwise from the position left after quenching the benzylsuccinate radical (Cys93 C-Cα-Cβ-Cγ dihedral angle of 182°) to the conformation closest to Gly829 (dihedral angle 52°). The MD simulations conducted for the E:P complex containing a radical Cys493 and a monoprotonated benzylsuccinate with its protonated group facing Arg508 indicate a stark difference in the population of the three main rotational energy minima of Cys493, compared to that of the E-Cys^rad^:S complex (Fig. S14 and S7). The population with C-Cα-Cβ-Cγ dihedral angles around a local maximum of 172°±22° turned out to be the most probable (68% of simulation time), followed by the populations around dihedral angles of 281°±27° (28%) and 49°±21° (only 4%). Consistently with the MD results, QM:MM modelling revealed that the conformation change of Cys493 from that in I4a to the one allowing HAT to Gly829 (I4a rot i.e. with dihedral angle of 54°) is associated with an increase of energy by 6.1 kJ/mol.

The calculated C-H and S-H distances of TS5 are 1.57 and 1.48 Å, respectively, and the S-H-C angle is 166.5° (Fig. 7). Analysis of the energies indicates that the HAT between Gly829 and Cys493 in the E:P complex is highly endergonic, requiring more than 40 kJ/mol (Fig. 8). This step is energetically symmetrical to step 1, i.e. the barrier for transferring the H atom from Gly to the thiyl radical is higher than the barrier of its transfer from Cys493 to the glycyl radical (99 kJ/mol vs 57 kJ/mol). In our earlier study on the HAT reaction between the glycyl and thiyl radical states of BSS, we obtained very similar results with either substrate- or product-bound BSS, while apo-BSS with an empty active site showed the reverse behaviour, i.e. lower energy of the glycyl-than the thiyl-radical state ^48^. Thus, our calculations indicate, that HAT between Gly829 and the thiyl radical would be the rate-limiting step of the overall reaction if it proceeded with bound product in the active site (with a barrier of 75.2 kJ/mol). Since this would effectively mask any kinetic isotope effect associated with labelled substrates, we have to assume that the last step of the reaction cycle proceeds via a different pathway. After completion of the C-C addition reaction, the radical appears to remain localized at Cys493 more probably than to move to Gly829 in the E:P complex. This situation obviously needs to be resolved to close the catalytic cycle, addressing the mode of product release. One may speculate that either glycyl/thiyl HAT occurs in a concerted reaction with the release of benzylsuccinate, or benzylsuccinate is already released from the thiyl state of BSS, followed by an energetically favourable glycyl/thiyl HAT in the apoenzyme ^48^. Unfortunately, the product release, as well as the binding of the substrates concomitant with restructurings of the enzyme and transfer of radical hydrogen, cannot yet be modelled with the currently available QM:MM or even QM:MM:MD methodology.

**Figure 7.**
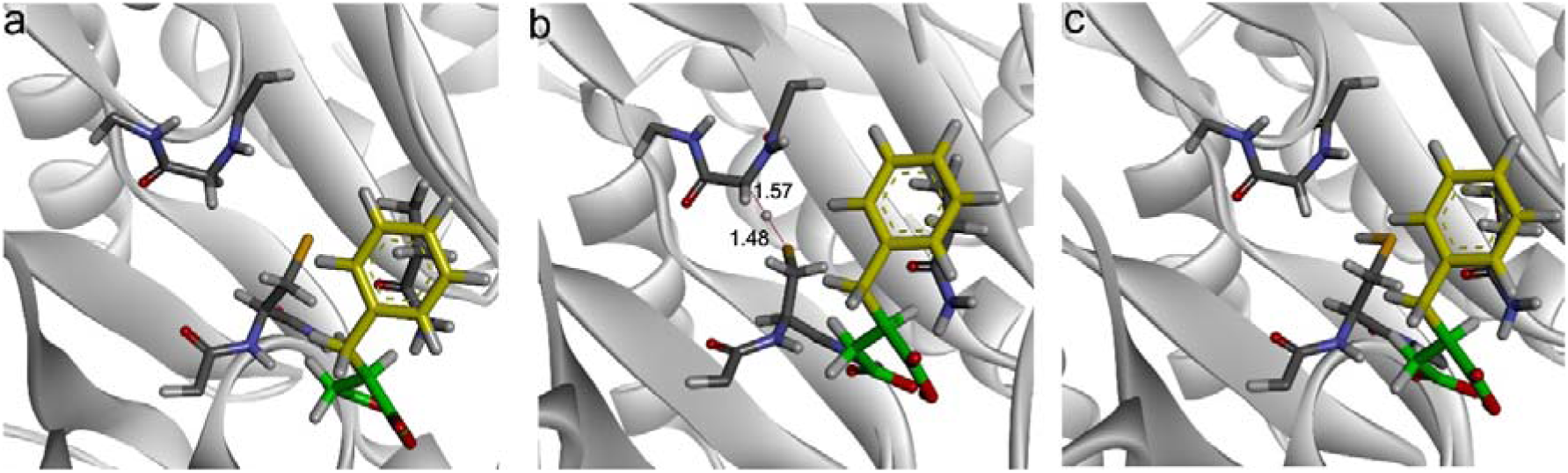
HAT between Gly829 and Cys493 for the E:P complex: a) I4-rot radical Cys rotated to the most stable conformation, b) TS5 pro*R* H atom transfer from Gly to Cys, c) radical Gly

**Figure 8.**
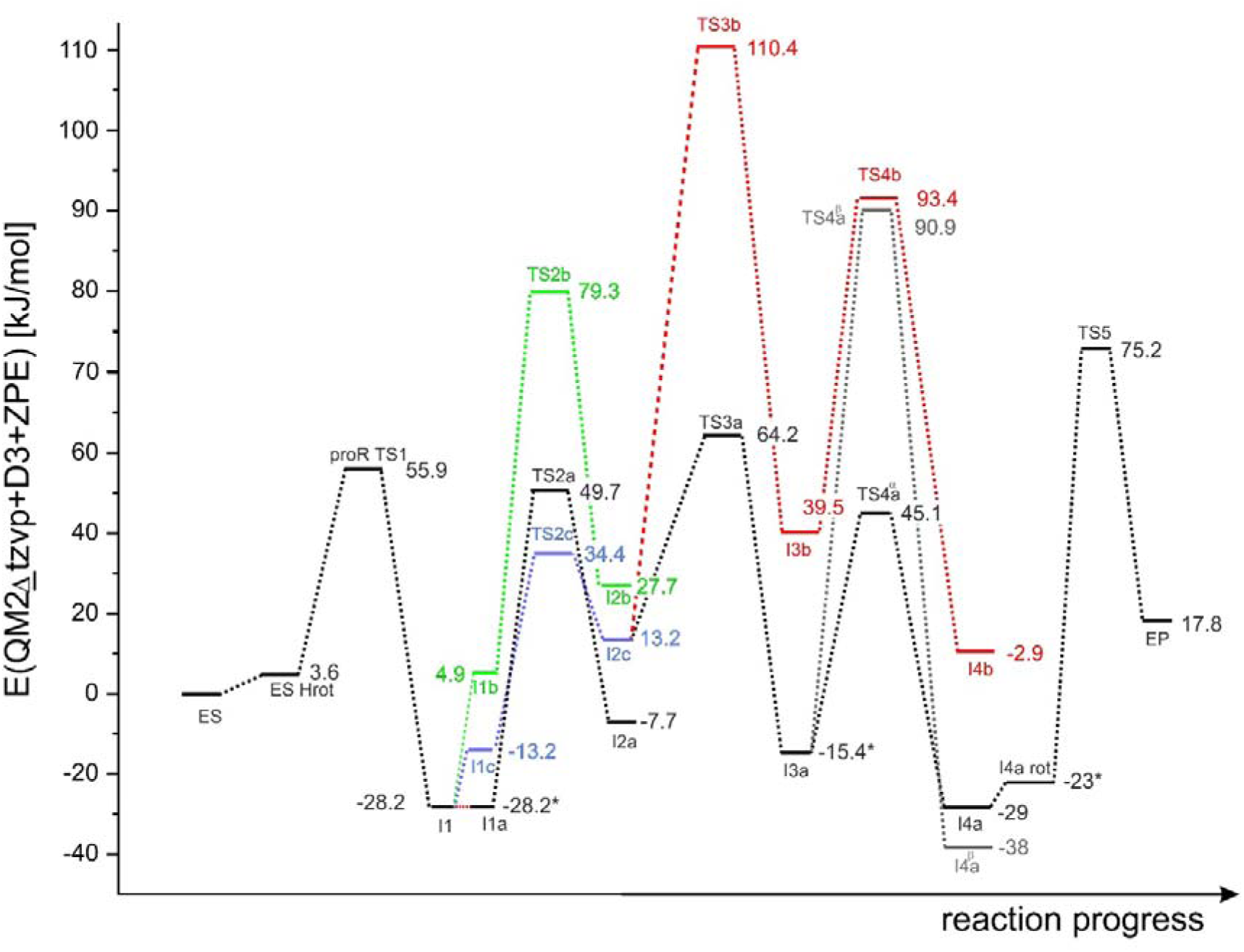
Energy diagram for pro*R* conformation of the fumarate; * indicates site of profile matching due to conformation changes of the model along the reaction.

### Kinetic isotope effect

One of the ways to test mechanistic hypotheses is to compare experimental data with the results of modelling. As the formation of benzylsuccinate from fumarate and toluene involves multiple transfers of an H atom, experiments with deuterium-labelled toluene substrates should lead to a kinetic isotope effect (KIE). It was previously demonstrated that significant KIE is observed with fully deuterated toluene, although reports were inconclusive on its precise value^61, 62^. However, modelling provides a means to calculate intrinsic KIE (i.e. effects associated with a particular molecular step of the reaction), and this can be used to estimate k_cat_. Calculation of the ratios of such rates for unlabeled and labelled substrates yields the predicted value of KIE that should be directly comparable to those obtained from experiments. Interestingly, such a method was previously applied for the study of the BSS mechanism by Li & Marsh^61^, where the experiment was compared with barriers derived from the gas-phase model^63^ of the reaction that did not take into account the influence of the enzyme.

We decided to use the same approach for initial reaction conditions, assuming however that the irreversible step is quenching of the benzylsuccinyl radical followed by the release of the product, and not the C-C bond formation step as suggested by the gas phase model ^64^. This is due to the fact that quenching of the benzylsuccinyl radical is associated with the lowest kinetic constant value of all partial steps of the mechanism (see Tables 1 and S5-6) when calculated with respect to the preceding stationary point.

Assuming quenching of the product radical as a rate-limiting step, the rates of the overall reaction should only depend on the concentration of the BSS enzyme containing the cysteine with SH group and benzylsuccinyl radical [BSS-C:BS.]:.

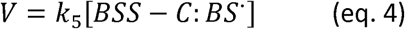

We assume that the reverse reaction (i.e. binding of benzylsuccinate and its conversion to the radical) can be neglected at the initial stage of the reaction, due to the low concentration of the available product, which results in a low probability of its binding back and reversing the reaction. With these assumptions, we have derived an equation describing the reaction velocity using the King-Altman algorithm^58^ (see SI for full derivation).

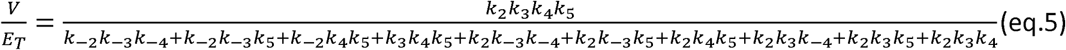

Using this approach, we calculated k_cat_ using kinetic constants of elementary steps derived from our QM:MM modelling, as calculated for toluene or d_8_-toluene. The predicted k_cat_ for toluene was 0.54 s^−1^ while that for d_8_-toluene was 0.21 s^−1^, which resulted in a predicted KIE of 2.56. The predicted value of KIE appears in very good agreement with the experimentally obtained value of 2.13±0.1, the kinetic isotopic effect on V_max_ determined in a direct assay (Fig. 8A). Values of KIE on 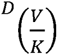 obtained in the competitive experiment (Fig. 8B) from three independent reactors were 4.01, 3.5 and 3.56, yielding an average value of 3.69±0.16. As the 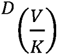 involves kinetic constants associated with substrate binding and release we can assume that the increase of the isotope effect in the competitive experiment is caused by the differences in enzyme affinity toward the toluene compared to binding of d_8_-toluene (i.e. higher K_m_ for d_8_-toluene).

### Reaction enantioselectivity

The reaction catalyzed by BSS is highly enantioselective, yielding exclusively (*R*)-benzylsuccinate as the product. In a previous study, we used cluster models to rationalize this behaviour, which indicated that (*S*)-benzylsuccinate can only be formed when the fumarate is bound in a conformation presenting its other face (pro*S*) to the attack by the benzyl radical. While binding of fumarate in this conformation can easily be envisaged, the pro*S* binding mode results in subtle changes in the positions of the proximal and distal carbon atoms capable of reacting with the benzyl radical.

To check if fumarate binding in either of the two poses provides any thermodynamic preference for the reaction, we have conducted MD simulations for the pro*R* and pro*S*-BSS:fumarate complexes. However, no significant differences were obtained between these two configurations, and any deviations were within the error of the method, especially if the variability introduced by repeated simulations is taken into account. The average ΔG values of binding according to the “Generalized Born” method were in the range of -46 to -32 kcal/mol for pro*R* fumarate and -46 to -37 kcal/mol for pro*S* fumarate, with SD in the range of 3-4 kcal/mol. A similar effect was observed for the ΔG values of fumarate binding according to the Poisson-Boltzmann method (see Supporting Information for more details, Fig. S15).

As the ΔG of fumarate binding seems not to be responsible for the observed pro*R* enantioselectivity of the reaction, we decided to conduct a similar analysis of the MD trajectories for the model with fumarate in pro*S*-conformation as in the case of the pro*R* pathway. We analyzed the dynamic behaviour of toluene and its approach toward the proximal or distal C atoms of the double bond of fumarate. Analysis of the distances from three independent MD simulations indicated that the toluene indeed is able to approach the fumarate bound in pro*S* orientation more easily at the proximal C3 atom than at the distal C2 atom (minimal and median distances 3.02 Å and 7.2 Å or 3.13 Å and 7.8 Å for the proximal or distal C atoms, respectively – Fig. S16). This pattern of preferred attacks is directly opposed to that observed with fumarate bound in pro*R* orientation. The simulations with fumarate in pro*S* orientation also show larger average distances to the toluene, compared to those with fumarate in pro*R* orientation (medians 5.6 and 6.5 Å for the distal and proximal C atom, respectively, Fig S11).

Furthermore, we analyzed the pro*S* reaction pathway also by QM:MM modelling, starting from the benzyl radical intermediate of the standard pathway (I2c) assuming that the binding mode of fumarate in either pro*R* or pro*S* conformation does not significantly influence H abstraction from toluene. A detailed description of the transition states is provided in the supplement (Figs. S12 and S13, Table S2 and S4). Unexpectedly, we observed the same regioselectivity pattern as in the case of the pro-*R* pathway i.e., a kinetic preference of the benzyl radical to attack on the distal carbon atom (C2), compared to the proximal C3 atom of the double bond of fumarate (44.7 vs 92.9 kJ/mol for TS3a vs TS3b, Fig. 10). The formation of the *S*-benzylsuccinyl radical is followed by a radical quenching process, which is associated with barriers of 60.3 kJ/mol (TS4a) and 114.6 kJ/mol (TS4b) for the quenching of proximal and distal radicals. Noticeably, the energy barrier for the C-C addition even along the kinetically preferential pathway a (at a distal C2 atom) is significantly higher (81.6 kJ/mol) than that of the pathway pro*R* (60.5 kJ/mol), thus favouring the production of *R*-over *S*-benzylsuccinate. The preferential mode of C-C bond formation is followed by a radical quenching process at a lower energy barrier (TS4a 60.3 kJ/mol) and would suggest the formation of *S*-benzylsuccinate.

**Figure 9.**
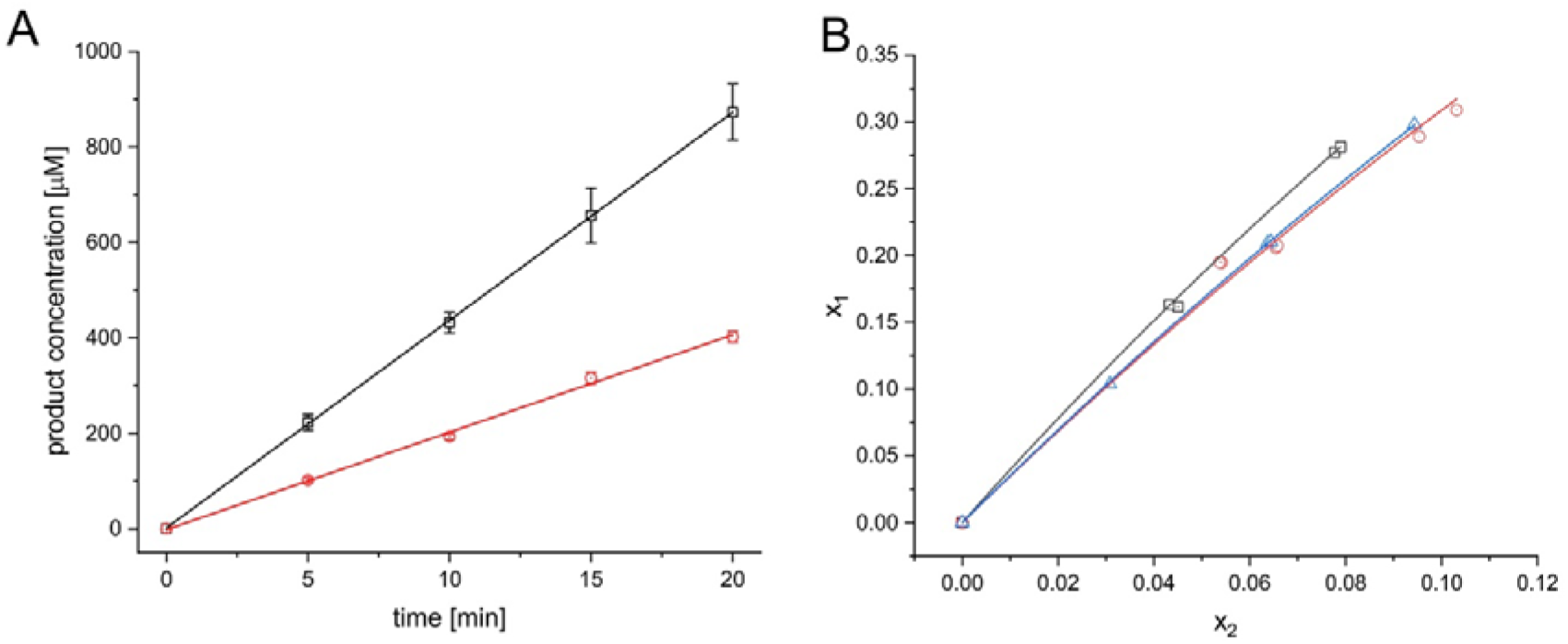
The results of the kinetic isotope effect experiment: A) direct measurement with a kinetic curve for toluene (black, squares) and d_8_-toluene (red, circles); error bars depict standard deviation from 3 experiments, B) competitive measurement – 3 independent replicas.

**Figure 10.**
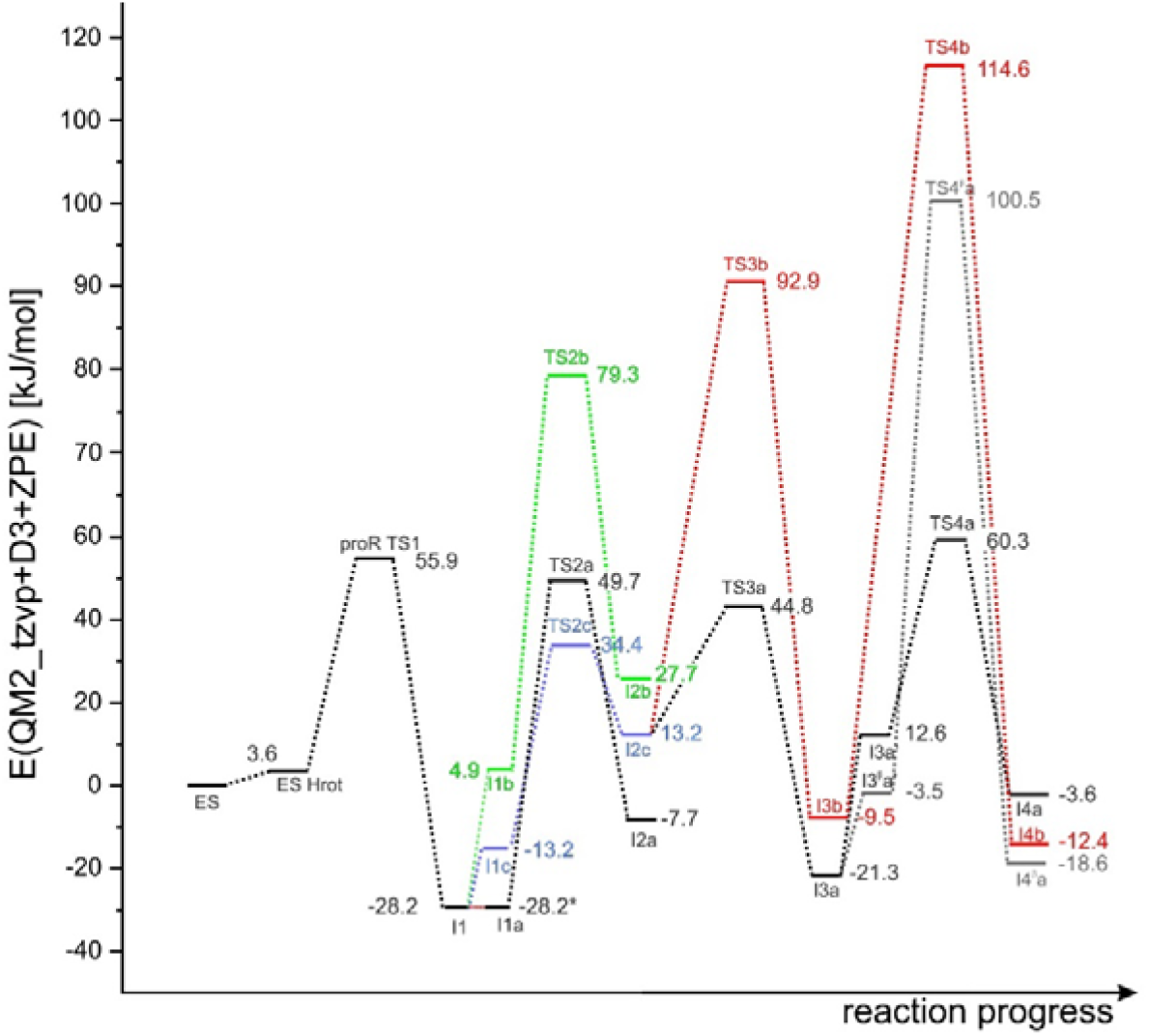
Energy diagram for pro*S* conformation of the fumarate. The pro*S* profile was matched at I2c.

**Figure 11.**
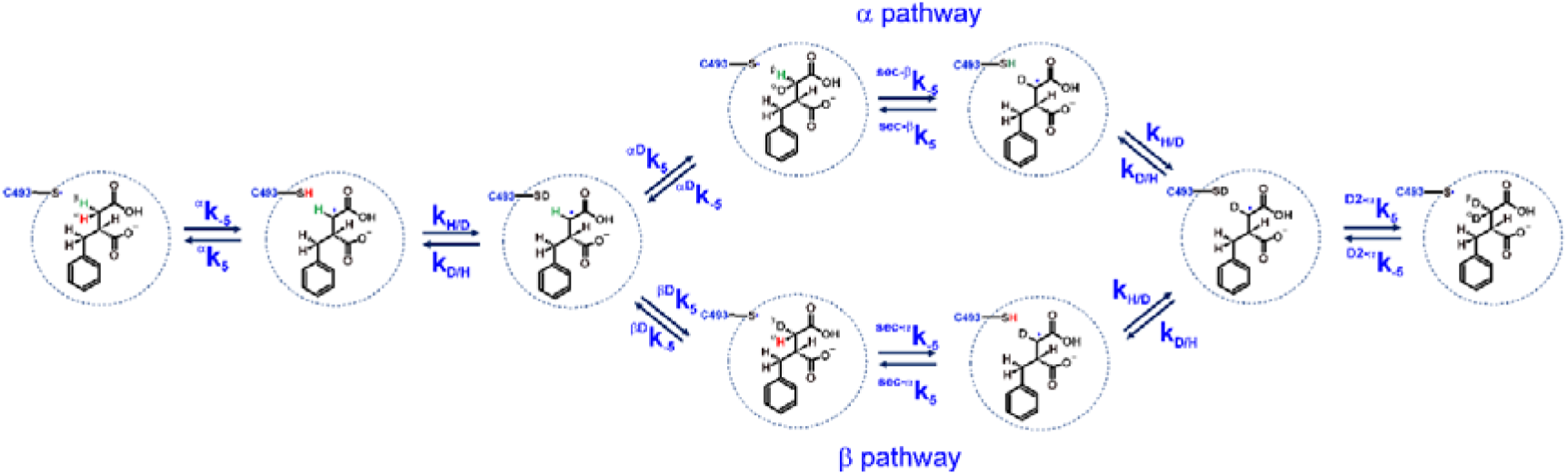
Two potential pathways leading to the formation of d_2_-benzylsuccinate product.

Interestingly, we also detected a different protonation mode of the *S*-benzylsuccinyl radical in the pro*S* pathway. Compared with the pro*R* pathway, there is no internal H-bonding detected between the carboxyl group of the product radical (I3a’ and I3^β^a’ - Fig. 10 and Table S4). This may contribute to the specificity of the enzyme for *R*-benzylsuccinate production because of the elevation of the energy barrier for the radical quenching step.

Finally, similarly, as in the pro*R* pathway, the HAT in *anti* conformation was associated with a significantly higher barrier (TS4^β^a 100.5 kJ/mol) which shows that the d_2_-benzylsuccinate formation observed in the experiment is not a result of unspecific exchange in the S-product.

We used the potential energy profiles obtained for both pro*R* and pro*S* reaction pathways, kinetic constants calculated for elementary steps of both pathways and kinetic rate constant calculated according to eq. 5 to elucidate the enzyme enantioselectivity. The estimated k_cat_ for reaction with toluene at 303 K along the pro*R* pathway was 0.12 s^−1^. When the calculation of the k ^S^k,^S^k, and ^S^k_5_ are used (Table 1) its value drops to 9.6*10^−4^ s^−1^. This means that the reaction leading to *R*-benzylsuccinate is 123-fold faster than that yielding *S*-benzylsuccinate. In case the nuclear tunnelling is taken into consideration the estimated k_cat_ for reaction along the pro*R* pathway was 0.19 s^−1^ while for pro*S* pathway 2.2*10^−3^ s^−1^ leading to production of *R*-benzylsuccinate 89-fold faster than *S*-benzylsuccinate. The estimation of the tunnelling effect also indicates that the overall rate may be accelerated by a factor of 1.6 for the pro*R* pathway and 2.3 for the pro*S* pathway, which indicates, that tunnelling acts against enantioselectivity of the enzyme although it does not compromise to any significant degree. The barrier-dependent kinetics is additionally modulated by a higher probability of C-C bond formation in the pro*S* pathway at the proximal C atom of the fumarate which would even further decrease the observed rate toward the *S*-product.

**Table 1.**
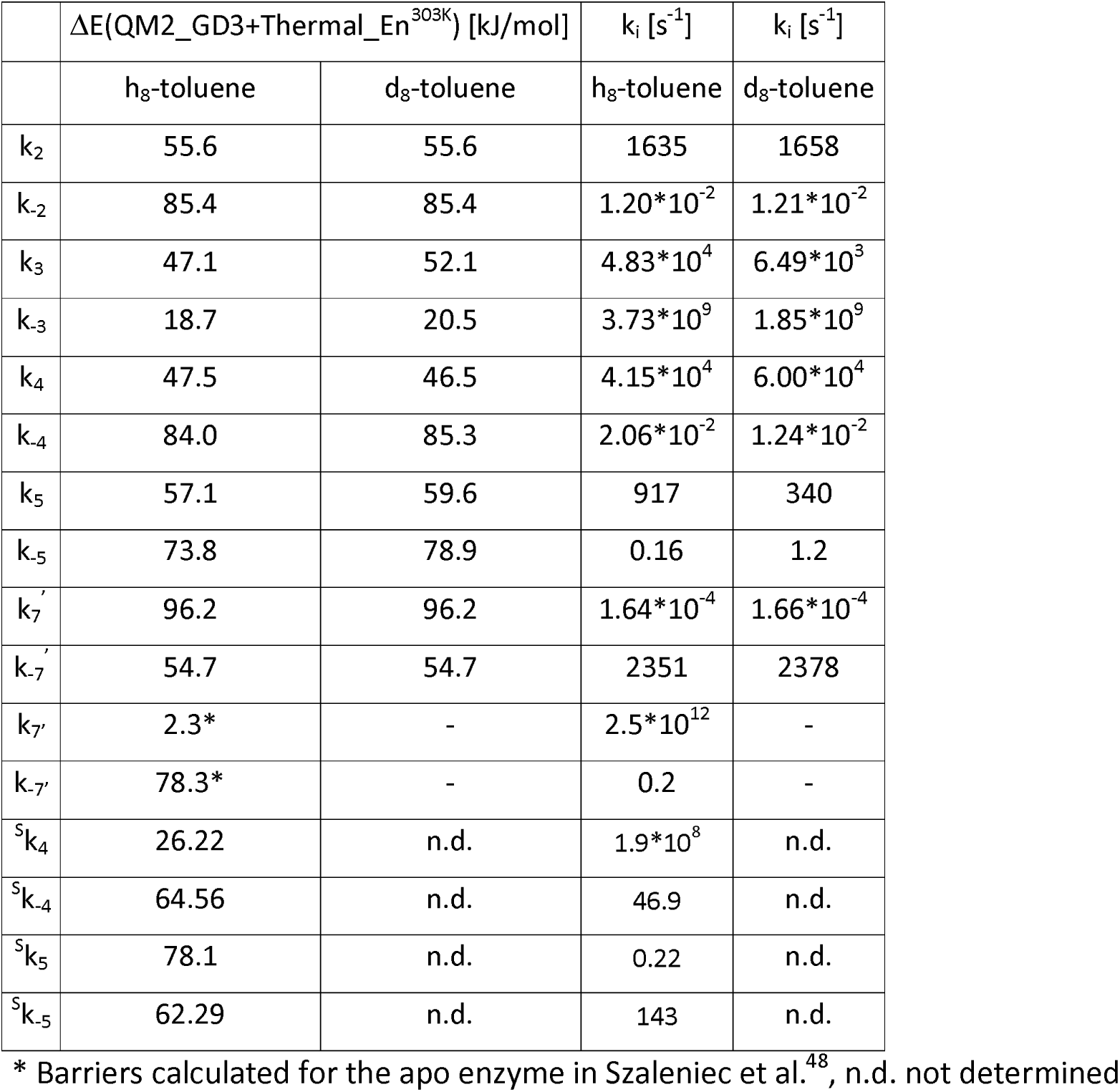
The kinetic constants calculated from thermal energy barriers in QM:MM for the pro*R* pathway with toluene and d_8_-toluene as well as for the 2 steps of the pro*S* pathway with toluene.

The analysis of values of Wigner’s tunneling correction also points out the kinetic steps that will be especially prone to the below-the-barrier tunneling acceleration. Interestingly, the highest acceleration factors were predicted for the final step of the reaction i.e., HAT from Gly to Cys in E:P complex (TS5, 3-fold acceleration) and the quenching of the benzylsuccinyl radical intermediate (TS4, 2.3-fold acceleration). As can be expected the weakest influence of the tunneling was predicted for a broad-barrier C-C bond formation (1.2-fold acceleration).

### H/D exchange

Recently we have described the isotope exchange of a D atom to H during a reaction catalyzed by BSS between d_8_-toluene and fumarate in H_2_O as well as a reverse exchange of H atom(s) to D during incubation of benzylsuccinate in D_2_O with BSS^48^. In the reaction pathway, the only possible step at which a deuteron can be exchanged for a hydrogen atom is when it is transferred from isotope-labelled toluene to Cys493, as the Cys residue readily forms a hydrogen bond with Gln707. Due to the lack of a stable expression system for BSS and its very sensitive nature to oxygen deactivation, it is not possible to directly measure the H/D rate on cysteine according to the NMR-based method proposed in the literature ^65^. Our kinetic measurements of H/D exchange are based only on a few experiments and provide only semi-quantitative data due to the inherent inconsistency in assaying the highly labile BSS activities. Despite that, we decided to use our calculations to estimate the most probable mechanism of the H/D exchange which leads to the formation of both d_1_- and d_2_-substituted benzylsuccinate during incubation in 40% D_2_O. To approach this problem, we have analyzed the potential pathways leading to the formation of d_1_- and d_2_-substituted benzylsuccinate (Fig. 10).

First, we examine the pathway of forming d_1_-benzylsuccinate from benzylsuccinate. Such a process requires the removal of an H atom from C2 of the product from the kinetically preferred α position at the rate ^α^k_-5_, followed by H/D exchange at the Cys-SH group and final back-transfer of the D atom to the benzylsuccinyl radical at the rate ^αD^k_5_ (Fig. 10, Table S11). To derive the kinetic equation describing the rate of formation of d_1_-benzylsucciante we assumed the following prerequisites: i) formation of d_1_-benzylsucciante is the irreversible step due to a surplus of unlabelled and very low concentrations of the deuterated product, ii) the enzyme is in equilibrium with benzylsuccinate, so the binding or release of the product is not influencing the observed exchange rate, iii) k is a product of k^HDX^ and the concentration of D^+^ in the enzyme, while k_D/H_ is a product of k^HDX^ and concentration of H^+^ in the enzyme. The latter assumption stems from the fact that the enzyme is in equilibrium with its solvent and as we do not know anything about the molecular mechanism of H/D or D/H at Cys493, we assume that the availability of protons/deuterons in the active site is the only relevant parameter. With these assumptions in mind, we have constructed the scheme (Scheme S1) of the reaction according to the King-Altman procedure (derivation eq. 9-14).

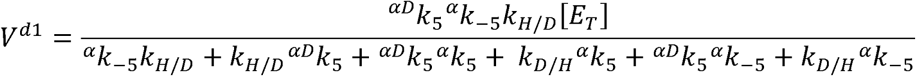

where V^d1^ is the reaction velocity of d_1_-benzylsucciante formation and [E_T_] is the total concentration of the enzyme In contrast, d_2_-benzylsuccinate can be obtained along two alternative mechanisms depicted in Fig. 5. After the formation of the benzylsuccinyl radical and H/D exchange at Cys493, the deuteron can be either transferred to the α position (i.e. TS4^α^a, kinetic constant ^αD^k_5_) or to the β position (i.e. TS4^β^a, kinetic constant ^βD^k). In the first case, the reaction continues by removal of the β H atom (kinetic constant ^sec-β^k_-5_), which is influenced by secondary KIE introduced by the D atom at the same C atom, while in the second case, the α H atom will be removed at a rate (kinetic constant ^sec-α^k_-5_) which is also modulated by secondary KIE. Both pathways lead to the formation of a d_1_-benzylsuccinyl radical. Then the d_2_-product can be obtained after another H/D exchange process at Cys493 and transfer of the D atom to the d_1_-benzylsuccinyl radical via the preferred α pathway at a rate (^D2α^k) that is influenced by primary and secondary KIE. The calculated kinetic constants for both α and β pathways are collected in Table S11. Again, as in the H/D exchange the proton/deuteron shifts are influenced by KIE, we decided to use energies corrected by thermal correction instead of Gibbs free energies. However, to test the robustness of our derivation we have also calculated the same rates using DG, obtaining qualitatively the same results (see SI). From these schemes (Scheme S2 and S3) we assumed that the last step of the process is irreversible due to the extremely low concentration of the products compared to the concentration of unlabelled benzylsuccinate. We have derived the overall equations for reaction velocities of d_2_-benzylsuccinate formation (V^d2^) for both pathways using the King-Altman method^58^ (see eq. S23-29 as well as Schemes S3-S4).

To compare the values of V^d1^/[E_T_] and both pathways of V^d2^/[E_T_] we need to assume the value of k^HDX^ from which the k^H/D^ and could be derived. Similarly, as in the case of k_6_, we have no way to measure this rate and the mechanism of the H/D exchange in BSS was not investigated. In order to estimate the range of potential kinetic constant of H/D exchange we decided first to approximate the rate of d_1_-benzylsuccinate formation observed in the experiment. The average rate of the d_1_-product formation was 1.34*10^−10^ M s^−1^ (1.94 μM in 4 h) ^48^. The process was catalyzed by 9.8 mg of the cell extract of which approximately 10% is BSS with a molecular weight of 226 kDa, yielding a concentration of 4.3*10^−9^ M. However, only one active site per heterohexadimer is active so the final estimated rate of d_1_-production is 0.06 s^−1^. Assuming a 4:6 ratio of k_H/D_ : k_D/H_ based on the 40% D_2_O concentration in the experiment, we approximated k_H/D_ as 111 s^−1^ and k_D/H_ as 167 s^−1^. These values were obtained based on a value for k^HDX^ of 5 s^−1^ M^−1^, assuming 22 M concentration of D_2_O and 33 M of H_2_O. With these assumptions, we obtained a value for V^d1^/[E_T_] of 0.06 s^−1^, while that of V^d2^/[E_T_] was 5.9*10^−11^ s^−1^ or 1.37*10^−10^ s^−1^ for pathway α or β, respectively. These results indicate that the double exchange via the β pathway is 2.3 times faster than via the α pathway which indicates that i) the β pathway is the preferential mechanism for the formation of d_2_-benzylsucciante and ii) the d_1_- and d_2_-products are being formed in a different manner. Furthermore, the analysis of relative concentrations of species along the d_2_-pathways indicates the highest concentration of α-D-benzylsuccinate for α pathway (65%), which suggests that at this point the monosubstituted product has a chance of dissociating. This is because the next step, the removal of the H-atom from the beta position is the rate-limiting step of the whole pathway. In contrast, along the β pathway, the highest concentrations are predicted for [Cys-S.:BS] and [Cys-SH:BS.] (approx. 50%) and the concentration of d_1_-product is negligible. It should be underlined, however, that experimental concentrations of d_2_-product were far higher than that predicted by microkinetic analysis (only 4.4-fold lower than d_1_-product). This indicates that the barrier for the exchange of β H atom may be significantly overestimated and in reality, the process has to proceed faster.

### Effect of tunnelling on HDX process

If Wigner tunnelling correction is included in the H/DX analysis one has to slightly adjust fitted kD/H and kH/D to match the approximated experimental rate. A new k^HDX^ is fitted as 4.2 s^−1^ M^−1^ which decreases the H/D and D/H rate constant to 93 and 140 s^−1^, respectively. Surprisingly, despite the acceleration of all transfers in the range of 1.9-2.5 the overall rates of HDX exchange leading to the d_2_-product decrease, yielding 1.4*10^−10^ s^−1^ for the α pathway and 2.54*10^−10^ s^−1^ along β pathway.

## Discussion

We show in this study the first detailed QM:MM modelling of the reaction mechanism of BSS from the substrate- to the product-bound state of the active site. While the model confirmed the previously proposed individual steps of the mechanism, we learned unexpected details on the involvement of active site residues and their movement during the reaction cycle. In particular, the essential Cys493 needs to rotate around its C_α_-C_β_ bond to achieve the necessary HAT reactions involved in generating the various radical intermediates of the mechanism. For its initial conversion into a thiyl radical (step 1), Cys493 must get close to and face the stable glycyl radical on Gly829 by assuming a C-C_α_-C_β_-S_γ_ site cavity, while the glycyl radical resides outside of the cavity. After the thiyl radical is formed, Cys493 rotates in counterclockwise orientation to reach a dihedral angle of 179° which offers the lowest-energy barrier for abstracting an H atom from the methyl group of a bound toluene (step 2) and the only available option for adding it to fumarate with inversion of its stereochemical conformation. The subsequent C-C bond formation by adding the generated benzyl radical to the double bond of fumarate (step 3) has to occur at the distal C-atom of fumarate for a productive reaction. The following step of transferring the H-atom back to the benzylsuccinyl radical from Cys493 (step 4) only works reasonably when the radical is positioned at the proximal C-atom. Therefore, it must be expected that the C-C bond formation is reversed if it occurs at the “wrong” C-atom of fumarate. Regarding the last step of the model, the transfer of an H atom back from Gly829 to the thiyl group of Cys493 (step 5), our model suggests that this does not happen as expected, because it is too endergonic to be overcome easily. We, therefore, suggest that the completion of the reaction needs to be coupled with the eventual release of the bound product benzylsuccinate.

In addition to the movements of the amino acid side chains actively involved in the reaction, we have also observed the requirement of other residues of the active site to move into slightly different positions with almost every step of the reaction cycle. The most apparent of these cases relates to the associated movement of the active site residue Tyr197, which is necessary to enable the HAT reaction between the thiyl radical and the methyl group of toluene (step 2) and probably acts to fine-tune this step of the reaction cycle, e.g. as a gating or modulating device.

### Enantioselectivity

The obtained potential energy surface for the pro*S*-pathway at first glance contradicts our previous results obtained from the significantly more simplistic QM-only analysis on the cluster model and docking-based binding energy estimation^44^. We have no longer observed any statistically significant preference for binding fumarate in the pro*R* manner over pro*S* in this study, regardless of its protonation mode. The obtained reaction energy profiles no longer present a clear switch in C-C bond formation preference at the proximal or distal carbon atom of the fumarate, depending on the fumarate binding mode. However, the analysis of MD simulations for the models approximating the reaction intermediate I2 indicated a difference in the probability of an attack on the distal vs proximal atom of the fumarate bound in the pro*S* or pro*R* position. The increased chance of forming a C-C bond at the proximal atom for the pro*S*-bound fumarate combined with a slower rate favors the *R*-enantioselectivity of the reaction.

However, the decisive explanation of BSS enantioselectivity was provided by analysing the overall rate of the internal processes proceeding along the pro*R* or pro*S* pathways, indicating that the formation of the *R*-benzylsuccinyl radical is 115-fold faster than of the *S*-enantiomer. Those two factors together i.e. shift in the probability of the preferential near attack conformation in function of the fumarate binding mode and over 100-fold slower kinetics of *S*-benzylsuccinate formation along the kinetically preferential pathway a, in our view explain the high enantioselectivity of the benzylsuccinate synthase.

### Kinetic isotope effect

The results, obtained from two independent experiments, stand in line with previous observations of Li & Marsh conducted for BSS from *Thauera aromatica* T1^61^. The observed 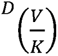 D(VJ from competitive method (3.7±0.3) was higher than that recorded for V_max_ (2.1 ± 0.1) similarly as in the Li & Marsh report (2.9±0.1 and 1.7±0.2, respectively). The KIE observed in the direct experiment was two-fold lower than in our previous experiments^62^ which can be explained by the improved analytical method used in this study. The value of the KIE predicted with the theoretical method (2.6) is in reasonably good agreement with both experimental studies. Nevertheless, we think that such a good agreement indicates that the obtained reaction pathway and its barriers deliver reasonable prediction capabilities. Furthermore, based on the difference of values between direct and competitive KIE we can suspect a strong influence of the different enzyme affinity substrate yielding preferential binding of toluene over its substituted d_8_-isotopologue.

### HDX

Finally, we propose the mechanism along which the observed low-level HDX effects occur during the reaction with d_8_-toluene or during the incubation of benzylsuccinate in D_2_O. Although the very sensitive nature of BSS and the low product yields prevented a direct NMR-based measurement of the HDX rate at Cys493 we gained some insight into that process using kinetic analysis combined with experiments. Taking several assumptions into consideration we used measured rates of d_1_ and d_2_-benzylsuccinate formation and rate equation based on QM:MM-calculations to estimate the potential rate of H/D exchange in the enzyme. The estimated microkinetic rates of the H/D and D/H exchanges were in the range of 0.3-0.5 s^−1^ which is comparable with the calculated rate of reaction (0.52 s^−1^). As a result, we can expect that the H/D exchange process is relatively slow and will influence the observed rate, as demonstrated by D/H exchange during the reaction in H_2_O with d_8_-toluene^48^. These results support the occurrence of partial reversibility of the BSS mechanism and indicate small deviations from enantioselectivity during the quenching of the benzylsuccinyl radical. Finally, we propose that the formation of d_2_-benzylsuccinate can only be achieved by the kinetically more challenging transfer of deuteron to position b in the benzylsuccinyl radical, followed by the less kinetically demanding removal of the H atom from position a, which is additionally supported by an inverse secondary kinetic isotope effect (0.7). This effect is very similar to the recently analysed HDX at Gly radical in our previous report^48^.

### Summary

In this study, we describe most of the catalytic aspects of the addition of fumarate to toluene catalyzed by benzylsuccinate synthase by theoretical modelling. It represents the first time that all structural and electronic aspects of the enzyme influencing the complex multistep reaction were taken into consideration for MD and QM:MM analysis. The obtained model did not only provide a qualitative understanding of the catalytic process but was robust enough to yield good estimates of experimentally obtained values such as kinetic isotope effects, enantioselectivity or rate ratios of forming labelled products in HDX setups. However, we are aware that the model used in the study represents only one of many potential conformations and does not take into account the dynamic behaviour of the enzyme. This type of calculation is still beyond reach due to the nature of the radical reaction and the complexity of the process.

## Supporting information

PDB structures

PDB structures

KIE

## Acknowledgements

The authors acknowledge the financial support provided by Deutsche Forschungsgemeinschaft/National Science Center Poland under Beethoven Life grant He2190/13-1 / 2018/31/F/NZ1/01856 as well as Polish high-performance computing infrastructure PLGrid (HPC Center: ACK Cyfronet AGH) for providing computer facilities and support within computational grant no. PLG/2023/016888, PLG/2022/016024 and PLG/2021/015218.

## Supplementary Information

The supplementary information contains details of methods such as: derivation of all kinetic equations, reaction schemes and analytical procedures as well as results such as RMSD of all MD simulations, additional figures describing mechanism, results of MD analysis, tables with all data from QM calculations (energies and vibration corrections), tables with kinetic constants as well as all PDB structures of stationary points used in the study.

