## Supplementary material for "Unravelling the Enantioselective Mechanism of Benzylsuccinate Synthase: Insights into Anaerobic Hydrocarbon Degradation Through Multiscale Modelling and Kinetics": SI

### Index

### METHODS

#### LC-MS/MS methods

Table S1 Parameters of jet-stream ESI ion source used in the LC-MS/MS analysis.

| Parameter | Value (+) | Value (-) |
|---|---|---|
| **Gas Temp [°C]** | 300 | 300 |
| **Gas Flow [L/min]** | 10 | 10 |
| **Nebulizer [psi]** | 45 | 45 |
| **Sheath Gas Heater [°C]** | 300 | 300 |
| **Sheath Gas Flow [L/min]** | 10 | 10 |
| **Capillary [V]** | 3500 | 3500 |
| **V Charging [V]** | 500 | 1000 |

Table S2. Parameters of MRM method for the analysis of benzylsuccinate, $d^7$-benzylsuccinate and $d^8$-benzylsuccinate.

| Compound | Precursor ion [m/z] | Product ion [m/z] | Dwell [ms] | Fragmentor [V] | Collision energy [v] | Cell accelerator [V] | Polarity |
|---|---|---|---|---|---|---|---|
| **benzylsuccinate** | 207.1 | 163.0 | 200 | 107 | 10 | 4 | negative |
| **$d_8$-benzylsuccinate** | 215.0 | 171.0 | 200 | 107 | 10 | 4 | negative |
| **$d_7$-benzylsuccinate** | 214.0 | 170.0 | 200 | 107 | 10 | 4 | negative |

Table S3. Parameters for analysis of product ions of benzylsuccinate 207, 209, 209 m/z signals (respectively, $[M-H]^-$, $[M+1-H]^-$ and $[M+2-H]^-$).

| Precursor ion [m/z] | MS2 from | MS2 to | Scan time [ms] | Fragmentor [V] | Collision energy [v] | Cell accelerator [V] | Polarity |
|---|---|---|---|---|---|---|---|
| 207.0 | 150.0 | 212.0 | 500 | 107 | 10 | 4 | negative |
| 208.0 | 150.0 | 212.0 | 500 | 107 | 10 | 4 | negative |
| 209.0 | 150.0 | 212.0 | 500 | 107 | 10 | 4 | negative |

#### Calibration curves for LC-MS/MS and LC-DAD benzylsuccinate quantitation

A) B)

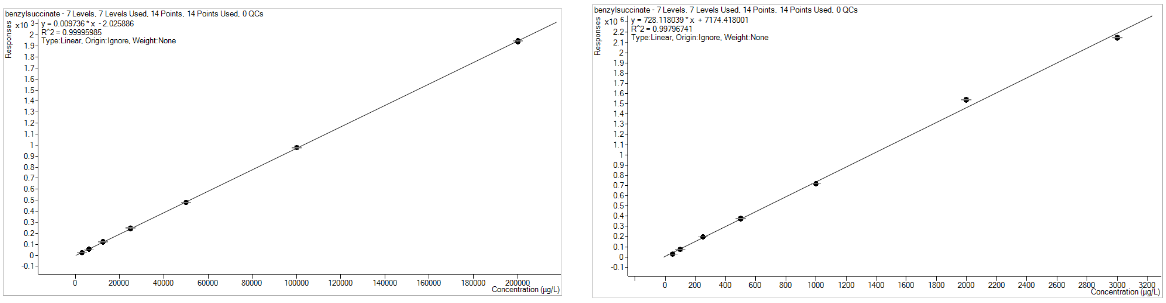

Figure S1. Calibration curves for LC-MS/MS quantitation of A) LC-DAD quantitation of benzylsuccinate; B) benzylsuccinate in SIM mode. Concentrations are given in µg/L.

#### Derivation of the rate equations

##### Whole reaction:

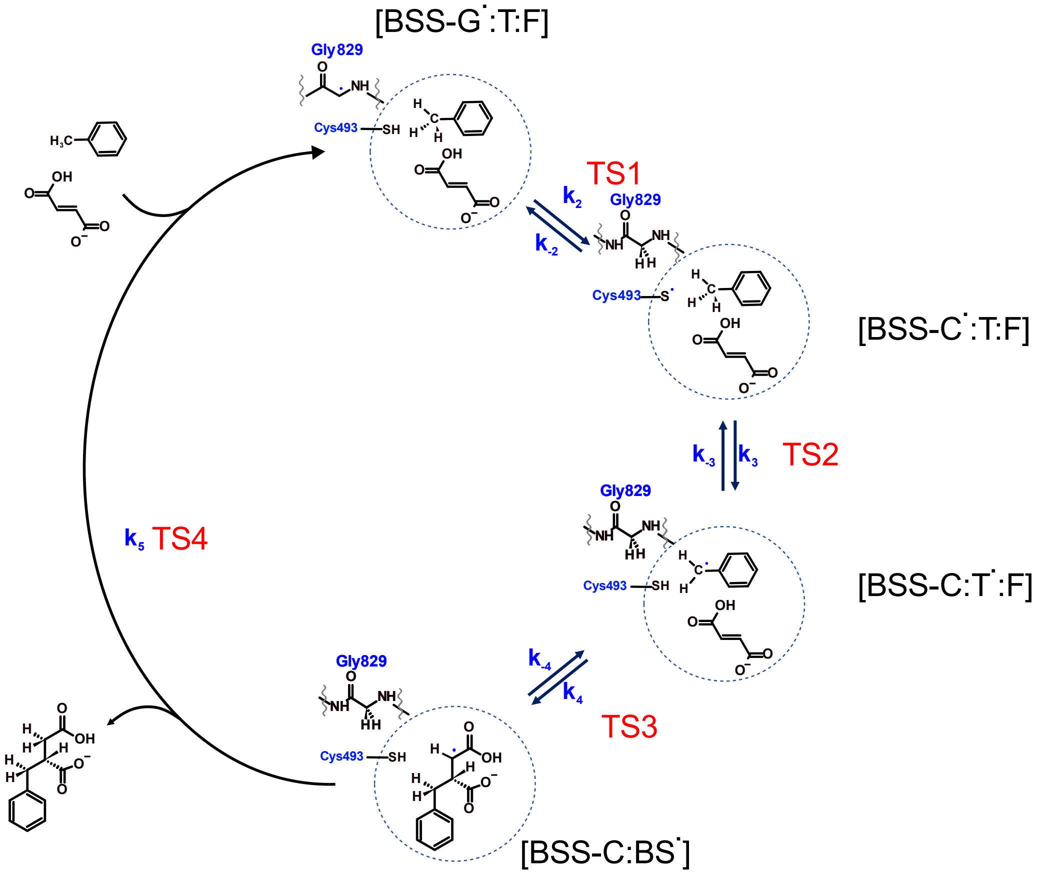

Scheme S1. The schematic representation of the whole reaction leading to the formation of benzylsuccinate assuming the irreversibility of the benzylsuccinate radical quenching step and no kinetic limitation from substrates binding/product release steps. [BSS-G·:T:F] – BSS with glycyl radical in complex with toluene and fumarate, [BSS-C·:T:F] – BSS with cysteinyl radical in complex with toluene and fumarate, [BSS-C:T·:F] – BSS with cysteine in complex with benzyl radical and fumarate, [BSS-C:BS·] – BSS with cysteine in complex with benzylsuccinyl radical

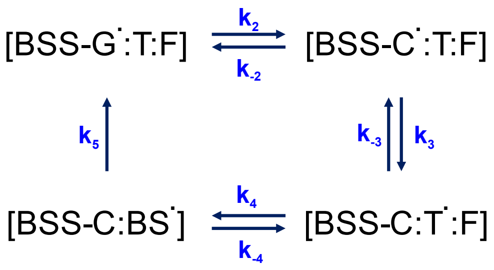

**I**

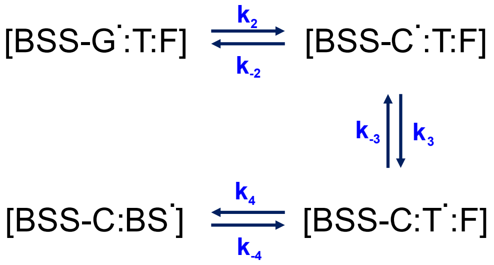

**II**

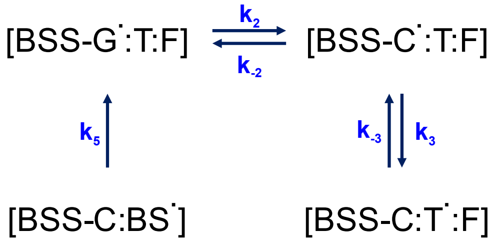

**III**

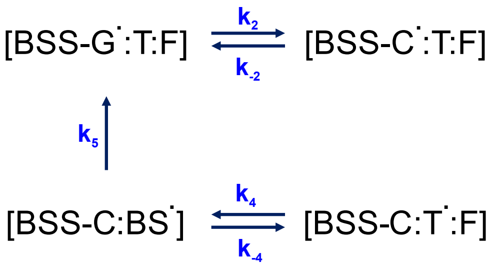

**IV**

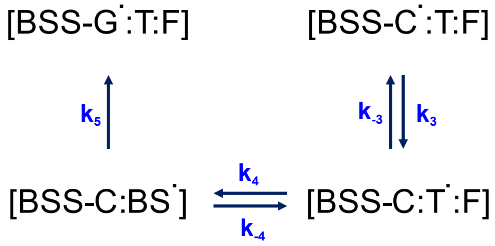

From the diagram describing internal enzyme reactions (at substrate saturation conditions) with irreversible radical quenching step ($k_5$) we construct four different subdiagrams (I-IV) without one transition. We construct equations describing the relative concentration of each enzyme species by multiplying microkinetic constants leading to that species but not involving it. We add terms for each diagram and introduce it to the nominator of the equation.

Eq. S1

$$\frac{[B - G^{\cdot}:T:F]}{E_T} = \frac{k_{-2}k_{-3}k_{-4} + k_{-2}k_{-3}k_5 + k_{-2}k_4k_5 + k_3k_4k_5}{\Delta}$$

Eq. S2

$$\frac{[B - C^{\cdot}:T:F]}{E_T} = \frac{k_2k_{-3}k_{-4} + k_2k_{-3}k_5 + k_2k_4k_5 + 0 \cdot k_{-3}k_{-4}}{\Delta}$$

Eq. S3

$$\frac{[B - C:T^{\cdot}:F]}{E_T} = \frac{k_2k_3k_{-4} + k_2k_3k_5 + 0 \cdot k_{-4} + 0 \cdot k_3k_{-4}}{\Delta}$$

Eq. S4

$$\frac{[B-C:BS^{\cdot}]}{E_T}=\frac{k_2k_3k_4+0\cdot k_4+0\cdot k_3k_4}{\Delta}$$

As at the substrate saturation conditions, the total enzyme concentration equals the sum of the individual species we can show that Δ is a sum of all nominators of all four equations.

Eq. S5

$$\frac{[B-G^{\cdot}:T:F]}{E_T}+\frac{[B-C^{\cdot}:T:F]}{E_T}+\frac{[B-C:T^{\cdot}:F]}{E_T}+\frac{[B-C:BS^{\cdot}]}{E_T}=1$$

Eq. S6

$$\Delta= k_{-2}k_{-3}k_{-4}+k_{-2}k_{-3}k_5+k_{-2}k_4k_5+k_3k_4k_5+k_2k_{-3}k_{-4}+k_2k_{-3}k_5+k_2k_4k_5+0\cdot k_{-3}k_{-4}+k_2k_3k_{-4}+k_2k_3k_5+0\cdot k_{-4}+0\cdot k_3k_{-4}+k_2k_3k_4+0\cdot k_4+0\cdot k_3k_4=k_{-2}k_{-3}k_{-4}+k_{-2}k_{-3}k_5+k_{-2}k_4k_5+k_3k_4k_5+k_2k_{-3}k_{-4}+k_2k_{-3}k_5+k_2k_4k_5+k_2k_3k_{-4}+k_2k_3k_5+k_2k_3k_4$$

The observed in the experiment reaction velocity is the rate of product formation. In that case (assuming a low concentration of the product) this will depend on the concentration of the enzyme before the irreversible step:

Eq. S7

$$V=k_5[BSS-C:BS^{\cdot}]$$

Combining Eq 7 with Eq. 4 we get:

Eq. S8

$$V=\frac{k_2k_3k_4k_5E_T}{k_{-2}k_{-3}k_{-4}+k_{-2}k_{-3}k_5+k_{-2}k_4k_5+k_3k_4k_5+k_2k_{-3}k_{-4}+k_2k_{-3}k_5+k_2k_4k_5+k_2k_3k_{-4}+k_2k_3k_5+k_2k_3k_4}$$

The $V/E_T$ value can be calculated using the kinetic constants derived from pro*R* or pro*S* reaction pathway thus enabling estimation of the relative rates leading to *R* or *S*-benzylsuccinate.

#### HDX leading to $d_1$-benzylsuccinate

**Assumption:**

Upon formation of $d_1$- benzylsuccinate (BS)it is released to the solvent and due to its very low concentration with respect to the unsubstituted BS the reaction is irreversible. The BS concentration is so high that the enzyme saturated by the product and binding of the product does not influence the overall kinetics.

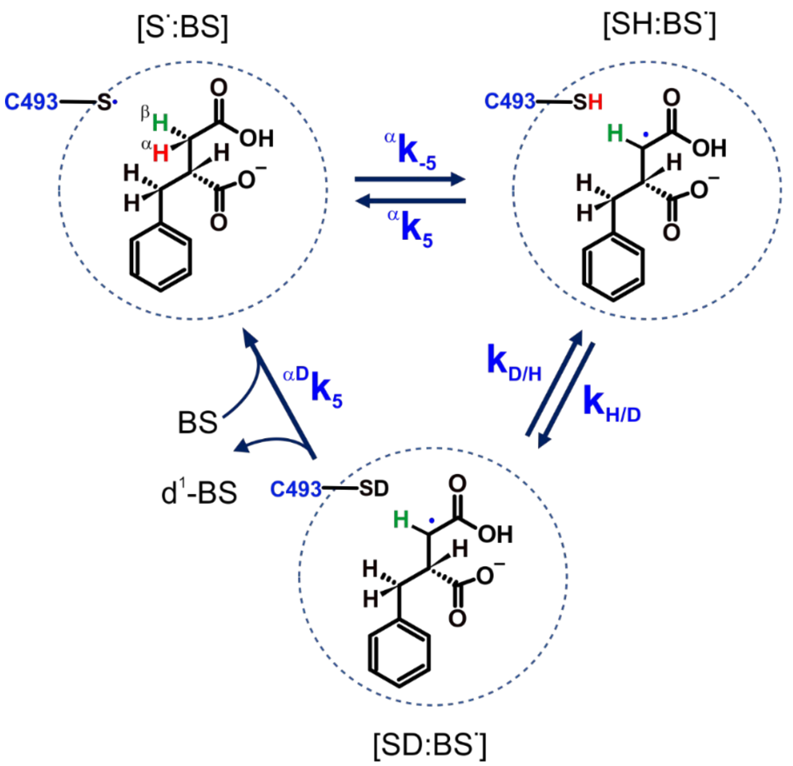

Scheme S2. The schematic representation of H/D exchange in the BSS:BS complex during incubation in $D_2O$ leading to the production of $d_1$-benzylsuccinate assuming irreversibility of the D transfer to benzylsuccinyl radical and no kinetic limitation by product binding/release. [S·:BS] – BSS with radical Cys in complex with benzylsuccinate, [SH:BS·] – BSS with Cys-SH in complex with benzylsuccinyl radical, [SD:BS·] – BSS with Cys-SD in complex with benzylsuccinyl radical

Eq. S9

$$\frac{[SD{:}BS^{\cdot}]}{E_T} = \frac{0\cdot k_{H/D} + 0\cdot {}^{\alpha}k_5 + {}^{\alpha}k_{-5}k_{H/D}}{\Delta} = \frac{{}^{\alpha}k_{-5}k_{H/D}}{\Delta}$$

Eq. S10

$$\frac{[S^{\cdot}{:}BS]}{E_T} = \frac{k_{H/D}{}^{\alpha D}k_5 + {}^{\alpha D}k_5{}^{\alpha}k_5 + k_{D/H}{}^{\alpha}k_5}{\Delta}$$

Eq. S11

$$\frac{[SH{:}BS^{\cdot}]}{E_T} = \frac{0\cdot k_{D/H} + {}^{\alpha D}k_5{}^{\alpha}k_{-5} + k_{D/H}{}^{\alpha}k_{-5}}{\Delta}$$

Eq. S12

$$\Delta = 0\cdot k_{H/D} + 0\cdot {}^{\alpha}k_5 + {}^{\alpha}k_{-5}k_{H/D} + k_{H/D}{}^{\alpha D}k_5 + {}^{\alpha D}k_5{}^{\alpha}k_5 + k_{D/H}{}^{\alpha}k_5 + 0\cdot k_{\frac{D}{H}} + {}^{\alpha D}k_5{}^{\alpha}k_{-5} + k_{\frac{D}{H}}{}^{\alpha}k_{-5}$$
$$== {}^{\alpha}k_{-5}k_{H/D} + k_{H/D}{}^{\alpha D}k_5 + {}^{\alpha D}k_5{}^{\alpha}k_5 + k_{D/H}{}^{\alpha}k_5 + {}^{\alpha D}k_5{}^{\alpha}k_{-5} + k_{D/H}{}^{\alpha}k_{-5}$$

Eq. S13

$$V^{d1} = {}^{\alpha D}k_5[SD{:}BS^{\cdot}]$$

Eq. S14

$$V^{d1} = \frac{{}^{\alpha D}k_5\,{}^{\alpha}k_{-5}k_{H/D}[E_T]}{{}^{\alpha}k_{-5}k_{H/D} + k_{H/D}\,{}^{\alpha D}k_5 + {}^{\alpha D}k_5\,{}^{\alpha}k_5 + k_{D/H}\,{}^{\alpha}k_5 + {}^{\alpha D}k_5\,{}^{\alpha}k_{-5} + k_{D/H}\,{}^{\alpha}k_{-5}}$$

##### HDX leading to $d_2$-benzylsuccinate – alpha pathway

**Assumption:**

Upon formation of $d_2$- benzylsuccinate it is released to the solvent and due to its very low concentration with respect to the unsubstituted benzylsuccinate the reaction is irreversible. The benzylsuccinate concentration is so high that the enzyme saturated by the product and binding of the product does not influence the overall kinetics.

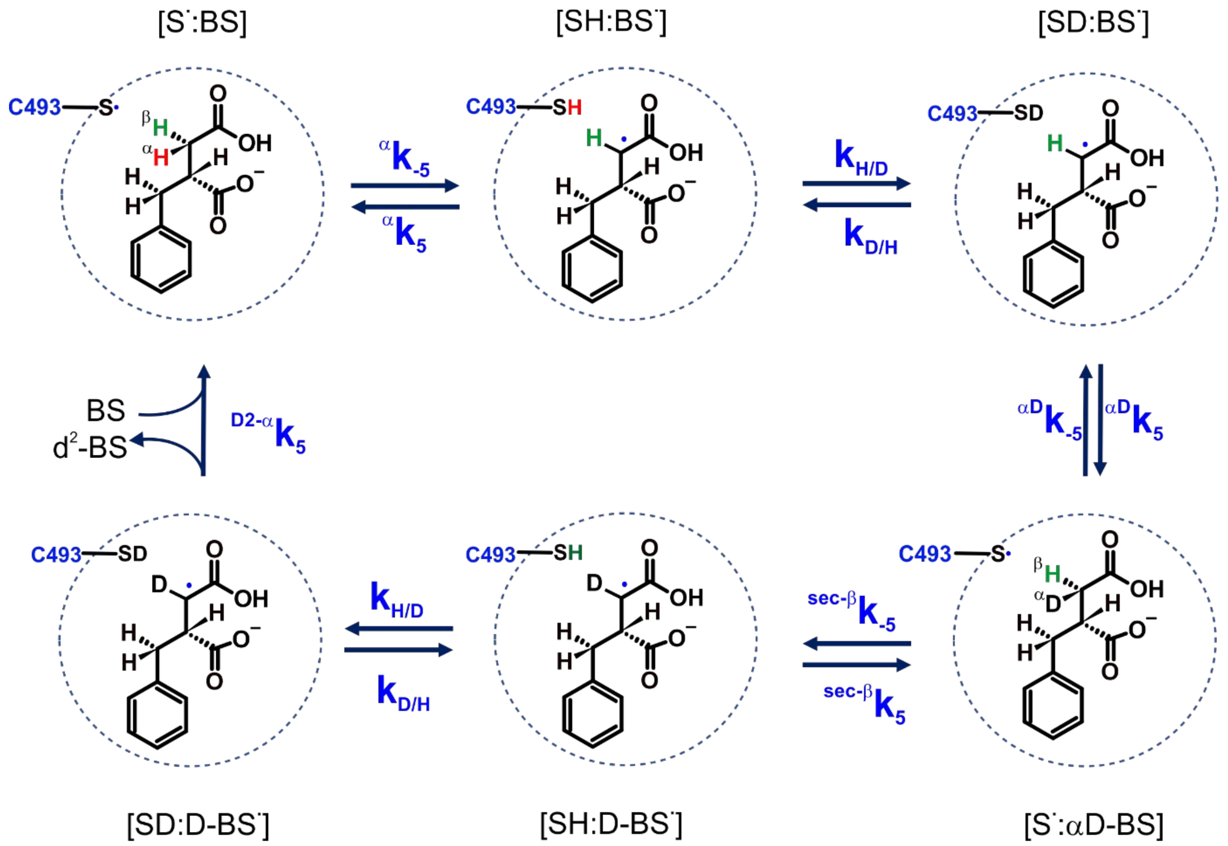

Scheme S3. The schematic representation of α pathway for H/D exchange in the BSS:BS complex during incubation in $D_2O$ leading to the production of $d_2$-benzylsuccinate assuming irreversibility of the D transfer to benzylsuccinyl radical and no kinetic limitation by product binding/release. [S˙:BS] – BSS with radical Cys in complex with benzylsuccinate, [SH:BS˙] – BSS with Cys-SH in complex with benzylsuccinyl radical, [SD:BS˙] – BSS with Cys-SD in complex with benzylsuccinyl radical, [S˙:αD-BS] BSS with radical Cys in complex with α-$d_1$-

benzylsuccinate, [SH:D-BS·] - BSS with Cys-SH in complex with $d_1$-benzylsuccinyl radical, [SD:D-BS·] - BSS with Cys-SD in complex with $d_1$-benzylsuccinyl radical

Eq. S15

$$V = {}^{D2\alpha}k_5[SD:D - BS^{\cdot}]$$

Eq. S16

$$\frac{[SD:D{-}BS^{\cdot}]}{E_T} = \frac{0 \cdot k_{\frac{H}{D}}^2 {}^{\alpha D}k_5 {}^{sec\beta}k_{-5} + {}^{\alpha D}k_5 {}^{sec\beta}k_{-5} k_{\frac{H}{D}} {}^{\alpha}k_5 \cdot 0 + {}^{sec\beta}k_{-5} k_{\frac{H}{D}} k_{\frac{D}{H}} {}^{\alpha}k_5 \cdot 0 + k_{\frac{H}{D}} {}^{\alpha D}k_{-5} k_{\frac{D}{H}} {}^{\alpha}k_5 \cdot 0 + {}^{\alpha}k_{-5} k_{\frac{H}{D}}^2 {}^{\alpha D}k_5 {}^{sec\beta}k_{-5}}{\Delta} = \frac{{}^{\alpha}k_{-5} k_{H/D}^2 {}^{\alpha D}k_5 {}^{sec\beta}k_{-5}}{\Delta}$$

Eq. S17

$$\frac{[S^{\cdot}:BS]}{E_T} = \frac{k_{H/D}^2 {}^{\alpha D}k_5 {}^{sec\beta}k_{-5} {}^{D2\alpha}k_5 + {}^{\alpha}k_5 {}^{\alpha D}k_5 {}^{sec\beta}k_{-5} k_{H/D} {}^{D2\alpha}k_5 + k_{D/H} {}^{\alpha}k_5 {}^{sec\beta}k_{-5} k_{H/D} {}^{D2\alpha}k_5}{\Delta} + \frac{{}^{\alpha D}k_{-5} k_{D/H} {}^{\alpha}k_5 k_{H/D} {}^{D2\alpha}k_5 + {}^{sec\beta}k_5 {}^{\alpha D}k_{-5} k_{D/H} {}^{\alpha}k_5 {}^{D2\alpha}k_5 + k_{D/H}^2 {}^{sec\beta}k_5 {}^{\alpha D}k_{-5} {}^{\alpha}k_5}{\Delta}$$

Eq. S18

$$\frac{[SH:BS^{\cdot}]}{E_T} = \frac{k_{D/H}^2 {}^{sec\beta}k_5 {}^{\alpha D}k_{-5} \cdot 0 + {}^{\alpha D}k_5 {}^{sec\beta}k_{-5} k_{H/D} {}^{D2\alpha}k_5 {}^{\alpha}k_{-5} + k_{D/H} {}^{sec\beta}k_{-5} k_{H/D} {}^{D2\alpha}k_5 {}^{\alpha}k_{-5}}{\Delta} + \frac{{}^{\alpha D}k_{-5} k_{D/H} k_{H/D} {}^{D2\alpha}k_5 {}^{\alpha}k_{-5} + {}^{sec\beta}k_5 {}^{\alpha D}k_{-5} k_{D/H} {}^{D2\alpha}k_5 {}^{\alpha}k_{-5} + {}^{\alpha}k_{-5} k_{D/H}^2 {}^{sec\beta}k_5 {}^{\alpha D}k_{-5}}{\Delta}$$

Eq. S19

$$\frac{[SD:BS^{\cdot}]}{E_T} = \frac{k_{H/D} \cdot 0 \cdot k_{D/H} {}^{sec\beta}k_5 {}^{\alpha D}k_{-5} + {}^{\alpha}k_5 \cdot 0 \cdot k_{D/H} {}^{sec\beta}k_5 {}^{\alpha D}k_{-5} + {}^{sec\beta}k_{-5} k_{H/D}^2 {}^{D2\alpha}k_5 {}^{\alpha}k_{-5}}{\Delta} + \frac{{}^{\alpha D}k_{-5} k_{H/D}^2 {}^{D2\alpha}k_5 {}^{\alpha}k_{-5} + {}^{sec\beta}k_5 {}^{\alpha D}k_{-5} {}^{D2\alpha}k_5 {}^{\alpha}k_{-5} k_{H/D} + {}^{\alpha}k_{-5} k_{H/D} k_{D/H} {}^{sec\beta}k_5 {}^{\alpha D}k_{-5}}{\Delta}$$

$$\frac{[S^{\cdot}:\alpha DBS]}{E_T} = \frac{k_{H/D} {}^{\alpha D}k_5 \cdot 0 \cdot k_{D/H} {}^{sec\beta}k_5 + {}^{\alpha D}k_5 \cdot 0 \cdot k_{D/H} {}^{sec\beta}k_5 + k_{D/H}^2 {}^{\alpha}k_5 \cdot 0 \cdot {}^{sec\beta}k_5}{\Delta} + \frac{k_{H/D}^2 {}^{D2\alpha}k_5 {}^{\alpha}k_{-5} {}^{\alpha D}k_5 + {}^{sec\beta}k_5 {}^{D2\alpha}k_5 {}^{\alpha}k_{-5} k_{H/D} {}^{\alpha D}k_5 + {}^{\alpha}k_{-5} k_{H/D} {}^{\alpha D}k_5 k_{D/H} {}^{sec\beta}k_5}{\Delta}$$

$$= \frac{k_{H/D}^2 {}^{D2\alpha}k_5 {}^{\alpha}k_{-5} {}^{\alpha D}k_5 + {}^{sec\beta}k_5 {}^{D2\alpha}k_5 {}^{\alpha}k_{-5} k_{H/D} {}^{\alpha D}k_5 + {}^{\alpha}k_{-5} k_{H/D} {}^{\alpha D}k_5 k_{D/H} {}^{sec\beta}k_5}{\Delta}$$

$$\frac{[SH:DBS^{\cdot}]}{E_T}=\frac{0\cdot k_{H/D}{}^{\alpha D}k_5{}^{sec\beta}k_{-5}k_{D/H}+{}^{\alpha D}k_5{}^{sec\beta}k_{-5}{}^{\alpha}k_5\cdot 0\cdot k_{D/H}+k_{D/H}{}^{\alpha}k_5\cdot 0\cdot{}^{sec\beta}k_{-5}k_{D/H}}{\Delta}+$$

$$\frac{{}^{\alpha D}k_{-5}k_{D/H}{}^{\alpha}k_5\cdot 0\cdot k_{D/H}+{}^{D2\alpha}k_5{}^{\alpha}k_{-5}k_{H/D}{}^{\alpha D}k_5{}^{sec\beta}k_{-5}+k_{D/H}{}^{\alpha}k_{-5}k_{H/D}{}^{\alpha D}k_5{}^{sec\beta}k_{-5}}{\Delta}$$

$$=\frac{{}^{D2\alpha}k_5{}^{\alpha}k_{-5}k_{H/D}{}^{\alpha D}k_5{}^{sec\beta}k_{-5}+k_{D/H}{}^{\alpha}k_{-5}k_{H/D}{}^{\alpha D}k_5{}^{sec\beta}k_{-5}}{\Delta}$$

Eq. S20

$$\Delta={}^{\alpha}k_{-5}k_{H/D}^2{}^{\alpha D}k_5{}^{sec\beta}k_{-5}+k_{H/D}^2{}^{\alpha D}k_5{}^{sec\beta}k_{-5}{}^{D2\alpha}k_5+{}^{\alpha}k_5{}^{\alpha D}k_5{}^{sec\beta}k_{-5}k_{H/D}{}^{D2\alpha}k_5+$$
$$k_{D/H}{}^{\alpha}k_5{}^{sec\beta}k_{-5}k_{H/D}{}^{D2\alpha}k_5+{}^{\alpha D}k_{-5}k_{D/H}{}^{\alpha}k_5k_{H/D}{}^{D2\alpha}k_5+{}^{sec\beta}k_5{}^{\alpha D}k_{-5}k_{D/H}{}^{\alpha}k_5{}^{D2\alpha}k_5+$$
$$k_{D/H}^2{}^{sec\beta}k_5{}^{\alpha D}k_{-5}{}^{\alpha}k_5+{}^{\alpha D}k_5{}^{sec\beta}k_{-5}k_{H/D}{}^{D2\alpha}k_5{}^{\alpha}k_{-5}+$$
$$k_{D/H}{}^{sec\beta}k_{-5}k_{H/D}{}^{D2\alpha}k_5{}^{\alpha}k_{-5}+{}^{\alpha D}k_{-5}k_{D/H}k_{H/D}{}^{D2\alpha}k_5{}^{\alpha}k_{-5}+{}^{sec\beta}k_5{}^{\alpha D}k_{-5}k_{D/H}{}^{D2\alpha}k_5{}^{\alpha}k_{-5}+$$
$${}^{\alpha}k_{-5}k_{D/H}^2{}^{sec\beta}k_5{}^{\alpha D}k_{-5}+{}^{sec\beta}k_{-5}k_{H/D}^2{}^{D2\alpha}k_5{}^{\alpha}k_{-5}+{}^{\alpha D}k_{-5}k_{H/D}^2{}^{D2\alpha}k_5{}^{\alpha}k_{-5}+$$
$${}^{sec\beta}k_5{}^{\alpha D}k_{-5}{}^{D2\alpha}k_5{}^{\alpha}k_{-5}k_{H/D}+{}^{\alpha}k_{-5}k_{H/D}k_{D/H}{}^{sec\beta}k_5{}^{\alpha D}k_{-5}+k_{H/D}^2{}^{D2\alpha}k_5{}^{\alpha}k_{-5}{}^{\alpha D}k_5+$$
$${}^{sec\beta}k_5{}^{D2\alpha}k_5{}^{\alpha}k_{-5}k_{H/D}{}^{\alpha D}k_5+{}^{\alpha}k_{-5}k_{H/D}{}^{\alpha D}k_5k_{D/H}{}^{sec\beta}k_5+{}^{D2\alpha}k_5{}^{\alpha}k_{-5}k_{H/D}{}^{\alpha D}k_5{}^{sec\beta}k_{-5}+$$
$$k_{D/H}{}^{\alpha}k_{-5}k_{H/D}{}^{\alpha D}k_5{}^{sec\beta}k_{-5}$$

Eq. S21

$$V^{d2-\alpha}=\frac{{}^{\alpha}k_{-5}k_{H/D}^2{}^{\alpha D}k_5{}^{sec\beta}k_{-5}{}^{D2\alpha}k_5}{\Delta}E_T$$

$$V={}^{D2\alpha}k_5[SD:D-BS^{\cdot}]$$

Eq. S22

$$\frac{[SD:D-BS^{\cdot}]}{E_T}==\frac{k_{H/D}^2{}^{\alpha D}k_5{}^{sec\beta}k_{-5}+{}^{\alpha D}k_5{}^{sec\beta}k_{-5}k_{H/D}+{}^{sec\beta}k_{-5}k_{H/D}+k_{H/D}+{}^{\alpha}k_{-5}k_{H/D}^2{}^{\alpha D}k_5{}^{sec\beta}k_{-5}}{\Delta}$$

Eq. S23

$$\frac{[S^{\cdot}:BS]}{E_T}=\frac{k_{H/D}^2{}^{\alpha D}k_5{}^{sec\beta}k_{-5}{}^{D2\alpha}k_5+{}^{\alpha}k_5{}^{\alpha D}k_5{}^{sec\beta}k_{-5}k_{H/D}{}^{D2\alpha}k_5+k_{D/H}{}^{\alpha}k_5{}^{sec\beta}k_{-5}k_{H/D}{}^{D2\alpha}k_5}{\Delta}+$$
$$+\frac{{}^{\alpha D}k_{-5}k_{D/H}{}^{\alpha}k_5k_{H/D}{}^{D2\alpha}k_5+{}^{sec\beta}k_5{}^{\alpha D}k_{-5}k_{D/H}{}^{\alpha}k_5{}^{D2\alpha}k_5+k_{D/H}^2{}^{sec\beta}k_5{}^{\alpha D}k_{-5}{}^{\alpha}k_5}{\Delta}$$

Eq. S24

$$\frac{[SH:BS^{\cdot}]}{E_T} = \frac{k_{D/H}^2{}^{sec\beta}k_5{}^{\alpha D}k_{-5} + {}^{\alpha D}k_5{}^{sec\beta}k_{-5}k_{H/D}{}^{D2\alpha}k_5{}^{\alpha}k_{-5} + k_{D/H}{}^{sec\beta}k_{-5}k_{H/D}{}^{D2\alpha}k_5{}^{\alpha}k_{-5}}{\Delta} + \frac{{}^{\alpha D}k_{-5}k_{D/H}k_{H/D}{}^{D2\alpha}k_5{}^{\alpha}k_{-5} + {}^{sec\beta}k_5{}^{\alpha D}k_{-5}k_{D/H}{}^{D2\alpha}k_5{}^{\alpha}k_{-5} + {}^{\alpha}k_{-5}k_{D/H}^2{}^{sec\beta}k_5{}^{\alpha D}k_{-5}}{\Delta}$$

Eq. S25

$$\frac{[SD:BS^{\cdot}]}{E_T} = \frac{k_{H/D}k_{D/H}{}^{sec\beta}k_5{}^{\alpha D}k_{-5} + k_{D/H}{}^{sec\beta}k_5{}^{\alpha D}k_{-5} + {}^{sec\beta}k_{-5}k_{H/D}^2{}^{D2\alpha}k_5{}^{\alpha}k_{-5}}{\Delta} + \frac{{}^{\alpha D}k_{-5}k_{H/D}^2{}^{D2\alpha}k_5{}^{\alpha}k_{-5} + {}^{sec\beta}k_5{}^{\alpha D}k_{-5}{}^{D2\alpha}k_5{}^{\alpha}k_{-5}k_{H/D} + {}^{\alpha}k_{-5}k_{H/D}k_{D/H}{}^{sec\beta}k_5{}^{\alpha D}k_{-5}}{\Delta}$$

$$\frac{[S^{\cdot}:\alpha DBS]}{E_T} = \frac{k_{H/D}{}^{\alpha D}k_5k_{D/H}{}^{sec\beta}k_5 + {}^{\alpha D}k_5k_{D/H}{}^{sec\beta}k_5 + k_{D/H}{}^{sec\beta}k_5}{\Delta} + \frac{k_{H/D}^2{}^{D2\alpha}k_5{}^{\alpha}k_{-5}{}^{\alpha D}k_5 + {}^{sec\beta}k_5{}^{D2\alpha}k_5{}^{\alpha}k_{-5}k_{H/D}{}^{\alpha D}k_5 + {}^{\alpha}k_{-5}k_{H/D}{}^{\alpha D}k_5k_{D/H}{}^{sec\beta}k_5}{\Delta}$$

$$\frac{[SH:DBS^{\cdot}]}{E_T} = \frac{k_{H/D}{}^{\alpha D}k_5{}^{sec\beta}k_{-5}k_{D/H} + {}^{\alpha D}k_5{}^{sec\beta}k_{-5}k_{D/H} + {}^{sec\beta}k_{-5}k_{D/H}}{\Delta} + \frac{k_{D/H} + {}^{D2\alpha}k_5{}^{\alpha}k_{-5}k_{H/D}{}^{\alpha D}k_5{}^{sec\beta}k_{-5} + k_{D/H}{}^{\alpha}k_{-5}k_{H/D}{}^{\alpha D}k_5{}^{sec\beta}k_{-5}}{\Delta}$$

Eq. S26

$$\Delta= k_{H/D}^{2}\,{}^{\alpha D}k_5\,{}^{sec\beta}k_{-5} + {}^{\alpha D}k_5\,{}^{sec\beta}k_{-5}k_{H/D} + {}^{sec\beta}k_{-5}k_{H/D} + k_{H/D} + {}^{\alpha}k_{-5}k_{H/D}^{2}\,{}^{\alpha D}k_5\,{}^{sec\beta}k_{-5} + k_{H/D}^{2}\,{}^{\alpha D}k_5\,{}^{sec\beta}k_{-5}\,{}^{D2\alpha}k_5 + {}^{\alpha}k_5\,{}^{\alpha D}k_5\,{}^{sec\beta}k_{-5}k_{H/D}\,{}^{D2\alpha}k_5 + k_{D/H}\,{}^{\alpha}k_5\,{}^{sec\beta}k_{-5}k_{H/D}\,{}^{D2\alpha}k_5 + {}^{\alpha D}k_{-5}k_{D/H}\,{}^{\alpha}k_5k_{H/D}\,{}^{D2\alpha}k_5 + {}^{sec\beta}k_5\,{}^{\alpha D}k_{-5}k_{D/H}\,{}^{\alpha}k_5\,{}^{D2\alpha}k_5 + k_{D/H}^{2}\,{}^{sec\beta}k_5\,{}^{\alpha D}k_{-5}\,{}^{\alpha}k_5 + k_{D/H}^{2}\,{}^{sec\beta}k_5\,{}^{\alpha D}k_{-5} + {}^{\alpha D}k_5\,{}^{sec\beta}k_{-5}k_{H/D}\,{}^{D2\alpha}k_5\,{}^{\alpha}k_{-5} + k_{D/H}\,{}^{sec\beta}k_{-5}k_{H/D}\,{}^{D2\alpha}k_5\,{}^{\alpha}k_{-5} + {}^{\alpha D}k_{-5}k_{D/H}k_{H/D}\,{}^{D2\alpha}k_5\,{}^{\alpha}k_{-5} + {}^{sec\beta}k_5\,{}^{\alpha D}k_{-5}k_{D/H}\,{}^{D2\alpha}k_5\,{}^{\alpha}k_{-5} + {}^{\alpha}k_{-5}k_{D/H}^{2}\,{}^{sec\beta}k_5\,{}^{\alpha D}k_{-5} + k_{H/D}k_{D/H}\,{}^{sec\beta}k_5\,{}^{\alpha D}k_{-5} + k_{D/H}\,{}^{sec\beta}k_5\,{}^{\alpha D}k_{-5} + {}^{sec\beta}k_{-5}k_{H/D}^{2}\,{}^{D2\alpha}k_5\,{}^{\alpha}k_{-5} + {}^{\alpha D}k_{-5}k_{H/D}^{2}\,{}^{D2\alpha}k_5\,{}^{\alpha}k_{-5} + {}^{sec\beta}k_5\,{}^{\alpha D}k_{-5}\,{}^{D2\alpha}k_5\,{}^{\alpha}k_{-5}k_{H/D} + {}^{\alpha}k_{-5}k_{H/D}k_{D/H}\,{}^{sec\beta}k_5\,{}^{\alpha D}k_{-5} + k_{H/D}\,{}^{\alpha D}k_5k_{D/H}\,{}^{sec\beta}k_5 + {}^{\alpha D}k_5k_{D/H}\,{}^{sec\beta}k_5 + k_{D/H}\,{}^{sec\beta}k_5 + k_{H/D}^{2}\,{}^{D2\alpha}k_5\,{}^{\alpha}k_{-5}\,{}^{\alpha D}k_5 + {}^{sec\beta}k_5\,{}^{D2\alpha}k_5\,{}^{\alpha}k_{-5}k_{H/D}\,{}^{\alpha D}k_5 + {}^{\alpha}k_{-5}k_{H/D}\,{}^{\alpha D}k_5k_{D/H}\,{}^{sec\beta}k_5 + k_{H/D}\,{}^{\alpha D}k_5\,{}^{sec\beta}k_{-5}k_{D/H} + {}^{\alpha D}k_5\,{}^{sec\beta}k_{-5}k_{D/H} + {}^{sec\beta}k_{-5}k_{D/H} + k_{D/H} + {}^{D2\alpha}k_5\,{}^{\alpha}k_{-5}k_{H/D}\,{}^{\alpha D}k_5\,{}^{sec\beta}k_{-5} + k_{D/H}\,{}^{\alpha}k_{-5}k_{H/D}\,{}^{\alpha D}k_5\,{}^{sec\beta}k_{-5}$$

Eq. S27

$$V^{d2-\alpha} = E_T\,{}^{D2\alpha}k_5\left(\frac{k_{H/D}^{2}\,{}^{\alpha D}k_5\,{}^{sec\beta}k_{-5} + {}^{\alpha D}k_5\,{}^{sec\beta}k_{-5}k_{H/D} + {}^{sec\beta}k_{-5}k_{H/D} + k_{H/D} + {}^{\alpha}k_{-5}k_{H/D}^{2}\,{}^{\alpha D}k_5\,{}^{sec\beta}k_{-5}}{\Delta}\right)$$

#### HDX leading to d$_2$-benzylsuccinate – beta pathway

**Assumption:**

Upon the formation of d$^2$-benzylscuccinate (BS) it is released to the solvent and due to its very low concentration with respect to the unsubstituted BS the reaction is irreversible. The BS concentration is so high that the enzyme saturated by the product and binding of the product does not influence the overall kinetics.

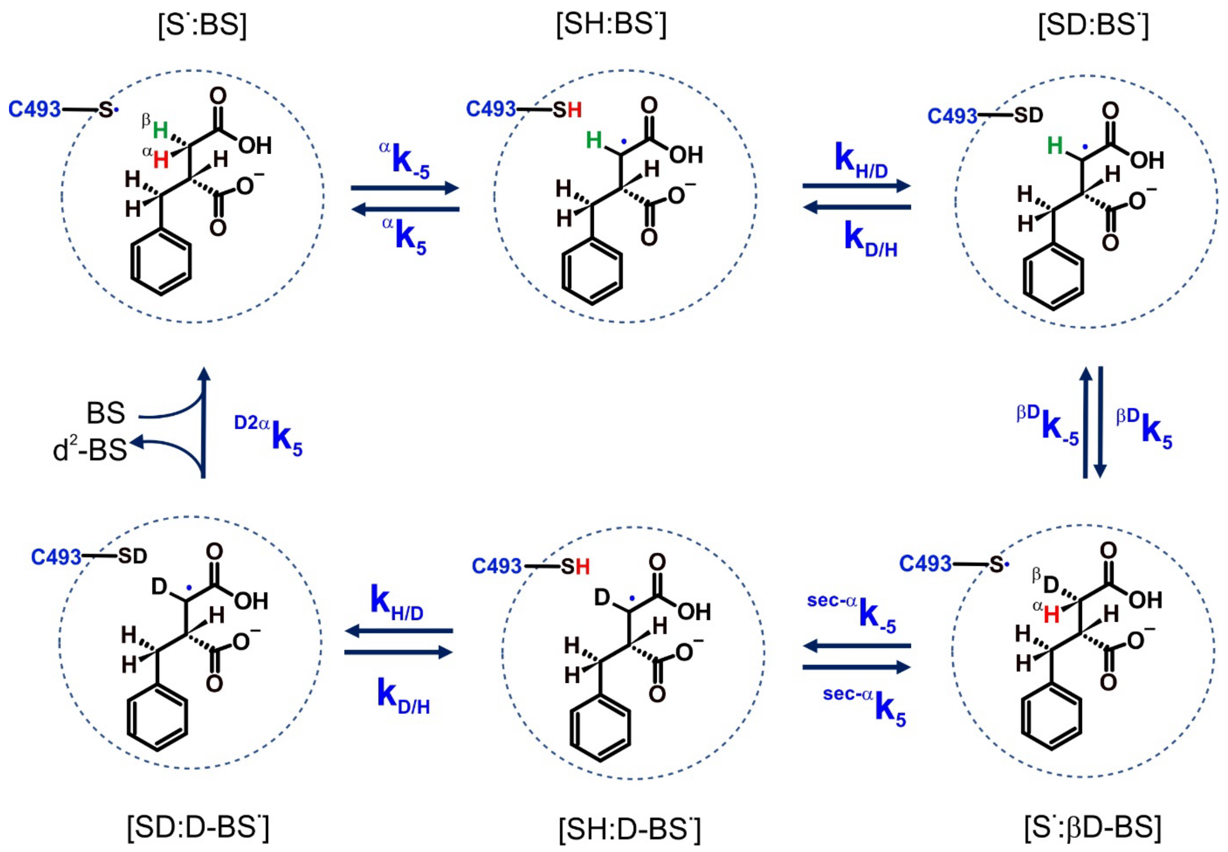

Scheme S4. The schematic representation of β pathway for H/D exchange in the BSS:BS complex during incubation in $D_2O$ leading to the production of $d_2$-benzylsuccinate assuming irreversibility of the D transfer to benzylsuccinyl radical and no kinetic limitation by product binding/release. [S·:BS] – BSS with radical Cys in complex with benzylsuccinate, [SH:BS·] – BSS with Cys-SH in complex with benzylsuccinyl radical, [SD:BS·] – BSS with Cys-SD in complex with benzylsuccinyl radical, [S·:βD-BS] BSS with radical Cys in complex with β-$d_1$-benzylsuccinate, [SH:D-BS·] - BSS with Cys-SH in complex with $d_1$-benzylsuccinyl radical, [SD:D-BS·] - BSS with Cys-SD in complex with $d_1$-benzylsuccinyl radical

Eq. S28

$$V^{d2-\beta} = {}^{D2\alpha}k_5[SD{:}D - BS^{\cdot}]$$

Eq. S29

$$\frac{[SD{:}D{-}BS^{\cdot}]}{E_T} = \frac{0 \cdot k_{\frac{H}{D}}^2\,{}^{\beta D}k_5\,{}^{sec\alpha}k_{-5} + {}^{\beta D}k_5\,{}^{sec\alpha}k_{-5}k_{\frac{H}{D}}\,{}^{\alpha}k_5 \cdot 0 + {}^{sec\alpha}k_{-5}k_{\frac{H}{D}}k_{\frac{D}{H}}\,{}^{\alpha}k_5 \cdot 0 + k_{\frac{H}{D}}\,{}^{\beta D}k_{-5}k_{\frac{D}{H}}\,{}^{\alpha}k_5 \cdot 0 + {}^{\alpha}k_{-5}k_{H/D}^2\,{}^{\beta D}k_5\,{}^{sec\alpha}k_{-5}}{\Delta} = \frac{{}^{\alpha}k_{-5}k_{H/D}^2\,{}^{\beta D}k_5\,{}^{sec\alpha}k_{-5}}{\Delta}$$

Eq. S30

$$\frac{[S^{\cdot}:BS]}{E_T} = \frac{k_{H/D}^2 {}^{\beta D}k_5 {}^{seca}k_{-5} {}^{D2\alpha}k_5 + {}^{\alpha}k_5 {}^{\beta D}k_5 {}^{seca}k_{-5} k_{H/D} {}^{D2\alpha}k_5 + k_{D/H} {}^{\alpha}k_5 {}^{seca}k_{-5} k_{H/D} {}^{D2\alpha}k_5}{\Delta} +$$

$$+ \frac{{}^{\beta D}k_{-5} k_{D/H} {}^{\alpha}k_5 k_{H/D} {}^{D2\alpha}k_5 + {}^{seca}k_5 {}^{\beta D}k_{-5} k_{D/H} {}^{\alpha}k_5 {}^{D2\alpha}k_5 + k_{D/H}^2 {}^{seca}k_5 {}^{\beta D}k_{-5} {}^{\alpha}k_5}{\Delta}$$

Eq. S31

$$\frac{[SH:BS^{\cdot}]}{E_T} = \frac{0 \cdot k_{D/H}^2 {}^{seca}k_5 {}^{\beta D}k_{-5} + {}^{\beta D}k_5 {}^{seca}k_{-5} k_{H/D} {}^{D2\alpha}k_5 {}^{\alpha}k_{-5} + k_{D/H} {}^{seca}k_{-5} k_{H/D} {}^{D2\alpha}k_5 {}^{\alpha}k_{-5}}{\Delta} +$$

$$\frac{{}^{\beta D}k_{-5} k_{D/H} k_{H/D} {}^{D2\alpha}k_5 {}^{\alpha}k_{-5} + {}^{seca}k_5 {}^{\beta D}k_{-5} k_{D/H} {}^{D2\alpha}k_5 {}^{\alpha}k_{-5} + {}^{\alpha}k_{-5} k_{D/H}^2 {}^{seca}k_5 {}^{\beta D}k_{-5}}{\Delta}$$

Eq. S32

$$\frac{[SD:BS^{\cdot}]}{E_T} = \frac{k_{H/D} \cdot 0 \cdot k_{D/H} {}^{seca}k_5 {}^{\beta D}k_{-5} + {}^{\alpha}k_5 \cdot 0 \cdot k_{D/H} {}^{seca}k_5 {}^{\beta D}k_{-5} + {}^{seca}k_{-5} k_{H/D}^2 {}^{D2\alpha}k_5 {}^{\alpha}k_{-5}}{\Delta} +$$

$$\frac{{}^{\beta D}k_{-5} k_{H/D}^2 {}^{D2\alpha}k_5 {}^{\alpha}k_{-5} + {}^{seca}k_5 {}^{\beta D}k_{-5} {}^{D2\alpha}k_5 {}^{\alpha}k_{-5} k_{H/D} + {}^{\alpha}k_{-5} k_{H/D} k_{D/H} {}^{seca}k_5 {}^{\beta D}k_{-5}}{\Delta}$$

Eq. S33

$$\frac{[S^{\cdot}:\beta DBS]}{E_T} = \frac{k_{H/D} {}^{\beta D}k_5 \cdot 0 \cdot k_{D/H} {}^{seca}k_5 + {}^{\alpha}k_5 \cdot 0 \cdot {}^{\beta D}k_5 k_{D/H} {}^{seca}k_5 + k_{H/D} {}^{\alpha}k_5 \cdot 0 \cdot k_{D/H} {}^{seca}k_5}{\Delta} +$$

$$\frac{k_{H/D}^2 {}^{D2\alpha}k_5 {}^{\alpha}k_{-5} {}^{\beta D}k_5 + {}^{seca}k_5 {}^{D2\alpha}k_5 {}^{\alpha}k_{-5} k_{H/D} {}^{\beta D}k_5 + {}^{\alpha}k_{-5} k_{H/D} {}^{\beta D}k_5 k_{D/H} {}^{seca}k_5}{\Delta}$$

Eq. S34

$$\frac{[SH:DBS^{\cdot}]}{E_T} = \frac{0 \cdot k_{H/D} {}^{\beta D}k_5 {}^{seca}k_{-5} k_{D/H} + {}^{\alpha}k_5 \cdot 0^{\beta D}k_5 {}^{seca}k_{-5} k_{D/H} + k_{H/D} {}^{\alpha}k_5 \cdot 0 \cdot {}^{seca}k_{-5} k_{D/H}}{\Delta} +$$

$$\frac{{}^{\beta D}k_{-5} k_{H/D} {}^{\alpha}k_5 \cdot 0 \cdot k_{D/H} + {}^{D2\alpha}k_5 {}^{\alpha}k_{-5} k_{H/D} {}^{\beta D}k_5 {}^{seca}k_{-5} + k_{D/H} {}^{\alpha}k_{-5} k_{H/D} {}^{\beta D}k_5 {}^{seca}k_{-5}}{\Delta}$$

$$= \frac{{}^{D2\alpha}k_5 {}^{\alpha}k_{-5} k_{H/D} {}^{\beta D}k_5 {}^{seca}k_{-5} + k_{D/H} {}^{\alpha}k_{-5} k_{H/D} {}^{\beta D}k_5 {}^{seca}k_{-5}}{\Delta}$$

Eq. S35

$$V^{d2-\beta} = \frac{{}^{\alpha}k_{-5} k_{H/D}^2 {}^{\beta D}k_5 {}^{seca}k_{-5} {}^{D2\alpha}k_5}{\Delta} E_T$$

Eq. S36

$\Delta = {}^{\alpha}k_{-5} k_{\frac{H}{D}}^2 {}^{\beta D}k_5 {}^{seca}k_{-5} +$

$k_{\frac{H}{D}}^2 {}^{\beta D}k_5 {}^{seca}k_{-5} {}^{D2\alpha}k_5 + {}^{\alpha}k_5 {}^{\beta D}k_5 {}^{seca}k_{-5} k_{\frac{H}{D}} {}^{D2\alpha}k_5 + k_{\frac{D}{H}} {}^{\alpha}k_5 {}^{seca}k_{-5} k_{\frac{H}{D}} {}^{D2\alpha}k_5 +$

$+ {}^{\beta D}k_{-5} k_{\frac{D}{H}} {}^{\alpha}k_5 k_{\frac{H}{D}} {}^{D2\alpha}k_5 + {}^{seca}k_5 {}^{\beta D}k_{-5} k_{\frac{D}{H}} {}^{\alpha}k_5 {}^{D2\alpha}k_5 + k_{\frac{D}{H}}^2 {}^{seca}k_5 {}^{\beta D}k_{-5} {}^{\alpha}k_5$

$$+{}^{\beta D}k_5{}^{sec\alpha}k_{-5}k_{\frac{H}{D}}{}^{D2\alpha}k_5{}^{\alpha}k_{-5} + k_{\frac{D}{H}}{}^{sec\alpha}k_{-5}k_{\frac{H}{D}}{}^{D2\alpha}k_5{}^{\alpha}k_{-5} + {}^{\beta D}k_5{}^{sec\alpha}k_{-5}k_{\frac{H}{D}}{}^{D2\alpha}k_5{}^{\alpha}k_{-5}$$

$$+k_{\frac{D}{H}}{}^{sec\alpha}k_{-5}k_{\frac{H}{D}}{}^{D2\alpha}k_5{}^{\alpha}k_{-5} +$$

$$+{}^{sec\alpha}k_{-5}k_{H/D}^2{}^{D2\alpha}k_5{}^{\alpha}k_{-5} + {}^{\beta D}k_{-5}k_{H/D}^2{}^{D2\alpha}k_5{}^{\alpha}k_{-5} + {}^{sec\alpha}k_5{}^{\beta D}k_{-5}{}^{D2\alpha}k_5{}^{\alpha}k_{-5}k_{H/D}$$

$$+{}^{\alpha}k_{-5}k_{H/D}k_{D/H}{}^{sec\alpha}k_5{}^{\beta D}k_{-5} +$$

$$+k_{H/D}^2{}^{D2\alpha}k_5{}^{\alpha}k_{-5}{}^{\beta D}k_5 + {}^{sec\alpha}k_5{}^{D2\alpha}k_5{}^{\alpha}k_{-5}k_{H/D}{}^{\beta D}k_5 + {}^{\alpha}k_{-5}k_{H/D}{}^{\beta D}k_5k_{D/H}{}^{sec\alpha}k_5$$

$$+k_{H/D}^2{}^{D2\alpha}k_5{}^{\alpha}k_{-5}{}^{\beta D}k_5 + {}^{sec\alpha}k_5{}^{D2\alpha}k_5{}^{\alpha}k_{-5}k_{H/D}{}^{\beta D}k_5 + {}^{\alpha}k_{-5}k_{H/D}{}^{\beta D}k_5k_{D/H}{}^{sec\alpha}k_5$$

$$+{}^{D2\alpha}k_5{}^{\alpha}k_{-5}k_{H/D}{}^{\beta D}k_5{}^{sec\alpha}k_{-5} + k_{D/H}{}^{\alpha}k_{-5}k_{H/D}{}^{\beta D}k_5{}^{sec\alpha}k_{-5}$$

### RESULTS

#### RMSD

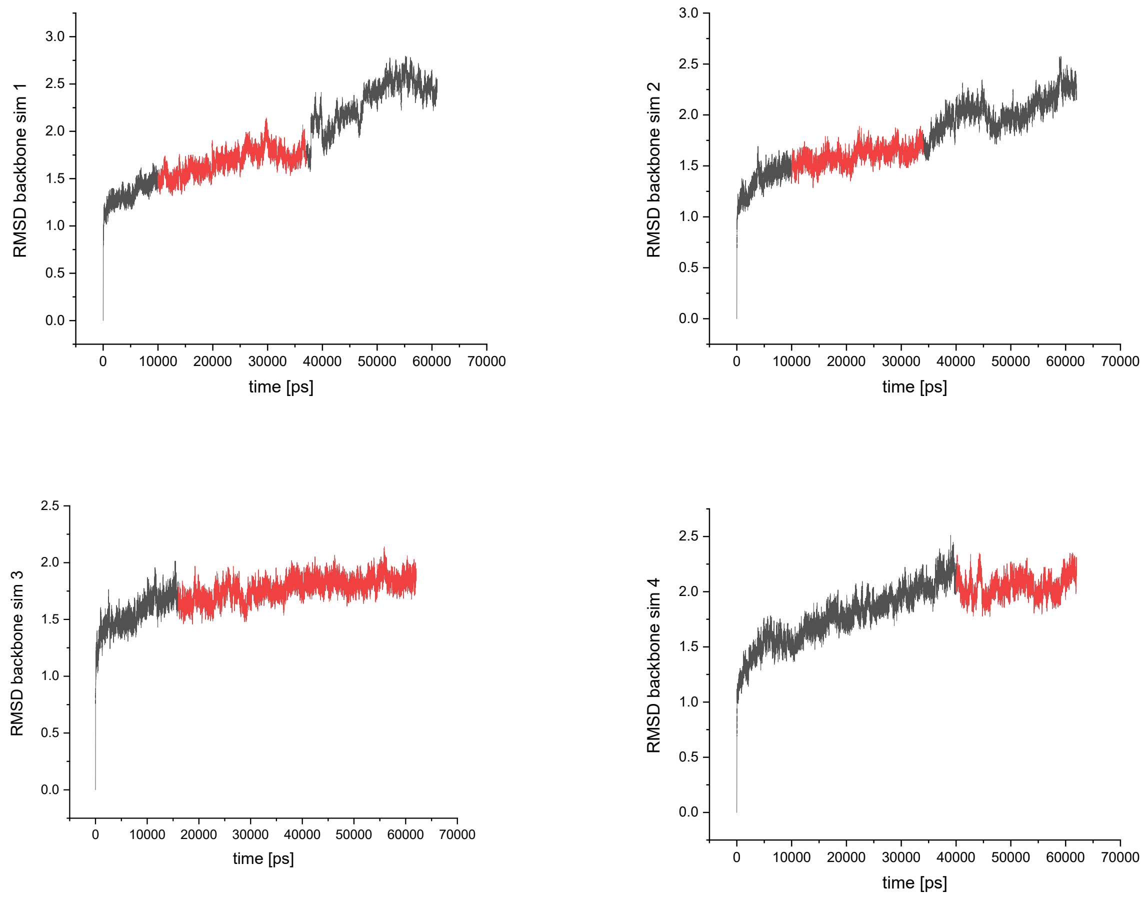

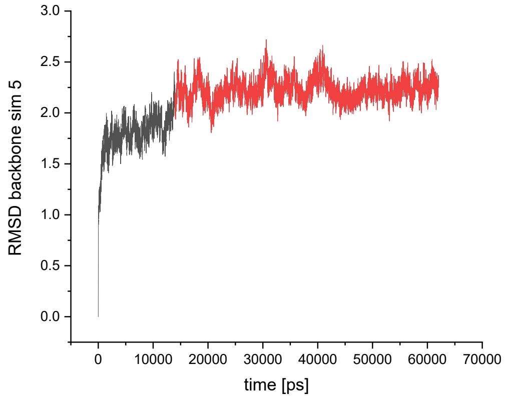

Figure S2. RMSD of the backbone for five MD simulations with radical Gly, toluene and mono-protonated fumarate in pro*R* orientation. The red parts were considered in statistical analysis.

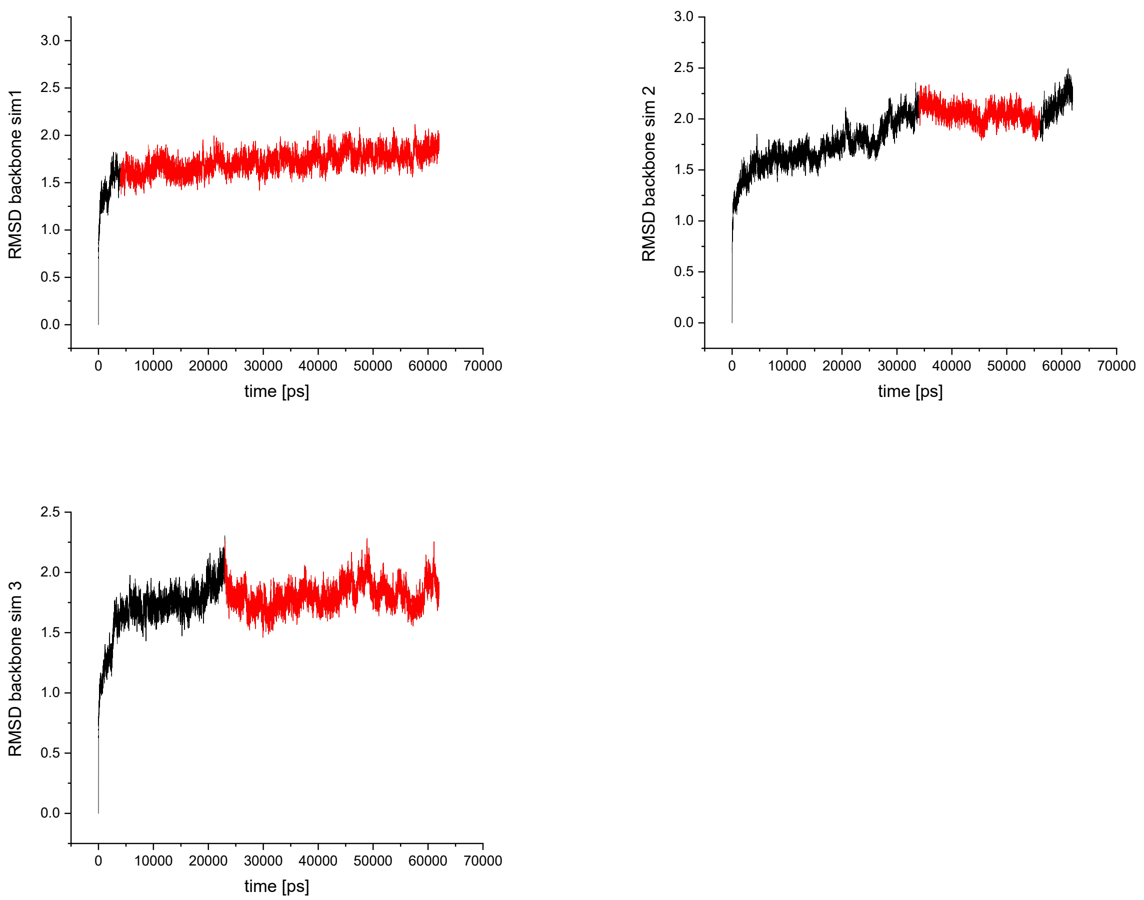

Figure S3. RMSD of the backbone for three MD simulations with radical Gly, toluene and mono-protonated fumarate in pro*S* orientation. The red parts were considered in statistical analysis.

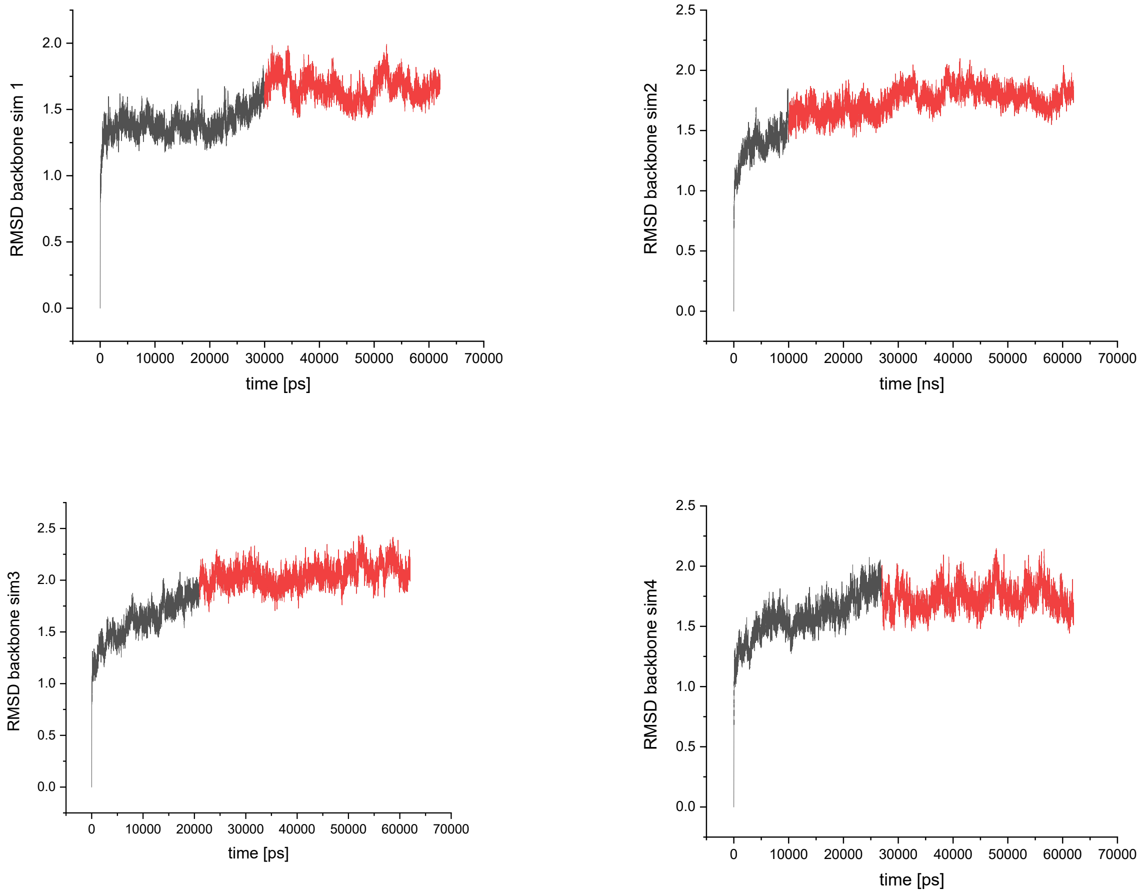

Figure S4. RMSD of the backbone for four MD simulations with radical Cys, toluene and mono-protonated fumarate. The red parts were considered in statistical analysis.

A B

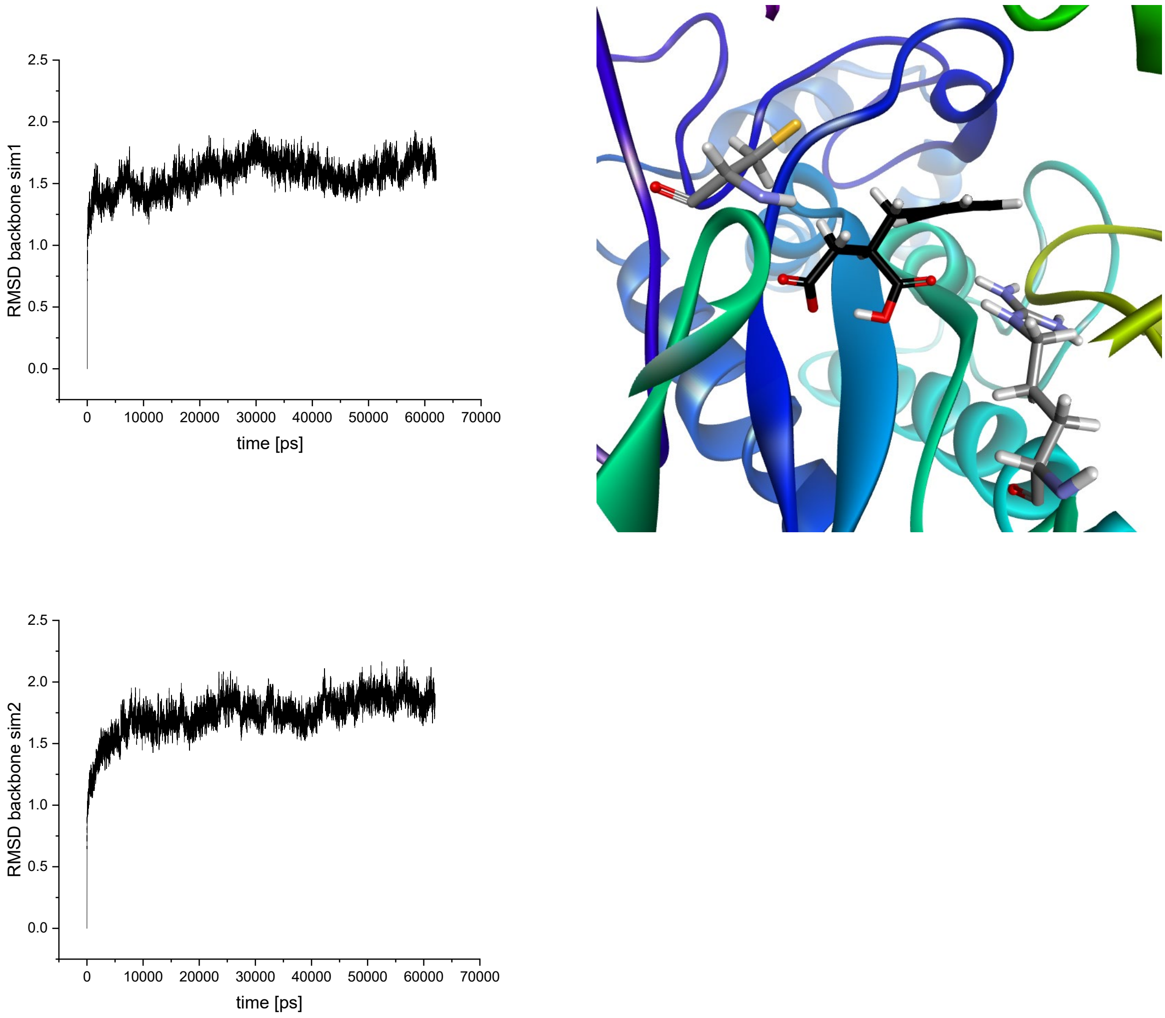

Figure S5. A) RMSD of the backbone for two MD simulations of BSS with radical Cys493 and mono-protonated benzylsuccinate (protonation at carboxyl group close to Arg508). B) figure presenting protonation mode

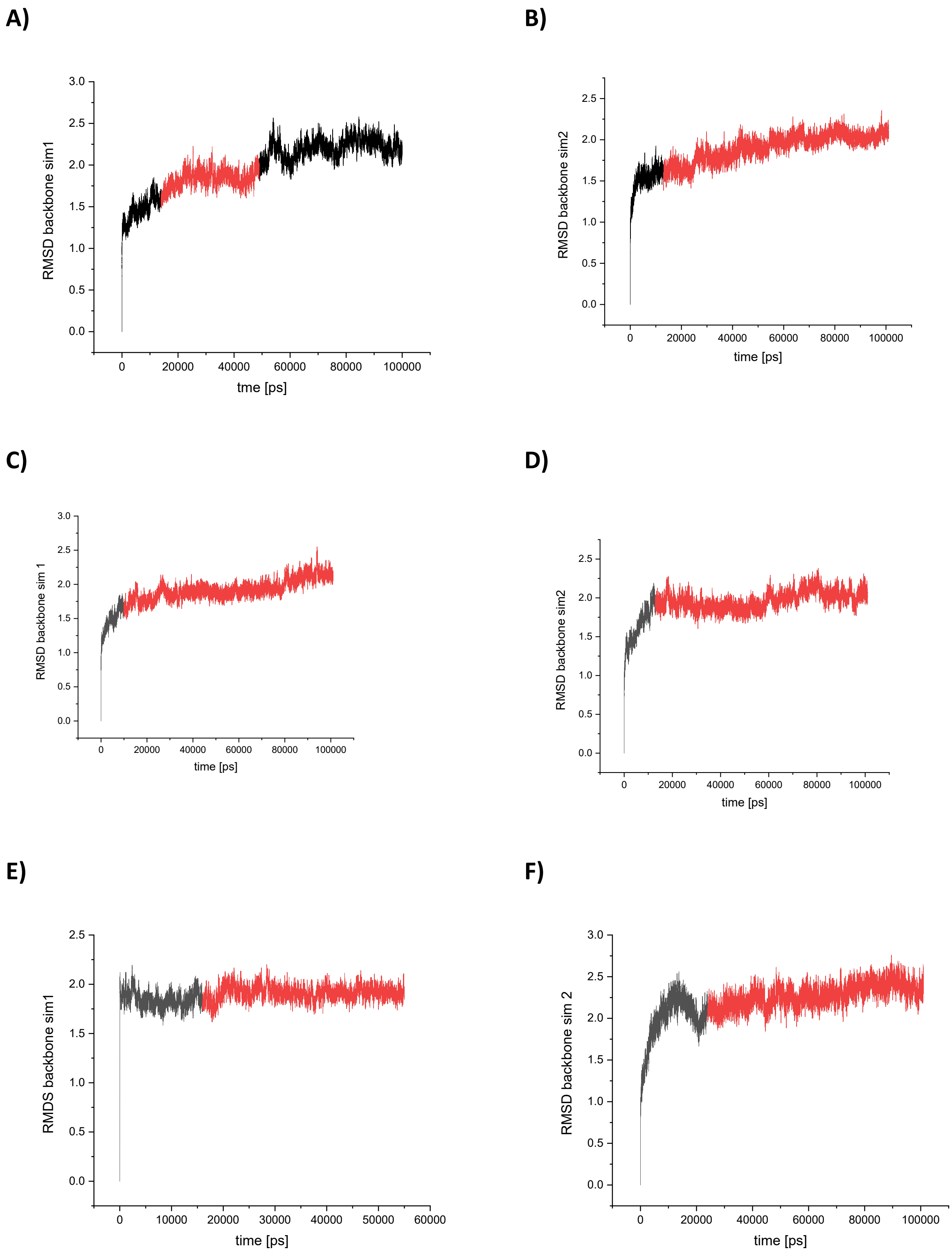

Figure S6. RMSD of the backbone for A and B) simulations with mono-protonated fumarate in pro*R* orientation, C and D) simulations with mono-protonated fumarate in pro*R* orientation with a rotated proximal carboxyl group, E and F) simulations with mono-protonated fumarate in pro*S* orientation. The red parts were considered in the MMPBSA statistical analysis.

#### Step 1 – activation of Cys493

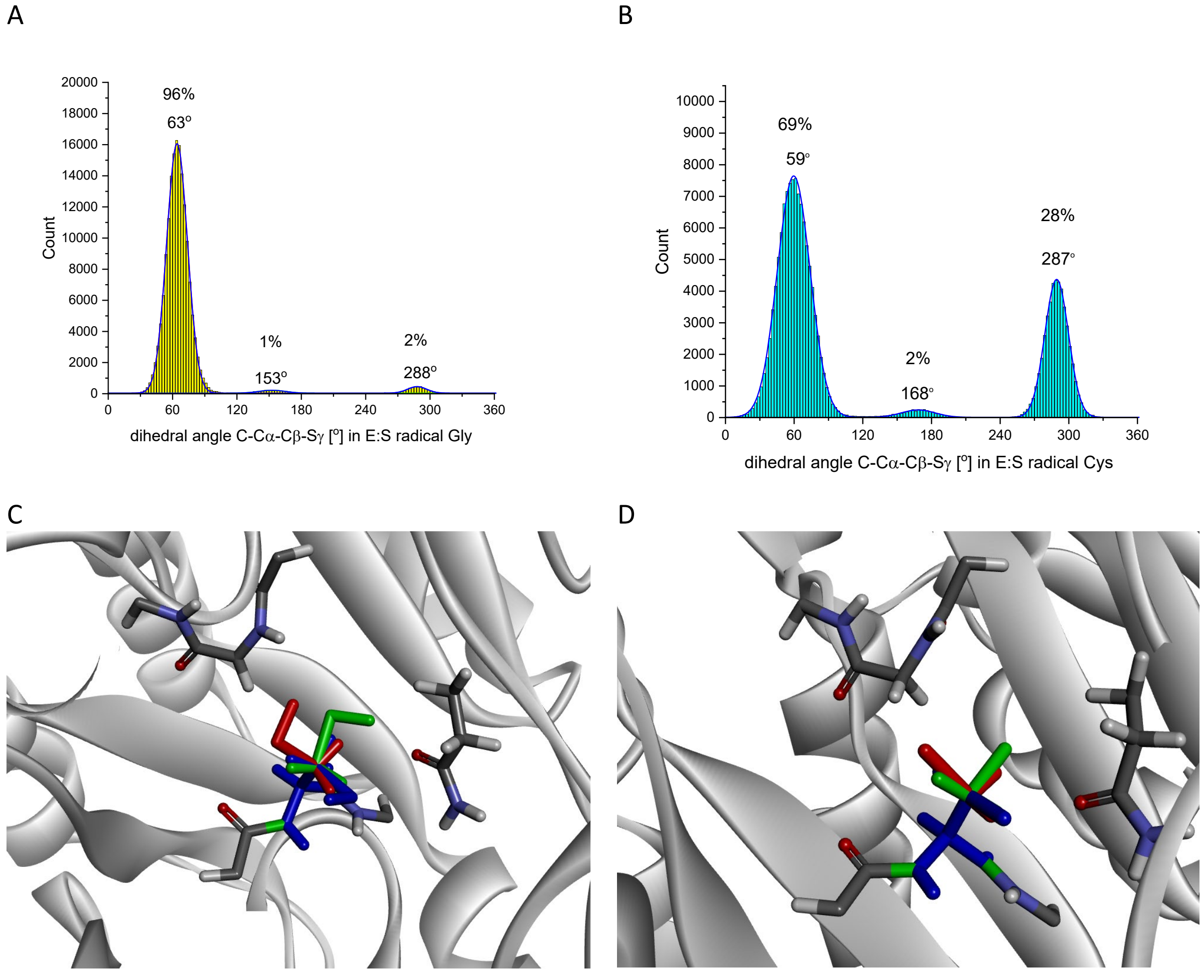

Figure S7. Distribution of C-Cα-Cβ-Sγ dihedral angle in Cys493 for A) 5 MD simulation of E:S with radical Gly829, B) 4 MD simulation of E:S with radical Cys493; Geometry representing C) 3 conformation populations of Cys493: 63° green, 153° red, 288° blue; D) 3 conformation populations of radical Cys493: 59° green, 168° red, 287° blue

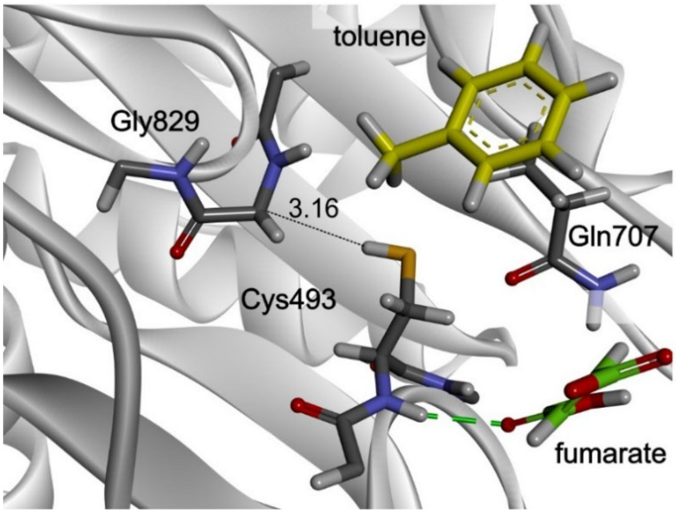

Figure S8. The structure of the E:S with rotated Cys493 SH group.

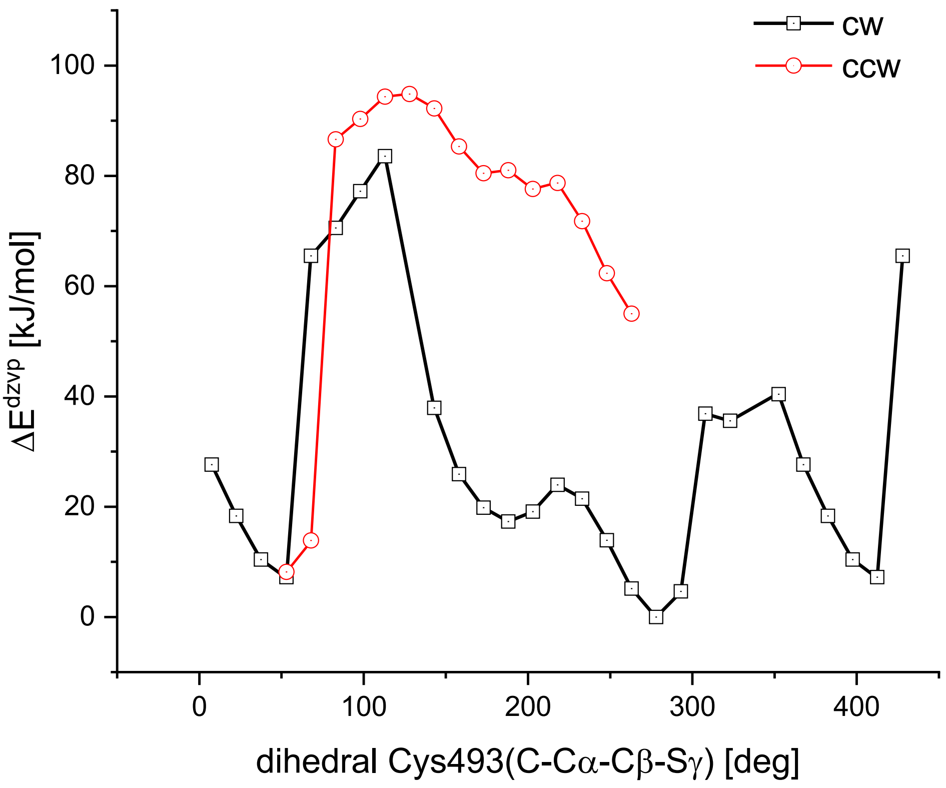

Figure S9.The rotational analysis for radical Cys493. The energy was collected at B3LYP:AMBER/dzvp level of theory with loose convergence criteria.

#### Step 2 – activation of toluene

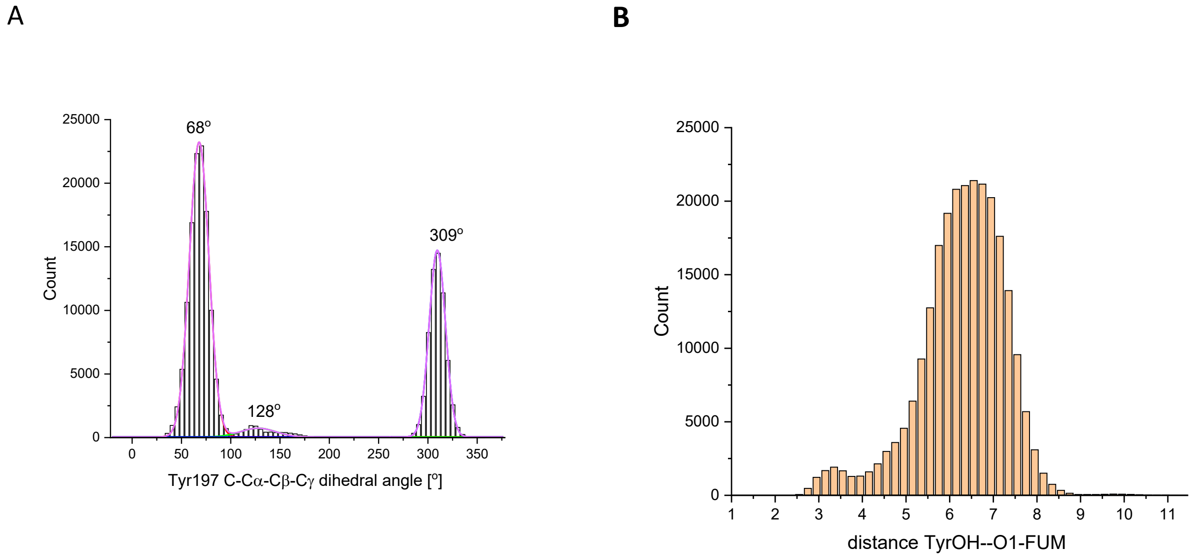

Figure S10. The analysis of Tyr197 during MD simulations: A) distribution of C-Cα-Cβ-Cγ dihedral angle of Tyr, B) distribution of the distance between Tyr OH group and O1 atom of fumarate.

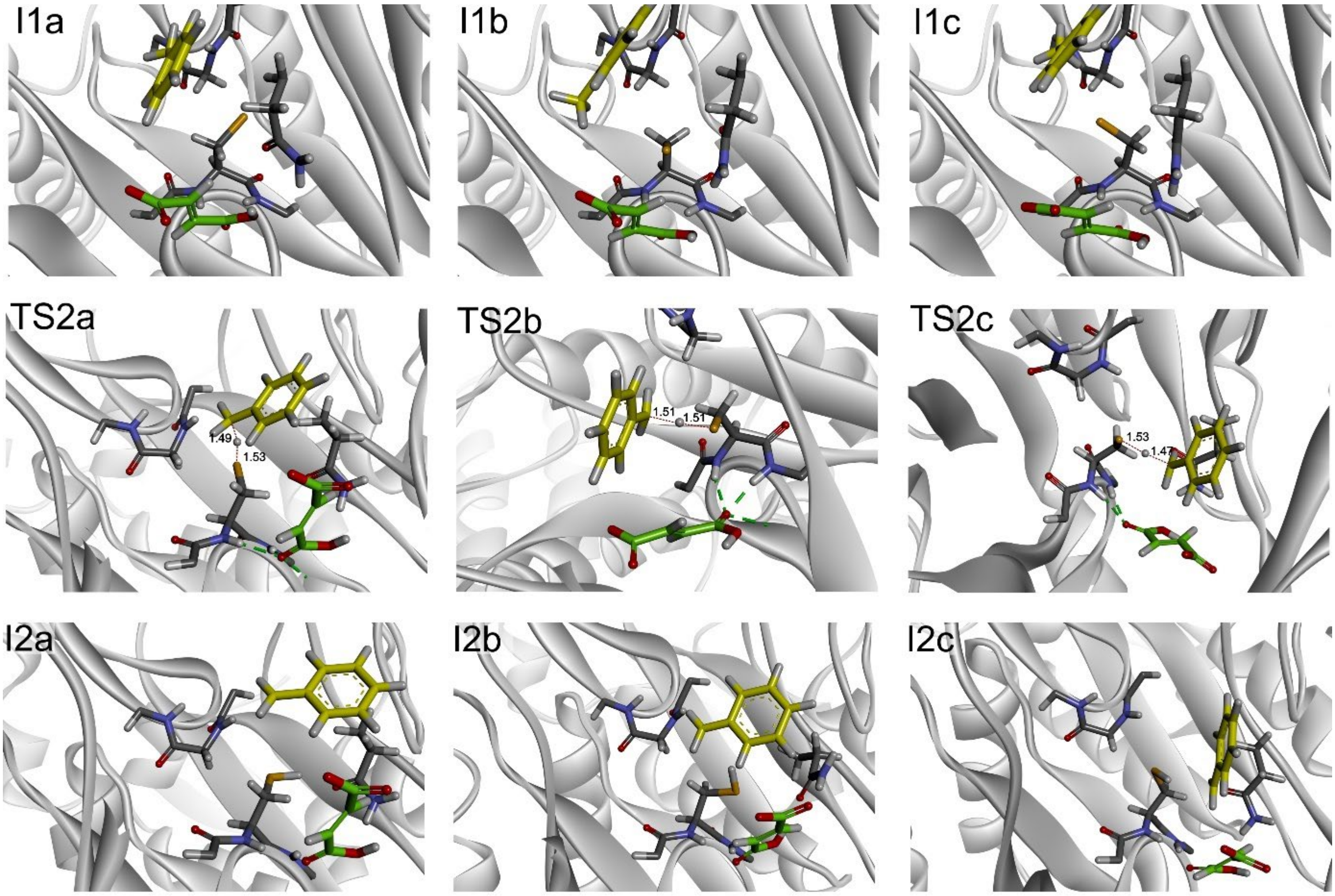

Figure S11. The geometries of toluene activation in all three variants depending on the conformation of Cys493.

#### Step 3 - C-C bond formation

##### Proximal C2

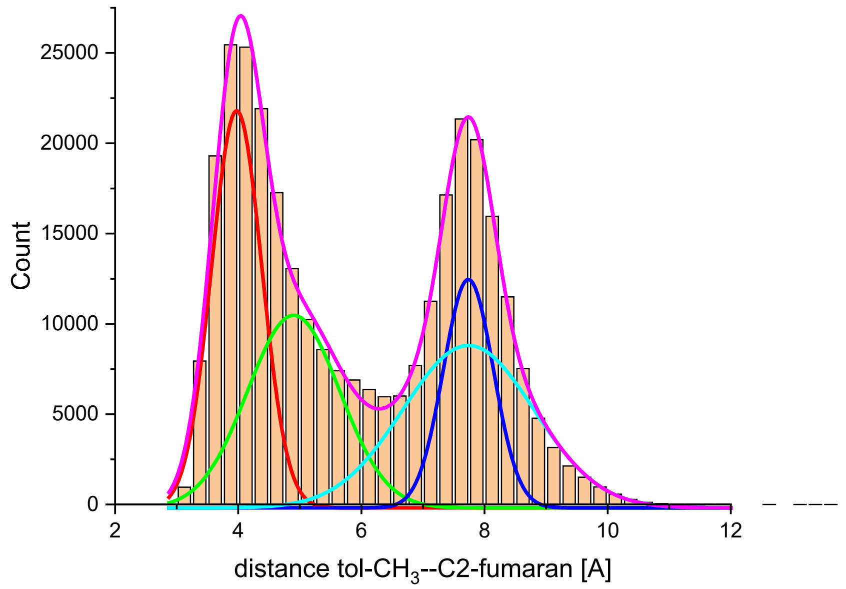

| N frames | Mean [Å] | Median [Å] | Min [Å] | Max [Å] |
|---|---|---|---|---|
| 308929 | 5.95 | 5.65 | 2.94 | 13.59 |

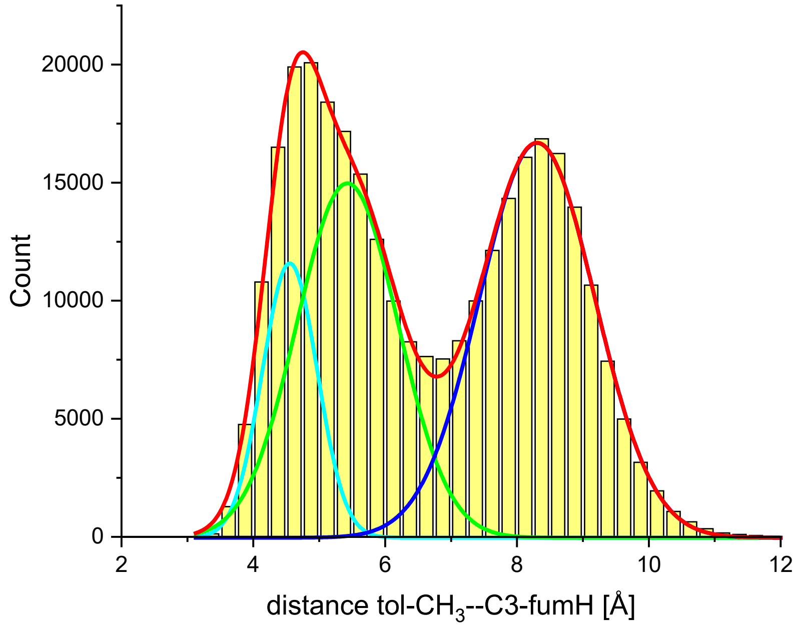

| N frames | Mean [Å] | Median [Å] | Min [Å] | Max [Å] |
|---|---|---|---|---|
| 308929 | 6.67 | 6.48 | 3.17 | 14.54 |

Figure S12. Distances between distal (C2) and proximal( C3) atoms of fumarate double bond and benzyl C atom of toluene during 5 independent 62 ns MD simulations of pro*R* bound monoprotonated fumarate.

Step 4 – quenching the radical benzylsuccinate intermediate

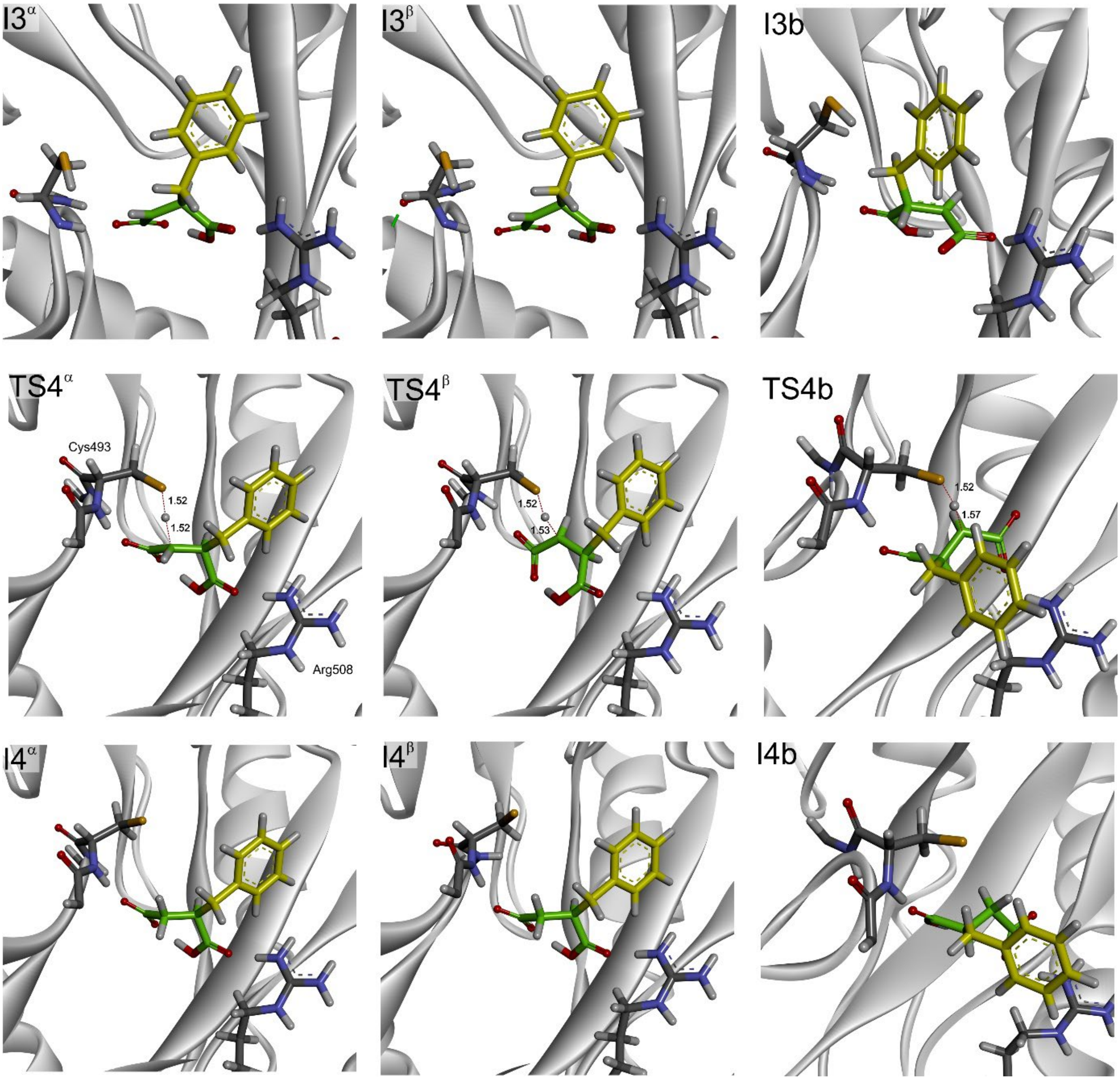

Figure S13. The geometries of benzylsuccinate radical quenching by HAT from Cys493.

Step 5 – transfer of the H atom between Gly and Cys for the E:P complex.

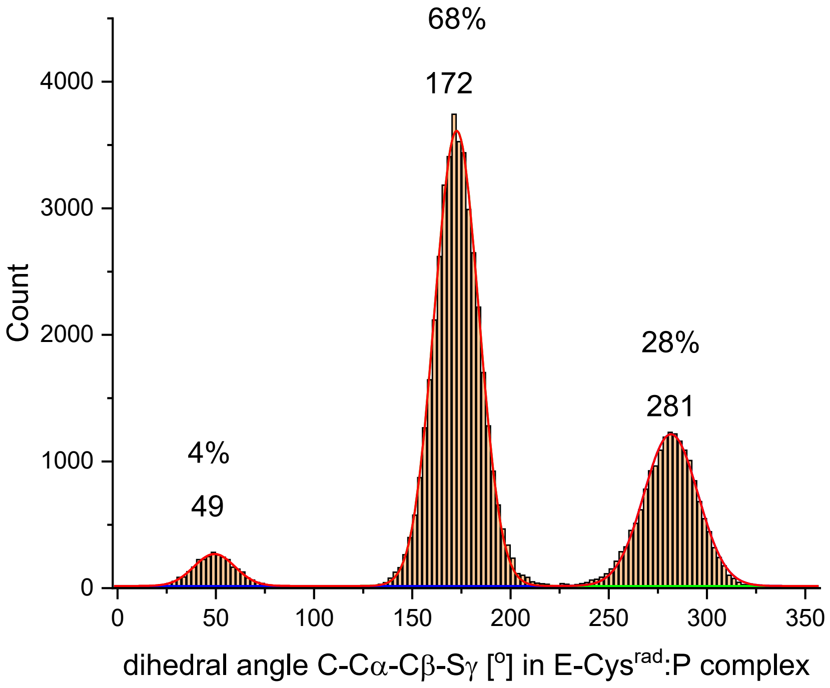

Figure S14. Distribution of dihedral angle C-Cα-Cβ-Cγ in radical Cys493 in two independent MD simulations conducted for E:P (fumarate mono-protonated on carboxyl group closer to Arg508).

#### Reaction enantioselectivity

A

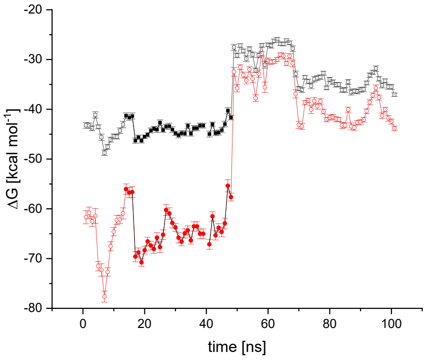

B

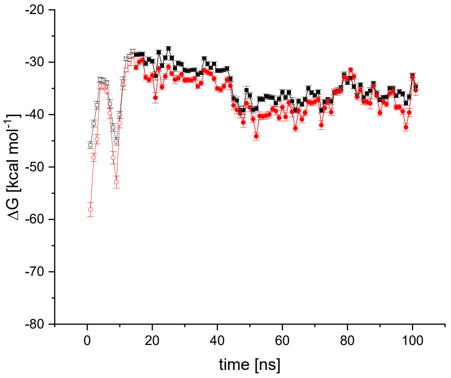

C

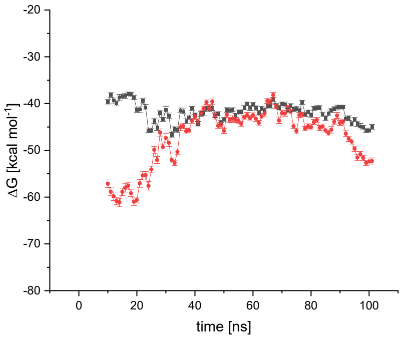

D

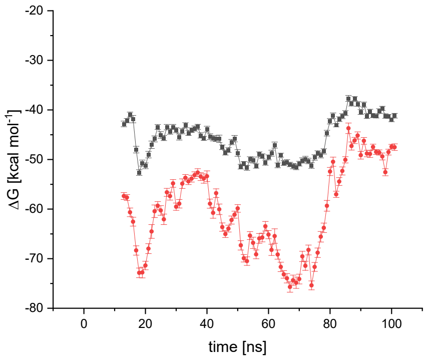

E

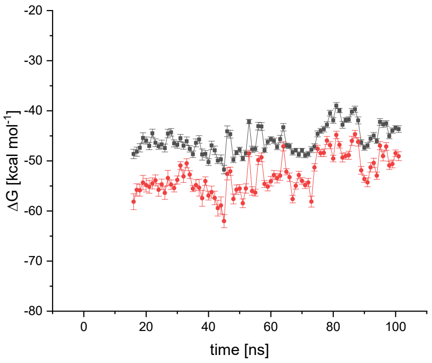

F

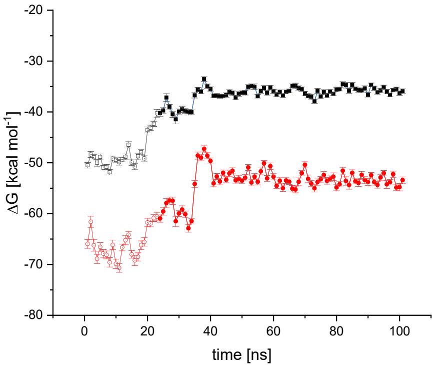

Figure S15. The results of MM/PBSA ΔG calculations from MD simulations of BSS:mono-protonated fumarate: A-B) pro*R* bound fumarate, C-D) pro*R* fumarate with a rotated protonated carboxyl group, E-F) pro*S* bound fumarate. Black line – Generalized Born, Red line – Poisson-Boltzmann, full symbol – points selected for the analysis

**Distal C2**

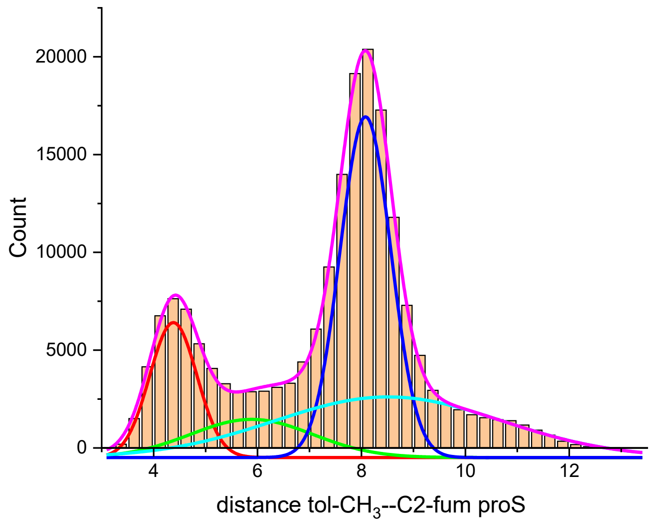

| N total | Mean [Å] | Median [Å] | Min [Å] | Max [Å] |
|---|---|---|---|---|
| 186000 | 7.33 | 7.81 | 3.13 | 13.22 |

**Proximal C3**

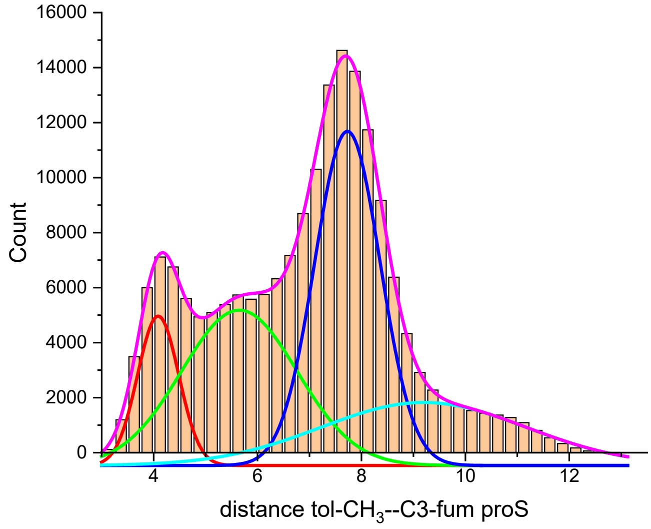

| N total | Mean [Å] | Median [Å] | Min [Å] | Max [Å] |
|---|---|---|---|---|
| 186000 | 6.91 | 7.20 | 3.02 | 13.13 |

Figure S16. Distances between distal (C2) and proximal( C3) atoms of fumarate double bond and benzyl C atom of toluene during 3 independent MD simulations of proS bound monoprotonated fumarate.

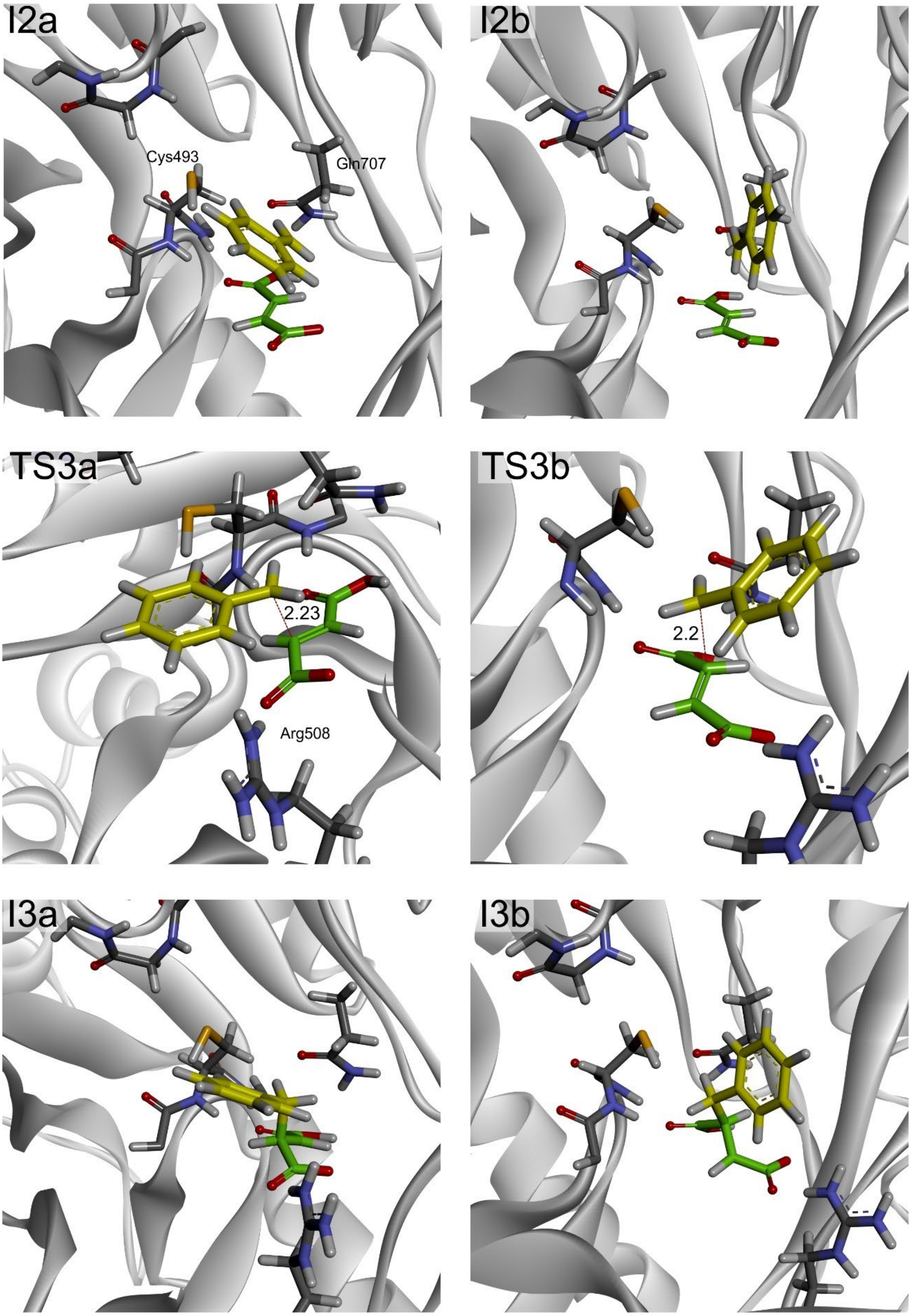

Figure S17. C-C bond formation for the pro*S* bound fumarate: a) C-C bond formed between benzyl radical and distal C3 atom of fumarate, b) C-C bond formed between benzyl radical and proximal C2 atom of fumarate

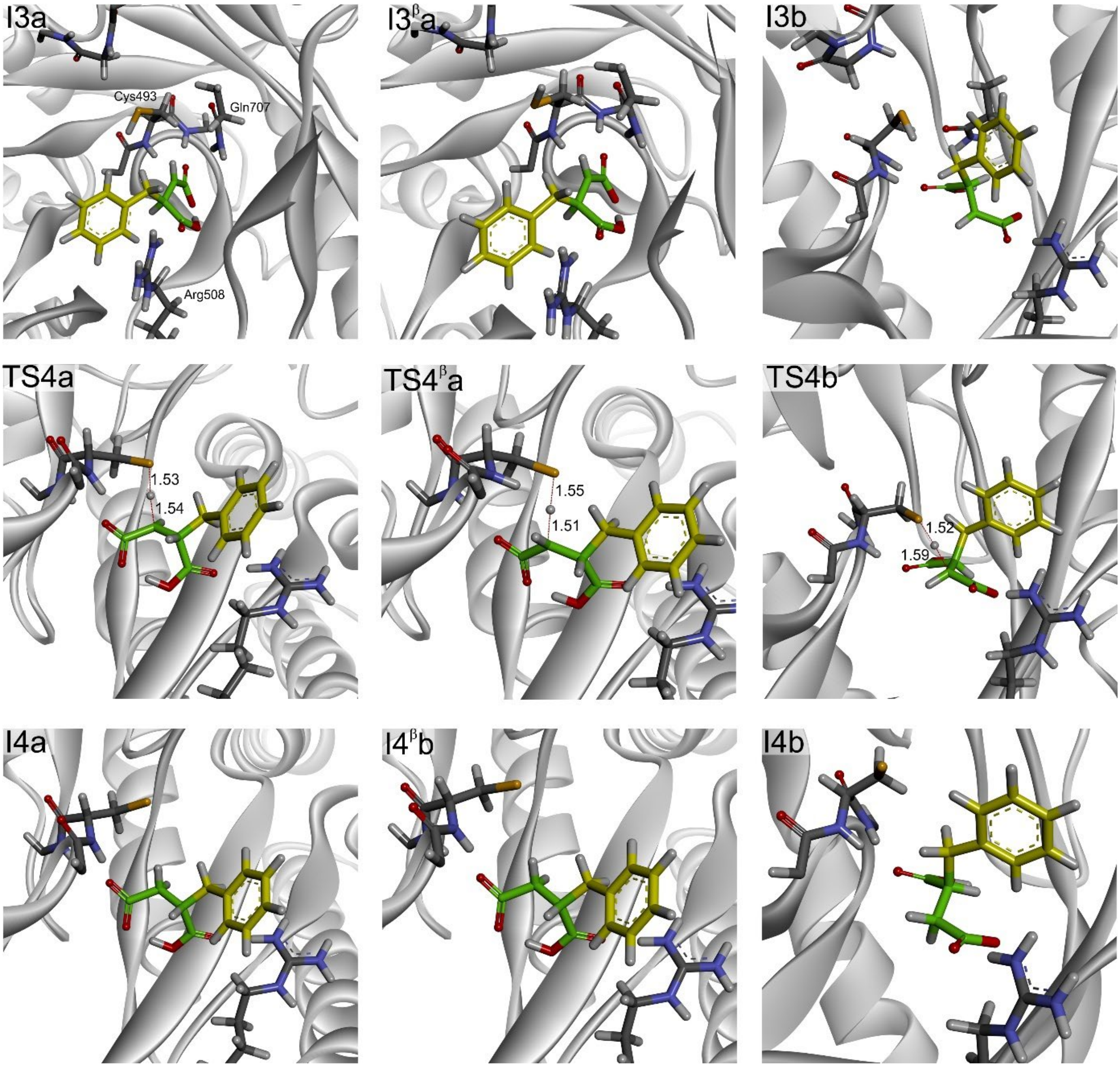

Figure S18. *S*-benzyl succinate radical quenching by Cys493: a) syn HAT from Cys to proximal C2 atom, a$^{\beta}$) anti HAT from Cys to proximal C2 atom, b) HAT from Cys to distal C3 atom.

#### QM:MM energies

Table S4. Energies and vibration corrections for the pro*R* pathway.

| [Ha]: | $E^{QM1}$ dzvp | $E^{QM1}$ tzvp | $E^{QM2}$ dzvp | $E^{QM2}$ tzvp | ZPE | Thermal | H | G |
|---|---|---|---|---|---|---|---|---|
| **Step 1 – activation of Cys493** | | | | | | | | |
| **ES** | -2365.55824493968 | -2366.20961790382 | -8363.75303975607 | -8366.11453637253 | 24.536596 | 26.05049 | 26.051434 | 23.045644 |
| **ES Hrot** | -2365.55541742932 | -2366.20838338787 | -8363.74757941795 | -8366.11196473799 | 24.535406 | 26.047769 | 26.048713 | 23.046833 |
| **TS1** | -2365.53795032585 | -2366.19090709356 | -8363.73073384547 | -8366.09067216491 | 24.535406 | 26.047769 | 26.048713 | 23.046833 |
| **I1** | -2365.58193794600 | -2366.23301809200 | -8363.76789923912 | -8366.13003621295 | 24.541351 | 26.054725 | 26.055669 | 23.050805 |
| **Step 2 – activation of toluene** | | | | | | | | |
| **I1a** | -2365.56870369175 | -2366.22015505998 | -8363.76724805284 | -8366.12678460707 | 24.545021 | 26.057341 | 26.058285 | 23.055568 |
| **TS2a** | -2365.53155577164 | -2366.18051221079 | -8363.72820225284 | -8366.09061137933 | 24.538529 | 26.049914 | 26.050859 | 23.050998 |
| **I2a** | -2365.55754977988 | -2366.20869661411 | -8363.75497339268 | -8366.11539186589 | 24.54145 | 26.053615 | 26.054559 | 23.053912 |
| **I1b** | -2365.56633542016 | -2366.21922226430 | -8363.75839686177 | -8366.11502261967 | 24.545859 | 26.05733 | 26.058274 | 23.059524 |
| **TS2b** | -2365.52889781096 | -2366.17954212640 | -8363.71632450491 | -8366.07923348779 | 24.538435 | 26.04933 | 26.050274 | 23.05372 |
| **I2b** | -2365.54476757033 | -2366.20869661411 | -8363.74006439215 | -8366.10177296496 | 24.54129 | 26.053183 | 26.054127 | 23.054493 |
| **I1c** | -2365.56646282554 | -2366.21885058225 | -8363.76453764535 | -8366.12173925949 | 24.545703 | 26.056605 | 26.057549 | 23.060942 |
| **TS2** | -2365.54112015292 | -2366.19014262673 | -8363.73779947531 | -8366.09838162977 | 24.540468 | 26.051058 | 26.052002 | 23.056516 |
| **I2c** | -2365.55023224354 | -2366.20132139113 | -8363.74813172313 | -8366.10911066041 | 24.543131 | 26.054684 | 26.055628 | 23.056729 |
| **Step 3 - C-C bond formation** | | | | | | | | |
| **I2c** | -2365.45194508504 | -2366.10068922705 | -8363.65402355762 | -8366.03305887327 | 24.547839 | 26.056803 | 26.057747 | 23.069128 |
| **TS3** | -2365.42852539412 | -2366.07341173620 | -8363.63936883283 | -8366.01487514838 | 24.549072 | 26.056749 | 26.057693 | 23.072086 |
| **I3** | -2365.48143694255 | -2366.11856395130 | -8363.67678717914 | -8366.04930408459 | 24.553187 | 26.059255 | 26.060199 | 23.080335 |
| **I2b** | -2365.45084761027 | -2366.10239087908 | -8363.65708136116 | -8366.03495042814 | 24.547638 | 26.057178 | 26.058122 | 23.067142 |
| **TS3b** | -2365.41300912051 | -2366.05928368785 | -8363.61849203256 | -8366.00035660046 | 24.550074 | 26.057441 | 26.058385 | 23.072397 |
| **I3b** | -2365.45081421471 | -2366.09407121021 | -8363.65066529819 | -8366.02959948585 | 24.552279 | 26.0595 | 26.060445 | 23.075012 |
| **Step 4 – quenching the radical benzylsuccinate intermediate** | | | | | | | | |
| **$I3^{\alpha}a$** | -2365.50700866888 | -2366.14346375052 | -8363.70579042290 | -8366.07779821857 | 24.552585 | 26.059258 | 26.060202 | 23.07772 |

| | | | | | | | | |
|---|---|---|---|---|---|---|---|---|
| **TS4 $^{\alpha}$a** | -2365.48588258744 | -2366.11938122718 | -8363.68242424925 | -8366.05242786375 | 24.550248 | 26.055587 | 26.056531 | 23.07844 |
| **I4$^{\alpha}$a** | -2365.52336320995 | -2366.15670556527 | -8363.71865395157 | -8366.08816044918 | 24.55777 | 26.063373 | 26.064317 | 23.084599 |
| **I3$^{\beta}$a** | -2365.52576542669 | -2366.16263312239 | -8363.73225598623 | -8366.10008982158 | 24.555723 | 26.061032 | 26.061976 | 23.082117 |
| **TS4 $^{\beta}$a** | -2365.47538833495 | -2366.11195044818 | -8363.68210748019 | -8366.05308283158 | 24.549209 | 26.055177 | 26.056122 | 23.074296 |
| **I4$^{\beta}$a** | -2365.53603896205 | -2366.16686854738 | -8363.74803588762 | -8366.11246640593 | 24.559458 | 26.06443 | 26.065374 | 23.0867 |
| **I3b** | -2365.45081421471 | -2366.09407121021 | -8363.65066529819 | -8366.02959948585 | 24.552279 | 26.0595 | 26.060445 | 23.075012 |
| **TS4b** | -2365.41615080984 | -2366.05948626481 | -8363.63186527267 | -8366.00595887116 | 24.549184 | 26.055144 | 26.056088 | 23.076993 |
| **I4b** | -2365.46395102289 | -2366.10700481023 | -8363.67341351608 | -8366.04754834518 | 24.554089 | 26.061216 | 26.06216 | 23.079125 |
| **Step 5 – transfer of the H atom between Gly and Cys for the E:P complex.** | | | | | | | | |
| **I4_rot** | -2365.52559254147 | -2366.15729078714 | -8363.71785348512 | -8366.08581950566 | 24.557753 | 26.063432 | 26.064376 | 23.084624 |
| **I4** | -2365.52384603110 | -2366.15562248325 | -8363.71442505103 | -8366.08357385082 | 24.558912 | 26.064159 | 26.065103 | 23.087309 |
| **TS5** | -2365.47814153330 | -2366.11109418045 | -8363.67055202975 | -8366.03996412632 | 24.552674 | 26.057067 | 26.058011 | 23.08285 |
| **EP** | -2365.49737337207 | -2366.13196334355 | -8363.69288270337 | -8366.06449495651 | 24.555323 | 26.060797 | 26.061741 | 23.083324 |

Table S5. Energies and vibration corrections for the pro*S* pathway.

| | $E^{QM1}$ dzvp | $E^{QM1}$ tzvp | $E^{QM2}$ dzvp | $E^{QM2}$ tzvp | ZPE | Thermal | H | G |
|---|---|---|---|---|---|---|---|---|
| **Step 3 - C-C bond formation** | | | | | | | | |
| **I2a** | -2365.45426513554 | -2366.10515765690 | -8363.66279734439 | -8366.04041271922 | 24.550016 | 26.058578 | 26.059522 | 23.070180 |
| **TS3a** | -2365.44062609987 | -2366.08968192676 | -8363.65491684864 | -8366.03077507779 | 24.552389 | 26.058983 | 26.059927 | 23.075266 |
| **I3a** | -2365.47062824028 | -2366.11851152391 | -8363.68324805964 | -8366.05684786764 | 24.553318 | 26.060496 | 26.061440 | 23.076421 |
| **I2b** | -2365.46114306825 | -2366.11077446835 | -8363.66653417132 | -8366.04206937598 | 24.548774 | 26.057541 | 26.058485 | 23.068212 |
| **TS3b** | -2365.42980640614 | -2366.07785854329 | -8363.63564001947 | -8366.01367091872 | 24.550715 | 26.057622 | 26.058566 | 23.074127 |
| **I3b** | -2365.46417775924 | -2366.11100758158 | -8363.66854643547 | -8366.04438976022 | 24.553698 | 26.060557 | 26.061501 | 23.077365 |
| **Step 4 – quenching the radical benzylsuccinate intermediate** | | | | | | | | |
| **I3a** | -2365.47823691481 | -2366.11591250153 | -8363.66773592451 | -8366.04153502962 | 24.550883 | 26.057953 | 26.058897 | 23.074527 |
| **TS4a** | -2365.46024728375 | -2366.09580278392 | -8363.64812122863 | -8366.02071905051 | 24.548263 | 26.054010 | 26.054954 | 23.074894 |
| **I4a** | -2365.49153293516 | -2366.12733865014 | -8363.67945526683 | -8366.05174965579 | 24.554956 | 26.061459 | 26.062403 | 23.079102 |
| **I3$^{\beta}$a** | -2365.48208643381 | -2366.11990094550 | -8363.67353819525 | -8366.04760948655 | 24.550851 | 26.057994 | 26.058938 | 23.073962 |
| **TS4$^{\beta}$a** | -2365.44107367365 | -2366.07751968091 | -8363.63483886080 | -8366.00637384456 | 24.549205 | 26.054500 | 26.055445 | 23.077480 |
| **I4$^{\beta}$a** | -2365.49120157568 | -2366.12646934420 | -8363.68526126910 | -8366.05718887944 | 24.554684 | 26.061274 | 26.062218 | 23.078338 |
| **I3b** | -2365.46598837033 | -2366.11215356271 | -8363.68255468387 | -8366.05785724590 | 24.555896 | 26.062134 | 26.063078 | 23.080288 |
| **TS4b** | -2365.42223170851 | -2366.06841117497 | -8363.63311799287 | -8366.00508215599 | 24.550389 | 26.056410 | 26.057354 | 23.076388 |
| **I4b** | -2365.46598837033 | -2366.13066424488 | -8363.68774125972 | -8366.06110552488 | 24.558061 | 26.064145 | 26.065089 | 23.081065 |

Table S6. Pro*R* pathway: differences in energies calculated for QM2 at B3LYP/6-311g+(2d,2p)/GD3:AMBER level of theory without ($\Delta E^{QM2}$ tzvp ) or with ZPE corrections ($\Delta E^{QM2+ZPE}$ tzvp) calculated with respect to ES geometry; corrections – energy correction introduced to the reaction energy profile matching corresponding stationary points after conformational changes of the model; final energy profile ($\Delta E^{QM2+ZPE}$ tzvp corrected)

| [kJ/mol] | $\Delta E^{QM2}$ tzvp | $\Delta E^{QM2+ZPE}$ tzvp | corrections | $\Delta E^{QM2+ZPE}$ tzvp corrected |
|---|---|---|---|---|
| **Step 1 – activation of Cys493** | | | | |
| **ES** | 0.00 | 0.00 | 0 | 0.00 |
| **ES Hrot** | 6.75 | 3.63 | 0 | 3.63 |
| **TS1** | 62.66 | 59.53 | 0 | 59.53 |
| **I1** | -40.69 | -28.21 | 0 | -28.21 |
| **Step 2 – activation of toluene** | | | | |
| **I1a** | -32.16 | -10.04 | -18.17 | -28.21 |
| **TS2a** | 62.82 | 67.89 | -18.17 | 49.72 |
| **I2a** | -2.25 | 10.50 | -18.17 | -7.67 |
| **I1b** | -1.28 | 23.04 | -18.17 | 4.87 |
| **TS2b** | 92.69 | 97.52 | -18.17 | 79.34 |
| **I2b** | 33.51 | 45.83 | -18.17 | 27.66 |
| **I1c** | -18.91 | 5.00 | -18.17 | -13.17 |
| **TS2** | 42.41 | 52.58 | -18.17 | 34.41 |
| **I2c** | 14.25 | 31.40 | -18.17 | 13.23 |
| **Step 3 - C-C bond formation** | | | | |
| **I2c** | 213.92 | 243.44 | -230.21 | 13.23 |
| **TS3** | 261.66 | 294.42 | -230.21 | 64.21 |
| **I3** | 171.27 | 214.83 | -230.21 | -15.38 |
| **I2b** | 208.95 | 237.94 | -224.71 | 13.23 |
| **TS3b** | 299.78 | 335.17 | -224.71 | 110.45 |
| **I3b** | 223.00 | 264.18 | -224.71 | 39.46 |
| **Step 4 – quenching the radical benzylsuccinate intermediate** | | | | |
| **$I3^{\alpha}a$** | 96.46 | 138.44 | -153.82 | -15.38 |
| **$TS4^{\alpha}a$** | 163.07 | 198.91 | -153.82 | 45.09 |
| **$I4^{\alpha}a$** | 69.25 | 124.84 | -153.82 | -28.97 |
| **$I3^{\beta}a$** | 37.93 | 88.15 | -103.53 | -15.38 |
| **$TS4^{\beta}a$** | 161.35 | 194.46 | -103.53 | 90.93 |
| **I4a** | 5.43 | 65.46 | -103.53 | -38.07 |
| **I3b** | 223.00 | 264.18 | -224.71 | 39.46 |
| **TS4b** | 285.07 | 318.12 | -224.71 | 93.41 |
| **I4b** | 175.88 | 221.80 | -224.71 | -2.91 |
| **Step 5 – transfer of the H atom between Gly and Cys for the E:P complex.** | | | | |
| **I4_rot** | 75.40 | 130.94 | -153.82 | -22.87 |
| **I4** | 81.29 | 139.88 | -162.75 | -22.87 |
| **TS5** | 195.79 | 238.00 | -162.75 | 75.25 |
| **EP** | 131.38 | 180.55 | -162.75 | 17.80 |

Table S4. Pro*S* pathway: differences in energies calculated for QM2 at B3LYP/6-311g+(2d,2p)/GD3:AMBER level of theory without ($\Delta E^{QM2}$ tzvp ) or with ZPE corrections ($\Delta E^{QM2+ZPE}$ tzvp) calculated with respect to I2b geometry; corrections – energy correction introduced to the reaction energy profile matching corresponding stationary points after conformational changes of the model; final energy profile ($\Delta E^{QM2+ZPE}$ tzvp corrected); I3$^{\alpha}$a' and I3$^{\beta}$a' represent respective I3 structures with shifter proton.

| [kJ/mol] | $\Delta E^{QM2}$ tzvp | $\Delta E^{QM2+ZPE}$ tzvp | corrections | $\Delta E^{QM2+ZPE}$ tzvp corrected |
|---|---|---|---|---|
| **Step 1 – activation of Cys493** | | | | |
| **I2c** | 14.25 | 31.40 | -18.17 | 13.23 |
| **Step 3 - C-C bond formation** | | | | |
| **I2a** | 4.35 | 7.61 | 5.62 | 13.23 |
| **TS3a** | 29.65 | 39.14 | 5.62 | 44.76 |
| **I3a** | -38.80 | -26.87 | 5.62 | -21.25 |
| **I2b** | 0.00 | 0.00 | 13.23 | 13.23 |
| **TS3b** | 74.56 | 79.66 | 13.23 | 92.89 |
| **I3b** | -41.45 | -22.75 | 13.23 | -9.52 |
| **Step 4 – quenching the radical benzylsuccinate intermediate** | | | | |
| **I3$^{\alpha}$a'** | 1.40 | 6.94 | 5.62 | 12.56 |
| **TS4 $^{\alpha}$a** | 56.06 | 54.71 | 5.62 | 60.33 |
| **I4$^{\alpha}$a** | -25.42 | -9.18 | 5.62 | -3.56 |
| **I3$^{\beta}$a'** | -14.55 | -9.09 | 5.62 | -3.47 |
| **TS4 $^{\beta}$a** | 93.72 | 94.85 | 5.62 | 100.47 |
| **I4$^{\beta}$a** | -39.70 | -24.18 | 5.62 | -18.56 |
| **I3b** | -41.45 | -22.75 | 13.23 | -9.52 |
| **TS4b** | 97.11 | 101.35 | 13.23 | 114.58 |
| **I4b** | -49.98 | -25.60 | 13.23 | -12.37 |

#### Kinetic isotope effect

Table S7. Vibrational corrections calculated for toluene and $d_8$-toluene at 303 K, 1 atm and a scaling factor of 0.9806 for pro*R* pathway.

| | ZPE | Thermal | H | G |
|---|---|---|---|---|
| **ES** | 24.060562 | 25.647039 | 25.647999 | 22.489731 |
| **ES $d_8$** | 24.034850 | 25.622422 | 25.623382 | 22.462949 |
| **TS1** | 24.059419 | 25.644356 | 25.645315 | 22.490913 |
| **TS1 $d_8$** | 24.033692 | 25.619726 | 25.620686 | 22.464119 |
| **I1** | 24.065249 | 25.651198 | 25.652157 | 22.494700 |
| **I1 $d_8$** | 24.039525 | 25.626570 | 25.627529 | 22.467910 |
| **I1c** | 24.069516 | 25.652960 | 25.653919 | 22.504934 |
| **I1c $d_8$** | 24.043781 | 25.628323 | 25.629283 | 22.478131 |
| **TS2c** | 24.064383 | 25.647535 | 25.648494 | 22.500633 |
| **TS2c $d_8$** | 24.040425 | 25.624824 | 25.625784 | 22.475778 |
| **I2c** | 24.066995 | 25.651131 | 25.652090 | 22.500720 |
| **I2c $d_8$** | 24.042313 | 25.627749 | 25.628709 | 22.474893 |
| **I2c** | 24.071611 | 25.653174 | 25.654133 | 22.513249 |
| **I2c $d_8$** | 24.046967 | 25.629833 | 25.630793 | 22.487452 |
| **TS3** | 24.072820 | 25.653069 | 25.654028 | 22.516258 |
| **TS3 $d_8$** | 24.047858 | 25.629374 | 25.630333 | 22.490216 |
| **I3** | 24.076855 | 25.655490 | 25.656450 | 22.524540 |
| **I3 $d_8$** | 24.051479 | 25.631310 | 25.632269 | 22.498184 |
| **I3a** | 24.076265 | 25.655491 | 25.656451 | 22.521895 |
| **I3a $d_8$** | 24.050888 | 25.631305 | 25.632264 | 22.495519 |
| **TS4a** | 24.073973 | 25.651857 | 25.652816 | 22.522758 |
| **TS4a $d_8$** | 24.049555 | 25.628623 | 25.629582 | 22.497554 |
| **I4** | 24.081349 | 25.659483 | 25.660443 | 22.528735 |
| **I4 $d_8$** | 24.055046 | 25.634321 | 25.635280 | 22.501628 |
| **I4** | 24.082470 | 25.660248 | 25.661208 | 22.531462 |
| **I4 $d_8$** | 24.056155 | 25.635076 | 25.636035 | 22.504331 |
| **TS5** | 24.076352 | 25.653280 | 25.654239 | 22.527184 |
| **TS5 $d_8$** | 24.050029 | 25.628098 | 25.629058 | 22.500049 |
| **EP** | 24.078949 | 25.656978 | 25.657937 | 22.527532 |
| **EP $d_8$** | 24.052637 | 25.631807 | 25.632767 | 22.500403 |

Table S8. Kinetic constants and intrinsic kinetic isotope effects (iKIE) calculated for each step of the proR reaction pathway for the highest level of theory and thermal energy corrections.

| | **Δ($E^{QM2}$ tzvp+Therm) [kJ/mol]** | | **k [$s^{-1}$]** | **iKIE** | **Δ($E^{QM2}$ tzvp+Therm) [kJ/mol]** | | **k [$s^{-1}$]** | **iKIE** |
|---|---|---|---|---|---|---|---|---|
| | forward | | | | reverse | | | |
| **ES** | 0.00 | | | | 29.78 | | | |
| **ES $d_8$** | 0.00 | | | | 29.80 | | | |
| **TS1** | 55.61 | $k_2$ | 1635 | **0.99** | 85.39 | $k_{-2}$ | 1.20.E-02 | **1.0** |
| **TS1 $d_8$** | 55.58 | $^Dk_2$ | 1657 | | 85.38 | $^Dk_{-2}$ | 1.21.E-02 | |
| **I1** | -29.78 | | | | 0.00 | | | |
| **I1 $d_8$** | -29.80 | | | | 0.00 | | | |
| **I1c** | 0.00 | | | | -28.35 | | | |
| **I1c $d_8$** | 0.00 | | | | -31.65 | | | |
| **TS2c** | 47.08 | $k_3$ | 48295 | **7.44** | 18.73 | $k_{-3}$ | $3.73*10^9$ | **2.0** |
| **TS2c $d_8$** | 52.14 | $^Dk_3$ | 6489 | | 20.49 | $^Dk_{-3}$ | $1.85*10^9$ | |
| **I2c** | 28.35 | | | | 0.00 | | | |
| **I2c $d_8$** | 31.65 | | | | 0.00 | | | |
| **I2c** | 0.00 | | | | 36.57 | | | |
| **I2c $d_8$** | 0.00 | | | | 38.77 | | | |
| **TS3** | 47.47 | $k_4$ | 41474 | **0.69** | 84.04 | $k_{-4}$ | $2.06*10^{-2}$ | **1.7** |
| **TS3 $d_8$** | 46.54 | $^Dk_4$ | 59979 | | 85.31 | $^Dk_{-4}$ | $1.24*10^{-2}$ | |
| **I3** | -36.57 | | | | 0.00 | | | |
| **I3 $d_8$** | -38.77 | | | | 0.00 | | | |
| **I3a** | 0.00 | | | | 4.68 | | | |
| **I3a d8** | 0.00 | | | | 7.25 | | | |
| **TS4a** | 57.07 | $k_5$ | 917 | **2.70** | 61.75 | $k_{-5}$ | 143 | **7.5** |
| **TS4a $d_8$** | 59.57 | $^Dk_5$ | 399 | | 66.81 | $^Dk_{-5}$ | 19 | |
| **I4** | -4.68 | | | | 0.00 | | | |
| **I4d8** | -7.25 | | | | 0.00 | | | |
| **I4** | 0.00 | | | | -41.51 | | | |
| **I4 $d_8$** | 0.00 | | | | -41.51 | | | |
| **TS5** | 96.20 | $k_6$ | $1.64*10^{-4}$ | **0.99** | 54.70 | $k_{-6}$ | 2351 | 1.0 |
| **TS5 $d_8$** | 96.18 | $^Dk_6$ | $1.66*10^{-4}$ | | 54.67 | $^Dk_{-6}$ | 2378 | |
| **EP** | 41.51 | | | | 0.00 | | | |
| **EP $d_8$** | 41.51 | | | | 0.00 | | | |

Table S9. Vibrational corrections calculated for toluene at 303 K, 1 atm and a scaling factor of 0.9806 for pro*S* pathway.

| | ZPE | Thermal | H | G |
|---|---|---|---|---|
| **I2a** | 24.073746 | 25.654890 | 25.655850 | 22.514251 |
| **TS3a** | 24.076102 | 25.655239 | 25.656198 | 22.519453 |
| **I3a** | 24.076983 | 25.656723 | 25.657683 | 22.520527 |
| **I3a'** | 24.074596 | 25.654234 | 25.655194 | 22.518692 |
| **TS4a** | 24.072026 | 25.650334 | 25.651293 | 22.519206 |
| **I4** | 24.078590 | 25.657638 | 25.658597 | 22.523217 |

Table S10. Kinetic constants calculated for each step of the pro*S* reaction pathway for the highest level of theory and thermal energy corrections.

| | Δ($E^{QM2}$ tzvp+Therm) [kJ/mol] | | k [$s^{-1}$] | Δ($E^{QM2}$ tzvp+Therm) [kJ/mol] | | k [$s^{-1}$] |
|---|---|---|---|---|---|---|
| **I2c** | 0.00 | | | 38.34 | | |
| **TS3** | 26.22 | $k_4$ | $1.9*10^8$ | 64.56 | $k_{-4}$ | 46.9 |
| **I3a** | -38.34 | | | 0.00 | | |
| **I3a** | 0.00 | | | -15.79 | | |
| **TS4a** | 78.1 | $k_5$ | 0.22 | 62.29 | $k_{-5}$ | 143 |
| **I4** | 15.8 | | | 0.00 | | |

#### H/D Exchange

Table S11. Kinetic constants and intrinsic kinetic isotope effects (iKIE) calculated for H/D exchange of the *R*-benzylsuccinate for the highest level of theory and thermal energy corrections.

| | Δ($E^{QM2}$ tzvp+Therm) [kJ/mol] | | k [$s^{-1}$] | iKIE | Δ($E^{QM2}$ tzvp+Therm) [kJ/mol] | | k [$s^{-1}$] | |
|---|---|---|---|---|---|---|---|---|
| | forward | | | | reverse | | | |
| **I3a** | 0.00 | | | | 5.66 | | | |
| **$^{\alpha}d_1$-I3a** | 0.00 | | | | 8.30 | | | |
| **$^{\beta}d_1$-I3a** | 0.00 | | | | 5.19 | | | |
| **$d_2$-I3a** | 0.00 | | | | 7.92 | | | |
| **$TS4^{\alpha}$** | 57.1 | $^{\alpha}k_5$ | 917 | 1.0 | 62.73 | $^{\alpha}k_{-5}$ | 97 | 1.0 |
| **$^{\alpha}d_1$-$TS4^{\alpha}$** | 59.7 | $^{\alpha D}k_5$ | 326 | 2.8 | 67.97 | $^{\alpha D}k_{-5}$ | 12 | 8.0 |
| **$^{\beta}d_1$-$TS4^{\alpha}$** | 62.5 | $^{sec\text{-}\alpha}k_5$ | 104 | 8.8 | 61.91 | $^{sec\text{-}\alpha}k_{-5}$ | 134 | 0.7 |
| **$d_2$-$TS4^{\alpha}$** | 59.3 | $^{D2\alpha}k_5$ | 375 | 2.4 | 67.24 | $^{D2\alpha}k_{-5}$ | 16 | 6.0 |
| **I4a** | -5.7 | | | | 0.00 | | | |
| **$^{\alpha}d_1$-I4a** | -8.3 | | | | 0.00 | | | |
| **$^{\beta}d_1$-I4a** | -5.2 | | | | 0.00 | | | |
| **$d_2$-I4a** | -7.9 | | | | 0.00 | | | |

| | | | | | | | | |
|---|---|---|---|---|---|---|---|---|
| **I3a** | 0.00 | | | | 26.11 | | | |
| **$^{\alpha}d_1$-I3a** | 0.00 | | | | 32.84 | | | |
| **$^{\beta}d_1$-I3a** | 0.00 | | | | 18.83 | | | |
| **$d_2$-I3a** | 0.00 | | | | 132.20 | | | |
| **TS4$^{\beta}$** | 108.4 | $^{\beta}k_5$ | $1.3*10^{-6}$ | 1.0 | 137.12 | $^{\beta}k_{-5}$ | $1.0*10^{-10}$ | 1.0 |
| **$^{\beta}d_1$-TS4$^{\beta}$** | 111.0 | $^{\beta D}k_5$ | $4.6*10^{-7}$ | 2.8 | 140.74 | $^{\beta D}k_{-5}$ | $1.5*10^{-11}$ | 7.1 |
| **$^{\alpha}d_1$-TS4$^{\beta}$** | 107.9 | $^{sec\text{-}\beta}k_5$ | $1.6*10^{-6}$ | 0.8 | 129.35 | $^{sec\text{-}\beta}k_{-5}$ | $3.5*10^{-12}$ | 29.7 |
| **$d_2$-TS4$^{\beta}$** | 110.5 | $^{D2\beta}k_5$ | $6.E*10^{-7}$ | 2.3 | 0.00 | $^{D2\beta}k_{-5}$ | $3.2*10^{-10}$ | 0.3 |
| **I4a** | -23.8 | | | | 0.00 | | | |
| **$^{\alpha}d_1$-I4a** | -26.1 | | | | 0.00 | | | |
| **$^{\beta}d_1$-I4a** | -32.8 | | | | 0.00 | | | |
| **$d_2$-I4a** | -18.8 | | | | 26.11 | | | |
| **$k_{H/D}$** | $k_{HDX}$*[$D_2O$]= 0.01565 $s^{-1}$ $M^{-1}$*22.22 M= 0.35 $s^{-1}$ | | | | | | | |
| **$k_{D/H}$** | $k_{HDX}$*[$H_2O$]=0.01565 $s^{-1}$ $M^{-1}$ *33.33 M= 0.52 $s^{-1}$ | | | | | | | |

#### Acknowledgements

The authors acknowledge the financial support provided by Deutsche Forschungsgemeinschaft/National Science Center Poland under Beethoven Life grant He2190/13-1 / 2018/31/F/NZ1/01856 as well as Polish high-performance computing infrastructure PLGrid (HPC Center: ACK Cyfronet AGH) for providing computer facilities and support within computational grant no. PLG/2023/016888, PLG/2022/016024 and PLG/2021/015218.
